# Amplification of avian influenza viruses along poultry marketing chains in Bangladesh: a controlled field experiment

**DOI:** 10.1101/2023.11.10.566573

**Authors:** Lisa Kohnle, Tridip Das, Md. Helal Uddin, Sanjib C. Nath, Md. Abu Shoieb Mohsin, Rashed Mahmud, Paritosh K. Biswas, Md. Ahasanul Hoque, Dirk U. Pfeiffer, Guillaume Fournié

**Affiliations:** City University of Hong Kong, China; Chattogram Veterinary and Animal Sciences University, Bangladesh; Charles Sturt University, Wagga Wagga, Australia; Royal Veterinary College, London, United Kingdom; Université de Lyon, INRAE, VetAgro Sup, UMR EPIA, Marcy l’Etoile, France; Université Clermont Auvergne, INRAE, VetAgro Sup, UMR EPIA, Saint Genes Champanelle, France

**Keywords:** avian influenza, influenza A, H9N2, H5N1, zoonotic disease, poultry, live bird market, network, value chain, veterinary epidemiology, One Health

## Abstract

The prevalence of avian influenza viruses (AIVs) is commonly found to increase dramatically from farms to live bird markets (LBMs). Viral transmission dynamics along marketing chains is, however, poorly understood. To address this gap, we implemented a field experiment altering chicken supply to an LBM in Chattogram, Bangladesh. Chickens traded along altered (intervention) and conventional (control) marketing chains were tested for AIVs. Upon arrival at the LBM, the odds of detecting AIVs did not differ between control and intervention groups. However, 12 hours later, intervention group odds were lower, particularly for broiler chickens, indicating that viral shedding in LBM resulted partly from infections during transport and trade. Curtailing AIV prevalence in LBMs requires mitigating risk in marketing chain nodes preceding chickens’ delivery at LBMs.

**Article Summary Line:** The high prevalence of avian influenza viruses in marketed chickens cannot be solely attributed to viral transmission within live bird markets but is also influenced by infections occurring prior to the chickens’ supply to these markets.

## Background

Multiple subtypes of avian influenza viruses (AIVs) are endemic in poultry populations throughout Asia. In Bangladesh, the H9N2 and H5N1 subtypes are predominant, threatening both commercial poultry production and the livelihoods of small-scale poultry farmers.^1–3^ Moreover, the co-circulation of both subtypes and their potential reassortment raise concerns about the emergence of new variants with pandemic potential.^4^

Chickens, the main source of animal protein in the country, are raised in diverse production systems and traded along complex marketing chains.^5,6^ The latter often involve multiple stakeholders (e.g., mobile traders) and therefore make transport and trade processes difficult to trace. Common risks include insufficient cleaning of transport vehicles, mixing of poultry of different origins or species, and prolonged transport durations (ranging from several hours to several days, depending on the production system and origin as well as the number of live bird markets (LBMs) visited).^5,6^ A marketing chain is hereinafter defined as all steps (transport, storage and transactions) taking place between poultry production sites and points of sale to consumers.

LBMs, where poultry from different origins and species are mixed,^5^ slaughtered and sold to consumers, are often referred to as viral reservoirs.^7–11^ AIVs are ubiquitous and their prevalence in poultry is typically high.^12–15^ Indeed, the estimated prevalence of influenza A(H9) (A(H5)) viruses in chickens marketed in Dhaka and Chattogram cities ranged between 6.8%– 13.1% (0.9%–1.3%), depending on chicken and LBM type.^13^ In contrast, influenza A(H9) (A(H5)) viruses were found in <0.5% (none) of farmed chickens in a cross-sectional study conducted in areas supplying Chattogram city’s LBMs,^16^ indicating a more than 10-fold increase in AIV prevalence along poultry marketing chains in Bangladesh.

Once introduced, poultry remain in LBMs for only a short period of time. In Chattogram city, the probability of a chicken spending >24 (48) hours in a market stall was estimated at 0.14 (0.03).^5^ Depending on the length of the latent period,^17^ most chickens may not spend enough time in LBMs to become infected with and start shedding AIVs. Therefore, the high AIV prevalence observed in marketed chickens may not only result from transmission occurring within LBMs but also at upstream stages of marketing chains. We hypothesise that chickens shedding AIVs in market stalls may have already been exposed at the farm gate or during transport before reaching LBMs at an advanced stage of infection.

We aimed to explore the extent to which transport and trade practices contribute to the increase in AIV prevalence from farms to LBMs. By implementing a controlled field experiment through which marketing chains were altered, we assessed whether reducing the risk of infection for chickens along all stages reduced viral shedding in LBMs.

## Methods

Chickens traded along altered (intervention) and conventional (control) marketing chains were tested for AIVs by screening oropharyngeal swab samples collected at different stages between farms and market stalls. The effect of the intervention in reducing bird-level prevalence in LBMs was assessed by comparing the proportions of positive chickens between intervention and control groups.

This study took place in south-eastern Bangladesh between March–August 2019 (Appendix A). Study locations comprised Chattogram Veterinary and Animal Sciences University (CVASU), production sites in Chattogram and neighbouring districts, and LBMs in Chattogram city. We targeted chickens reared for meat production, specifically broilers, raised in commercial farms, and indigenous backyard chickens, raised for meat and egg production in rural households. The field experiment was repeated for 64 batches (30 and 34 for broilers and backyard chickens, respectively; sample size estimation in Appendix B) of 10 chickens and comprised two successive parts (Figure 1, Table 1).

**Figure 1.**
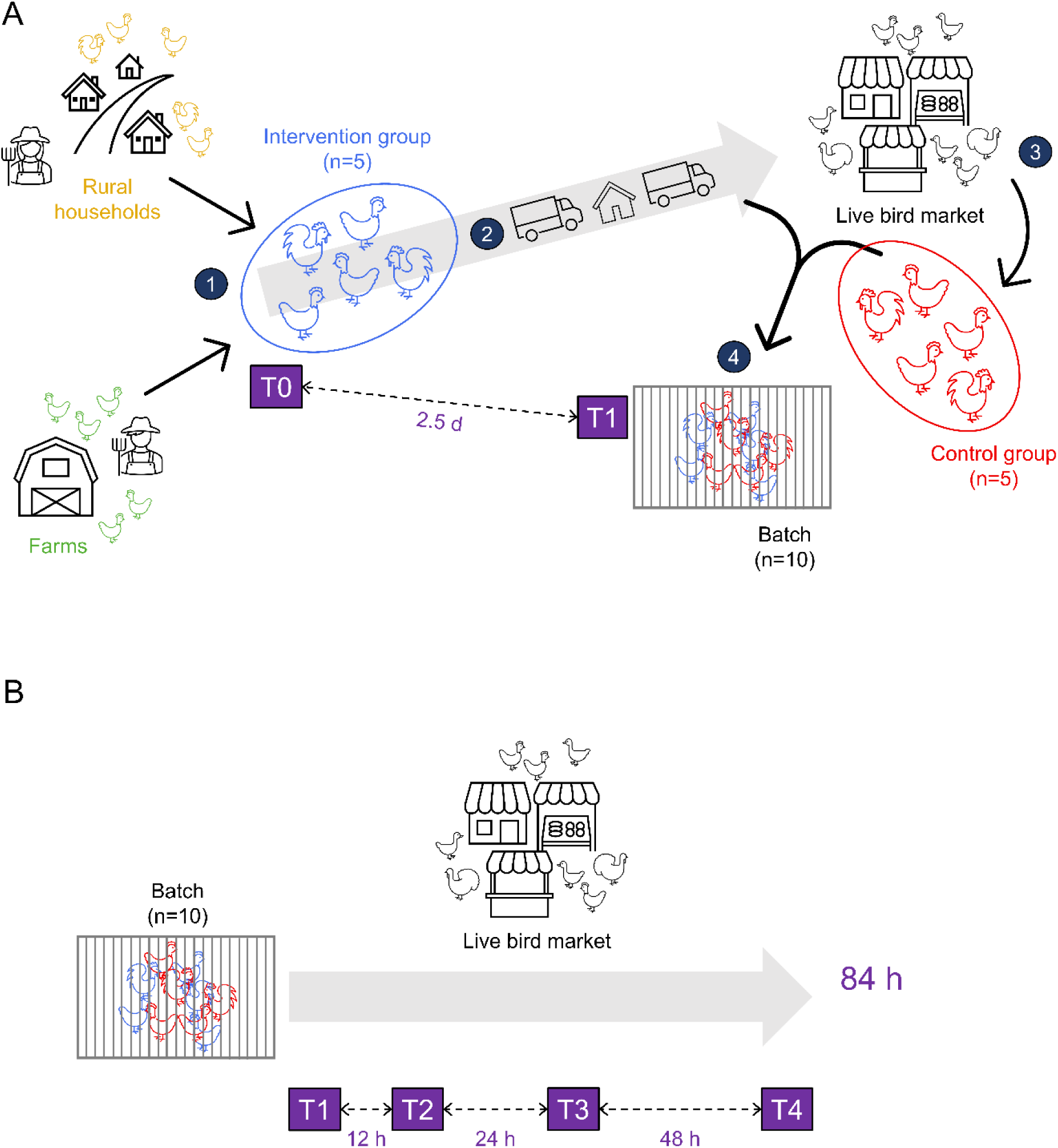
Structure and components of the field experiment. A) first part: (1) recruitment of intervention groups pre-tests (T0), (2) transport and storage of intervention groups, (3) recruitment of control groups, (4) post-tests (matching of intervention and control groups; B) second part: longitudinal sampling in market stalls.

**Table 1.**
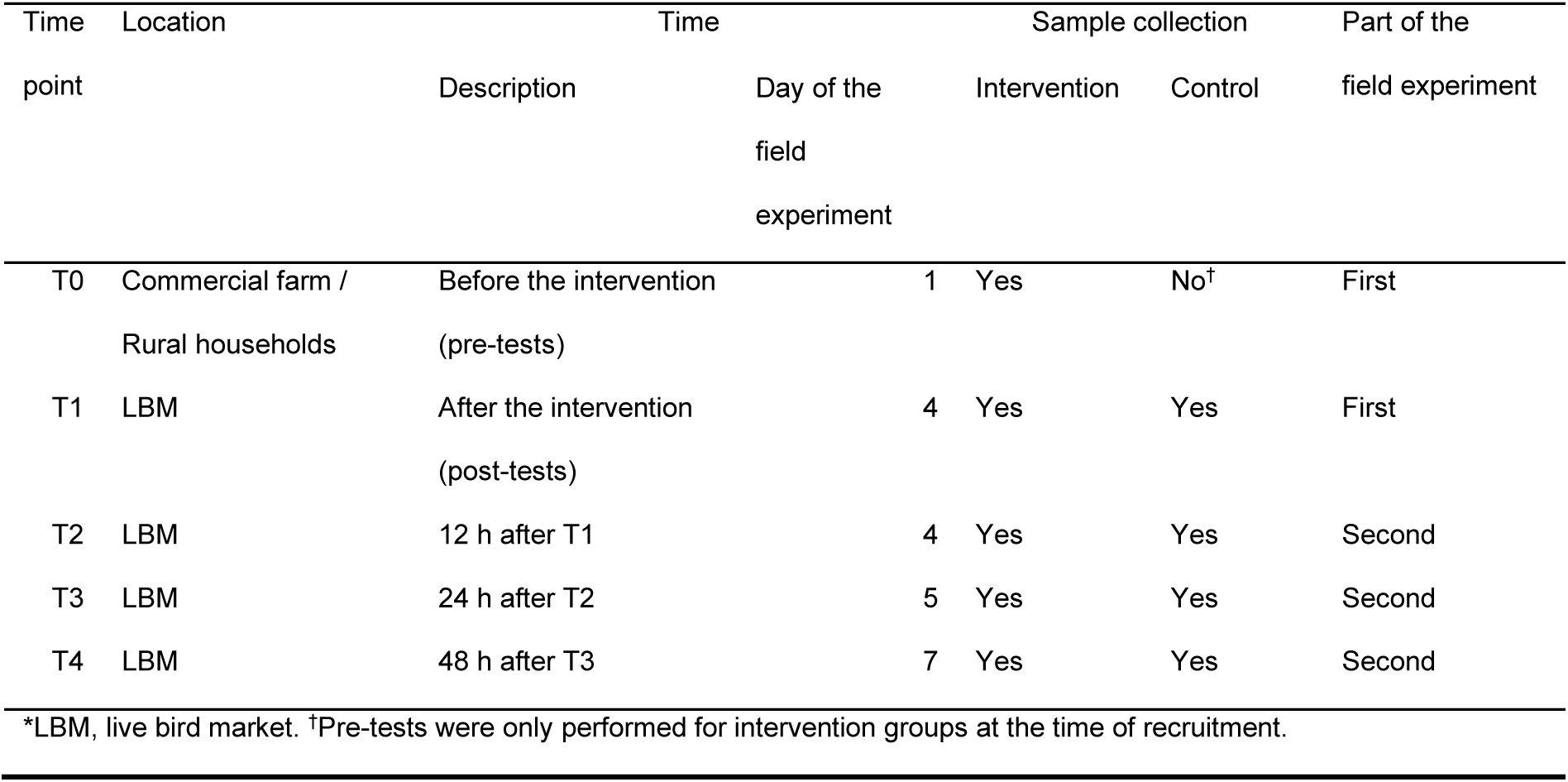
Characteristics of samples collected at different time points*.

### Field experiment

In the first part (Figure 1A, Table 1), for each batch, 5 chickens were purchased from either a commercial farm (broilers) or 5 rural households (backyard chickens) in a village. Those chickens were sampled upon purchase (T0, pre-tests) and exposed to the intervention. The practice of selling batches to multiple stakeholders over several days and their frequent visits in the last days of a production cycle likely increase the risk of viral incursion. We therefore aimed to select commercial farms with ≥2–3 days left on their current production cycle to minimise the risk of purchasing broilers already infected with AIVs. Farms and villages were recruited as follows. For each farm, a sub-district (‘upazila’) was randomly selected with a probability proportional to the estimated number of chickens it supplied to Chattogram city’s LBMs (Appendix C).^5^ In the absence of census data, eligible farms were identified through feed dealers operating in the selected sub-districts. Feed dealers, who provided farmers with production inputs and credits, were randomly selected from lists compiled by consulting other relevant stakeholders. The origins of backyard chickens supplied to Chattogram city’s LBMs are geographically diverse and only a fraction could be covered (Chattogram, Cox’s Bazar and Feni districts, accounting for 17.1% of the poultry flow). For villages, as census data were not available, we used QGIS 3.0.0-Girona to generate as many random coordinates within the selected study area as the number of villages to be recruited. The nearest village to each generated point was included. Within each village, we then selected 5 rural households willing to sell backyard chickens. Inclusion criteria for recruited chickens are described in Appendix C.

The intervention consisted in the rigorous implementation of standardised biosecurity measures during transport, storage and delivery of chickens to Pahartali Kacha Bazar, our study LBM (altered marketing chain, Appendix D). In brief, our research team acted as mobile traders, transporting intervention groups, one at a time, on our own vehicle to CVASU, where they were stored in a purpose-built poultry shed for 2.5 d. This was meant to reflect the extended transport durations observed along certain marketing chains and to assess whether intervention groups testing negative at the farm gate might have been latently infected. Keeping chickens from different batches always separate, thoroughly cleaning and disinfecting all surfaces in contact with chickens reduced risks of contamination between intervention groups and from the environment. After 2.5 d, we transferred our intervention groups to the LBM always before 7 AM, when most mobile traders supply poultry. Upon arrival, 5 chickens, each forming a control group, were purchased from a mobile trader. If none was found, 5 chickens were purchased from a stallholder reportedly supplied only shortly before (median=1.4 h). For backyard chickens, control groups were purchased in a different LBM, from which stallholders in the study LBM were supplied. Intervention and control groups were sampled (T1, post-tests), tagged with leg bands and caged together in one of two market stalls, selected according to stallholders’ willingness to participate and current capacity.

In the second part (Figure 1B), each batch of 10 chickens was kept in the same cage for 84 hours and sampled longitudinally (Table 1). Study chickens were exposed to the same market stall environment and practices as other poultry offered for sale. If a chicken died between two time points, a last sample was collected at the latter.

Ethical approval was obtained from both City University of Hong Kong and Chattogram Veterinary and Animal Sciences University.

### Sample collection

We collected oropharyngeal swabs from chickens at different time points (Table 1, Appendix E). They were placed in individual tubes containing viral transport medium, stored in a cool box during transport and frozen at –80°C in the laboratory.

### Laboratory analysis

We extracted viral RNA with MagMAX™-96 Viral RNA Isolation Kits (Thermo Fisher Scientific, Applied Biosystems^TM^) and performed real-time reverse transcription PCR (rRT-PCR) with AgPath-ID^TM^ One-Step RT-PCR Reagents (Thermo Fisher Scientific, Applied Biosystems^TM^) to screen for the matrix (M), H9 haemagglutinin (HA) and H5 HA genes (Appendix F) according to the protocols of the Australian Centre for Disease Preparedness (ACDP, formerly known as the Australian Animal Health Laboratory) at the Commonwealth Scientific and Industrial Research Organisation (CSIRO, Australia).^18^ A sample was considered positive if the cycle threshold (C_t_) value was <40. We also considered an alternative positivity threshold, C_t_<33, to minimise concerns about a positive sample with high C_t_ value having resulted from contamination of the bird’s oropharynx as opposed to actual viral shedding following infection. Testing algorithms are further described in Appendix F. A subset of 152 samples were sequenced, with all H9- and H5-positive samples being related to H9N2 (n=129), H5N1 (n=17), or both (n=6). Complete test results, including C_t_ values, are available in Appendix G.

### Data analysis

For each gene and positivity threshold, we computed the cumulative number and proportion of chickens positive at each time point (T0–T4) with respect to group (intervention and control) and chicken type (broilers and backyard chickens) (Table 2). An intervention or control group is hereinafter defined as a sub-batch and broilers and backyard chickens as strata. Time points were clustered within chickens and chickens were clustered within sub-batches (up to T1) or batches (after T1).

**Table 2.**
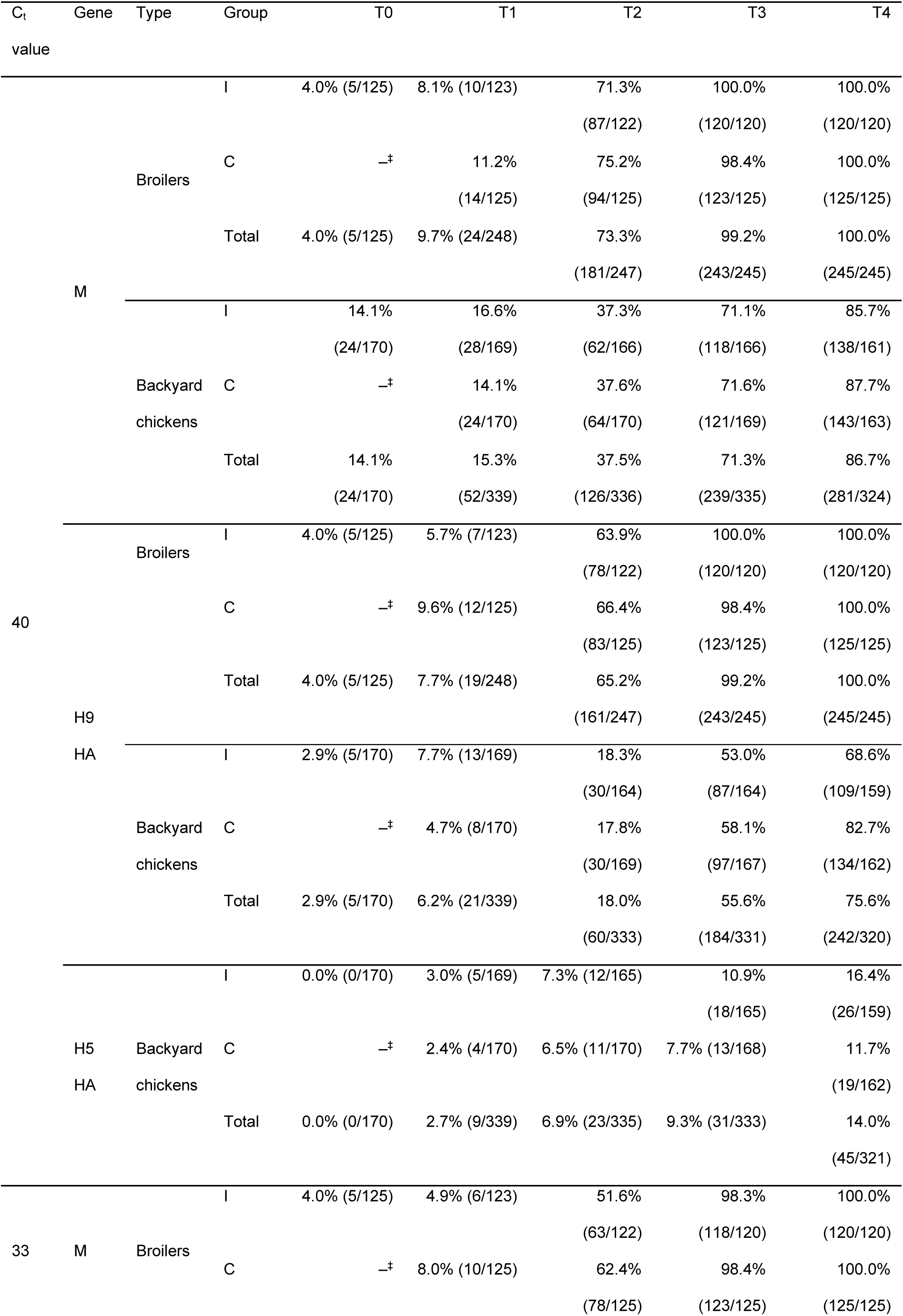
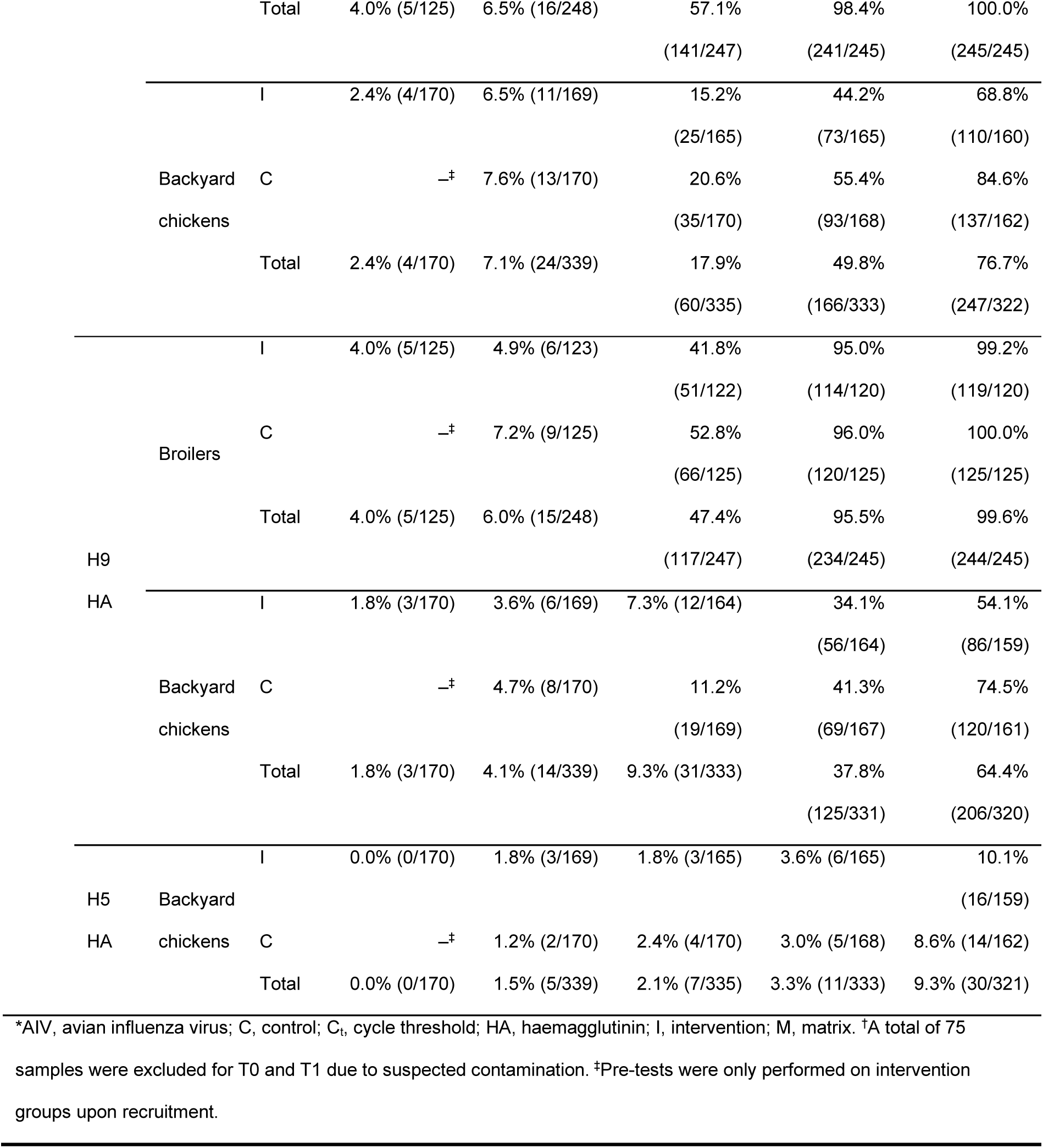
Cumulative incidence of AIVs in chickens at different time points^*†^.

To assess whether the intervention reduced the proportion of positive chickens entering the LBM (T1), we implemented, for each combination of gene, positivity threshold and stratum, a logistic regression model with random intercepts for sub-batches. The binary test status of chickens and group were used as response and explanatory variables, respectively. Confidence intervals were computed with the Wald method.

We investigated the effect of the intervention on the proportion of chickens in negative batches (at T1) becoming positive by all following time points (T2–T4) by applying conditional logistic regression models with batches as matched sets, for each combination of gene, positivity threshold and stratum.

Models were implemented in R v4.1.2,^19^ using the lme4^20^ and survival^21,22^ packages.

## Results

Of the 640 chickens recruited, 43 dropped out of the study: 41 died suddenly (Appendix Figure 10) and two were slaughtered for welfare reasons. Of those 43 chickens, 30, 21 and 11 had previously tested positive for the M, H9 HA and H5 HA genes, respectively. Seventy-five T0 and/or T1 samples from 8 successive broiler batches were discarded due to suspected sample contamination (Appendix F).

Our results indicate that only few chickens entered the LBM already infected or contaminated with AIVs (12.9% and 6.8% for C_t_<40 and C_t_<33, respectively), whereas cumulative incidence increased substantially over time, with most chickens testing positive for the M (92.4% and 86.8% for C_t_<40 and C_t_<33, respectively) and H9 HA genes (86.2% and 79.6% for C_t_<40 and C_t_<33, respectively) by T4 (after 84 h) (Table 2, Figure 2). The number of positive chickens in a batch at T1 ranged between 1–7 (1–6), for C_t_<40 (C_t_<33) (Figure 3), and a large proportion of AIV-positive chickens were also H9-positive. By T4, influenza A(H5) viruses were only detected in 45 (14%) backyard chickens from 14 batches, but not in broilers (Table 2, Figure 2).

**Figure 2.**
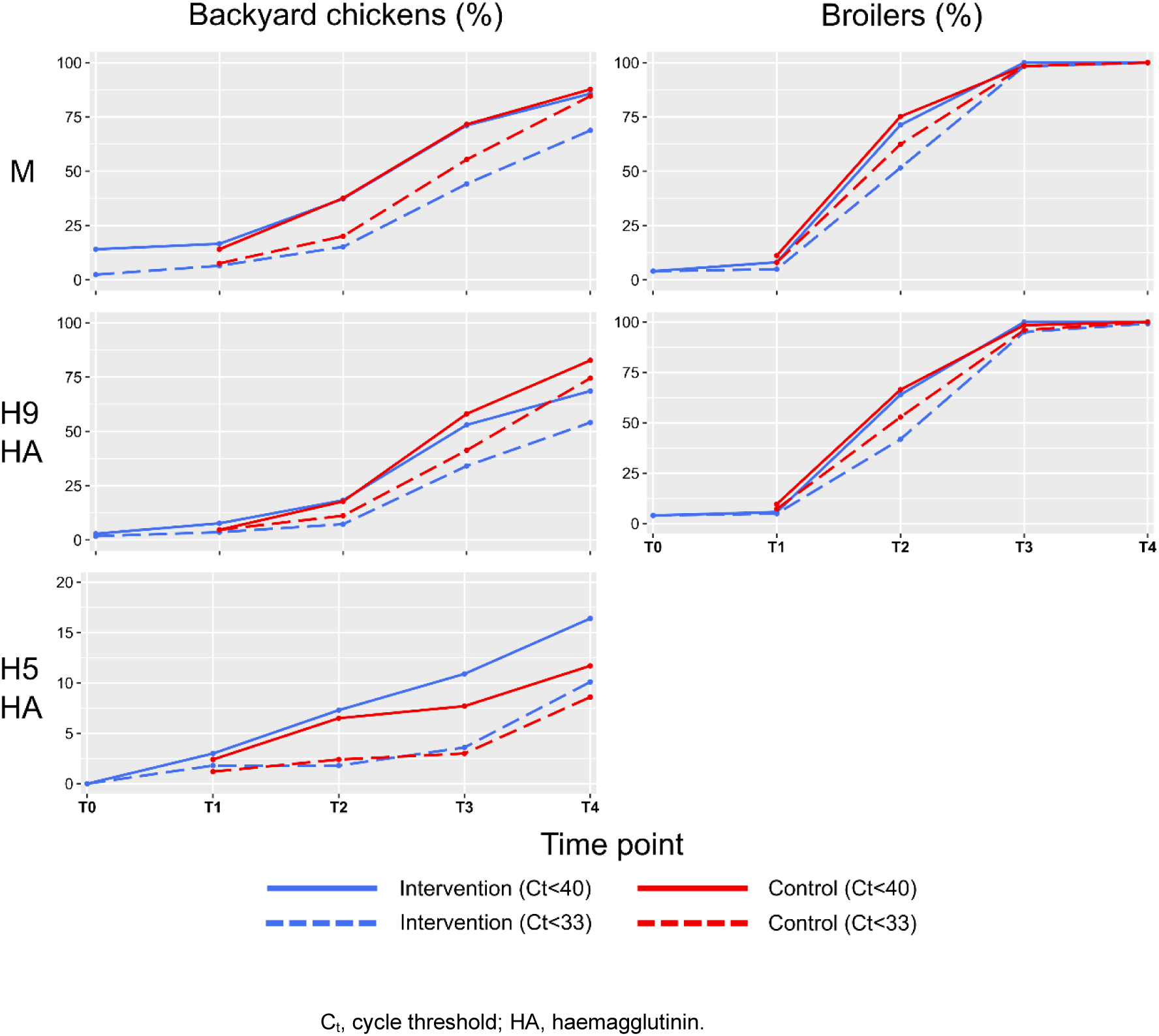
Temporal distribution of the cumulative proportion of positive chickens (M, H9 HA and H5 HA genes) for intervention and control groups, considering two different positivity thresholds (the y-axis scale is different for the H5 HA gene compared to the other genes due to the smaller number of positive samples)

**Figure 3.**
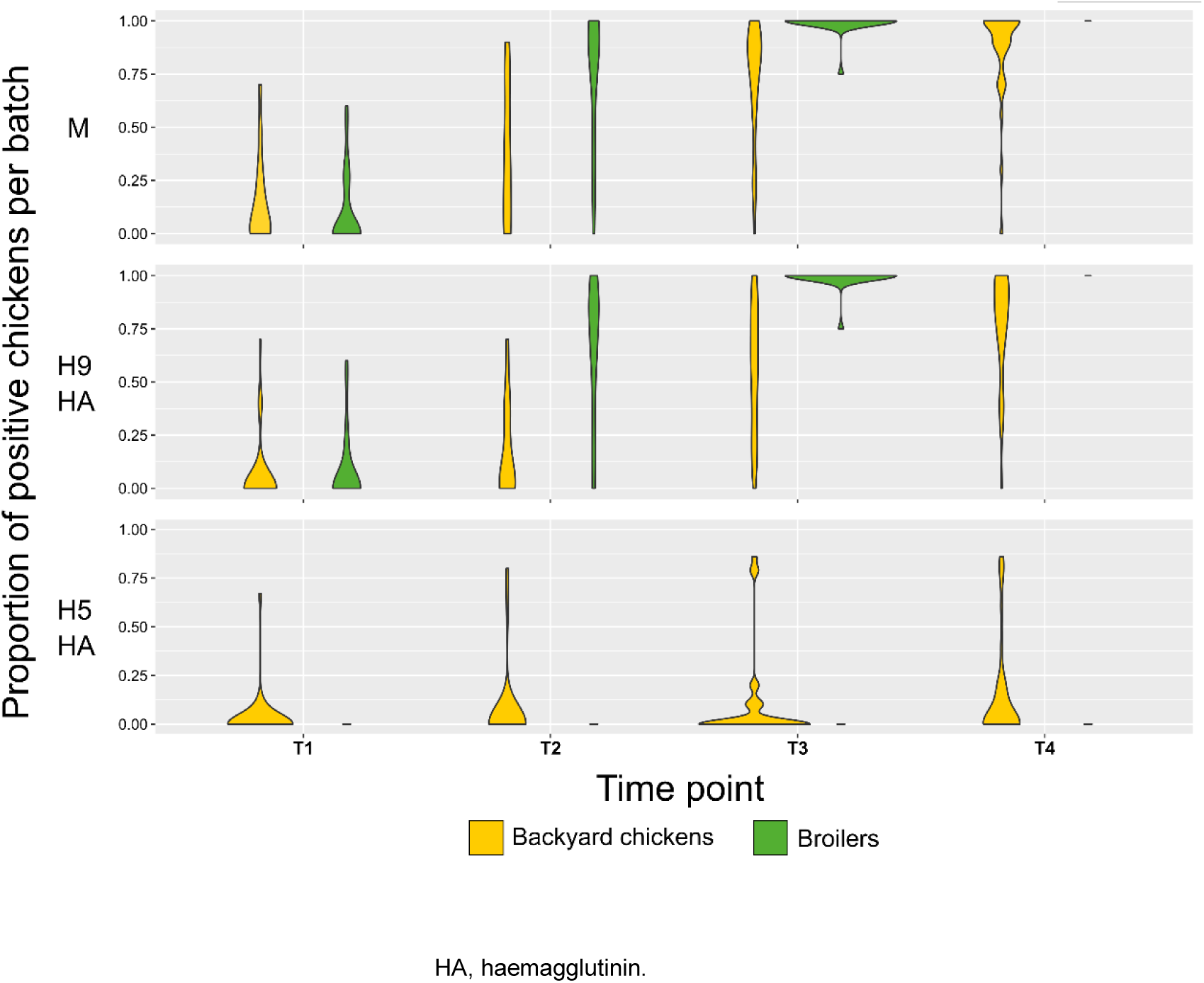
Proportion of positive chickens in a batch by T1–T4

The proportion of AIV-positive broilers at T1 was slightly higher for control than intervention groups, for both M and H9 HA genes and both positivity thresholds. This was also the case for backyard chickens for C_t_<33, whereas the opposite was true for C_t_<40. The intervention was, however, not associated with AIV detection at T1, as all odds ratio (OR) CIs were large and encompassed 1 for all combinations of gene, positivity threshold and stratum (Table 3).

**Table 3.**
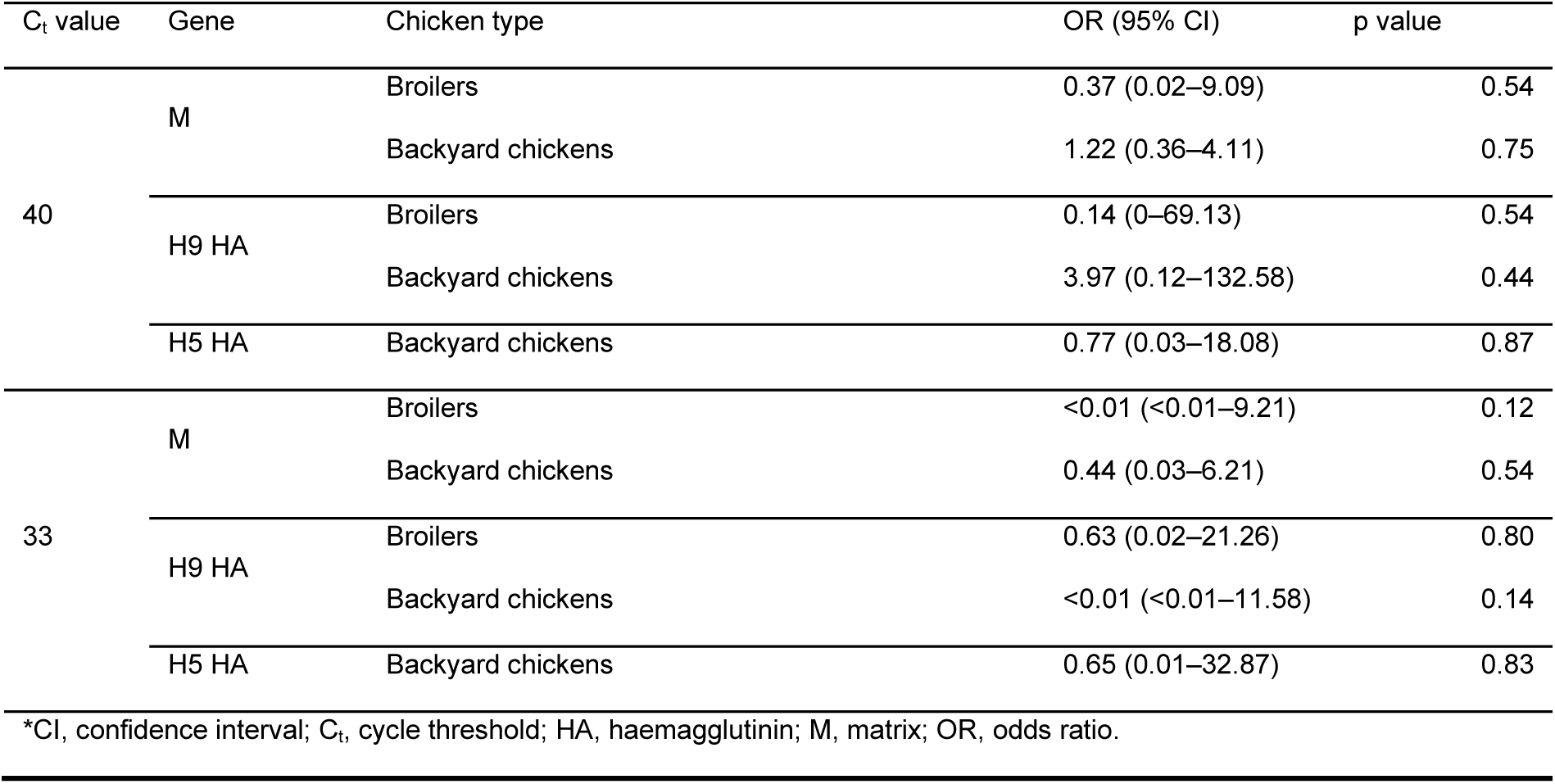
Outputs of the generalised linear mixed models with the intervention as fixed effect at T1*.

Although broilers were less frequently AIV-positive at T1 than backyard chickens, they accounted for a higher proportion of H9-positive chickens. Within 12 h upon arrival at the LBM (T2), the numbers of M- and H9-positive broilers had increased more than 7-fold, and after 36 h (T3) the cumulative incidence was ≥95% for both intervention and control groups. The rise in cumulative incidence was slower for backyard chickens, with a fraction (17.4%) of them having remained negative by the end of the study. Indeed, the numbers of M- and H9-positive backyard chickens increased 2-to-3-fold by T2, and cumulative incidence ranged between 37.8– 71.3% at T3, and between 64.4–86.7% at T4. Likewise, the proportion of positive chickens in affected batches increased over time (Figure 3).

Among broiler batches that tested negative at T1, the evidence of lower odds of AIV detection in intervention than control groups by T2 was stronger for C_t_<33 than C_t_<40 and for the M than the H9 HA gene (M gene, C_t_<33: OR=0.42 [95% CI: 0.20–0.87]) (Table 4). The absence or low number of negative broilers at T3–T4 meant that ORs could not be computed or were associated with wide CIs.

**Table 4.**
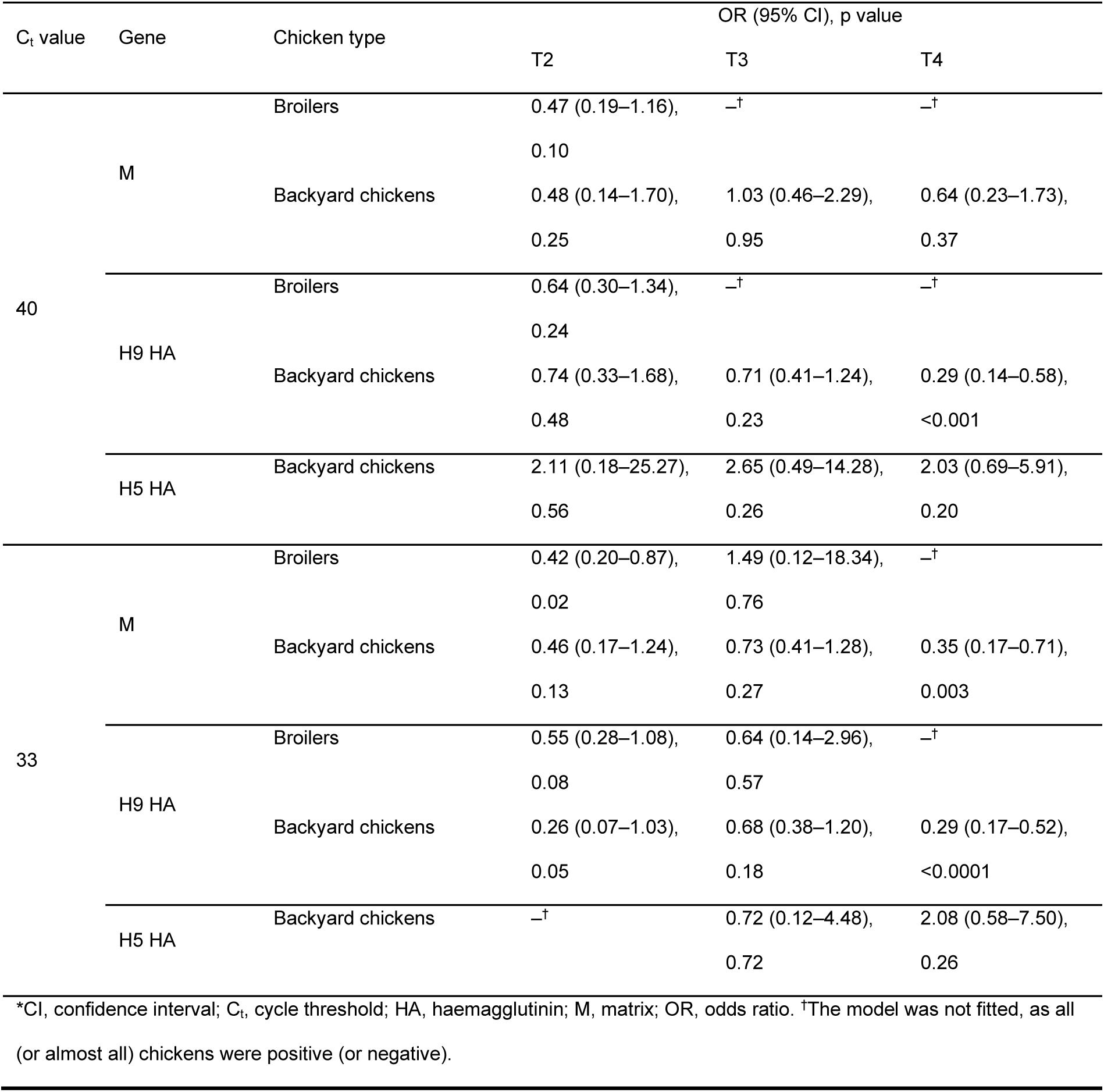
Outputs of the conditional logistic regression models with the intervention as fixed effect at different time points (T2– T4)^*^

There was also stronger evidence that the intervention lowered the odds of H9 detection in backyard chickens by T2 for C_t_<33 than C_t_<40, but the upper bound of the OR remained slightly higher than 1 (95% CI: 0.07–1.03). In contrast to broilers, this effect was stronger at T4, with the intervention being associated with similar OR∼0.3 (C_t_<33: 95% CI: 0.17–0.54) for both positivity thresholds (Table 4).

The intervention was not associated with variations in the odds of detecting the H5 subtype.

## Discussion

Our results suggest that a substantial proportion of chickens sold in LBMs in Chattogram district have already been exposed to AIVs at the farm gate or during transport. By altering usual marketing chains, we demonstrated that reducing the risk of infection along those stages decreased the frequency of AIV detection in market stalls. While there was no apparent effect of the intervention upon arrival at the LBM (T1), it was associated with lower odds of detection at later time points. To our knowledge, this is the first study in which poultry were tested for AIVs upon arrival at the LBM and longitudinally thereafter. A study collecting samples along different stages of poultry marketing chains in Guangdong, China, found that detection rates of AIVs steadily increased and were highest downstream.^23^

The prevalence of AIVs in chickens present in farms and rural households (T0) was higher than previously described.^16,24^ For broilers, all AIV-positive chickens originated from a single farm, whereas the particularly high proportion of positive backyard chickens (14.1%) may have resulted from selection bias. Refusal rate was high and the willingness of rural households to sell chickens may have been associated with ongoing or recent disease outbreaks in the village or their own flock.^25^ Moreover, due to logistic constraints, we recruited backyard chickens in only a small fraction of the catchment area supplying Chattogram city’s LBMs,^5^ where prevalence of infection may be heterogeneous. If rapid diagnostic tests^26^ had been available in the field, we would have been able to ensure that only negative chickens were recruited for the intervention.

While the intervention was not associated with AIV detection at T1, the odds of detecting AIVs at T2 were higher in control than intervention groups. This may have partly been due to the limited sample size and low number of positive chickens at T1, but also due to a higher proportion of chickens in control than intervention groups having been latently infected at T1, resulting in a delayed effect of the intervention.

We introduced a lower positivity threshold, C_t_<33, to increase the likelihood of a positive test result being indicative of infection rather than mere contamination of the birds’ oropharynx^27^ or regressing, past infection.^28,29^ The differences between intervention and control groups being more pronounced at C_t_<33 are further evidence that control chickens were more likely to have already been infected with AIVs when entering the LBM. Selling poultry to multiple stakeholders over several days and transport conditions are likely to promote infection of chickens prior to their delivery at LBMs. In Chattogram city, mobile traders commonly supply multiple LBMs on a single trip, which, in the case of backyard chickens, may even be located in different cities.^5^

Mobile traders’ role in introducing AIVs into LBMs has been described in multiple settings.^30^ In Dhaka city, environmental contamination was higher in LBMs hosting both wholesalers and retailers than retail-only LBMs.^13,31^ Similar observations have been reported from mixed LBMs in eastern China,^32^ where AIVs were also found on transport vehicles.^23^ Moreover, mobile traders and shared transport of birds have been associated with the dissemination of AIVs between LBMs^33,34^ and between farms^35,36^, respectively. Our results also indicate that newly introduced susceptible chickens rapidly become infected and start shedding AIVs. High AIV prevalence in chickens entering LBMs, re-offering of poultry left unsold the previous day^37,38^ – a common practice in Bangladesh^5,12^ –, lack of effective cleaning and disinfection,^12,39^ and the high density at which poultry are kept in market stalls are factors likely promoting a high pressure of infection on susceptible chickens. Further research is required to quantify the relative contribution of those different factors to AIV transmission.

AIV infection spread more slowly among backyard chickens, with some of them, contrary to broilers, remaining negative throughout the study. Due to their longer production period and different environment, backyard chickens were more likely to have already been exposed to multiple AIV subtypes,^16^ conferring them a level of cross- or subtype-specific immune protection. High C_t_ values (>35) were reported during recruitment (T0), suggesting reduced viral shedding due to regressing infection.^28,29^ Moreover, backyard chickens are often thought to be more resistant to infectious diseases compared to other chicken types.^40^ A recent experimental trial demonstrated that the microbiomes of hen-raised chicks (i.e., backyard chickens) remained more stable following exposure to H9N2 than those of chicks raised in isolation, promoting long-lasting immunity.^41^ On the other hand, co-infection with infectious bursal disease has been suspected to have an immunosuppressive effect and increase the incidence of AIVs in broilers.^42^ These findings highlight that different chicken types may play different roles in AIV transmission in LBMs.

Most M-positive chickens in this study were also positive for H9N2. Indeed, H9N2 is the most prevalent subtype affecting marketed chickens in Bangladesh.^13^ The proportion of samples testing positive for the M but not for the H9 HA gene was higher among backyard chickens than broilers, with H5N1 being detected only in the former. This contrast may be attributable to differences in the timing of sampling, as backyard chickens were recruited in March–May, coinciding with the peak period for H5N1 detection in Asia,^43,44^ including Bangladesh,^31^ whereas broilers were sampled from June onwards. Another factor may have been higher exposure of backyard chickens to ducks and wild birds, known risk factors for H5N1 infection,^45–48^ which may have also prompted infection with other AIV subtypes. The prevalence of H5N1 was, however, generally low, possibly resulting from cross-reactive cellular immunity induced by previous exposure to H9N2,^49^ which may have delayed the course of H5N1 infection.^50–52^

We successfully implemented a controlled field experiment to identify the stage of poultry marketing chains at which amplification of AIVs occurs. It is the first experimental trial to create a controlled environment under field conditions, exposing chickens to AIVs in a LBM. Recruiting intervention and control groups from the same farms and rural households would have allowed us to compare chickens with equivalent pre-test exposure. Unfortunately, such study design was not feasible due to the inability to incentivise mobile traders to synchronise their trips with our study protocol. Collecting both oropharyngeal and cloacal swabs could have helped distinguishing infection and shedding from mere contamination of the birds’ oropharynx through, for example, ingestion of contaminated drinking water.^14,53–55^ However, this may not have improved H9N2 detection, for which respiratory shedding is reportedly more common than cloacal shedding.^56–60^

Given that the configuration of poultry marketing chains and characteristics of the associated production systems can vary substantially by poultry type and region,^5^ such study should be conducted in other parts of Bangladesh, as well as in other countries, and involve other chicken or poultry types to fully generalise the results.

## Conclusions

We show that the high AIV prevalence in marketed chickens in Bangladesh is not solely attributable to viral transmission within LBMs but also to infection occurring upstream of marketing chains, prior to chickens’ supply to market stalls. Trade and transport networks should therefore be targeted to complement risk mitigation strategies already implemented in LBMs and farms.^61^

While the implementation of such interventions in the context of complex marketing chains, involving multiple stakeholders, is likely to be challenging, it is imperative to mitigate viral amplification and reduce human exposure to zoonotic AIVs along trade networks.

## Acknowledgements

We thank the research team at Chattogram Veterinary and Animal Sciences University (CVASU), Bangladesh, who assisted with the fieldwork, in particular with animal care, sample collection, and laboratory analysis. Ethical approval was obtained from both CVASU and City University of Hong Kong.

This study was supported by City University of Hong Kong; the BALZAC research programme “Behavioural Adaptations in Live Poultry Trading and Farming Systems and Zoonoses Control in Bangladesh” (Grant No. <u>BB/L018993/1</u>), one of 11 programmes funded under Zoonoses & Emerging livestock Systems (ZELS), a joint research initiative between Biotechnology and Biological Sciences Research Council (BBSRC), Defence Science and Technology Laboratory (DSTL), Department for International Development (DFID), Economic and Social Research Council (ESRC), Medical Research Council (MRC), and Natural Environment Research Council (NERC); the UKRI GCRF One Health Poultry Hub (Grant No. <u>BB/S011269/1</u>), one of 12 interdisciplinary research hubs funded under the UK government’s Grand Challenge Research Fund Interdisciplinary Research Hub initiative. G.F. was supported by the French National Research Agency and the French Ministry of Higher Education and Research.

## Biographical Sketch

Lisa Kohnle is currently working as a Scientific Officer at the European Food Safety Authority (EFSA). During her PhD at City University of Hong Kong, she spent time in Bangladesh to study transmission dynamics of avian influenza viruses in the field. Her research interests focus on the epidemiology of infectious diseases in a One Health context.

## Address for Correspondence

Mailing address: Borgo Collegio Maria Luigia 22, 43121 Parma, Italy

Mobile phone number: +39 3452105169

## Supplemental Materials

Appendices

