## Appendices for "Amplification of avian influenza viruses along poultry marketing chains in Bangladesh: a controlled field experiment"

Article DOI: xxx

### Appendices

#### Appendix A: Study location and period

The south-eastern part of Bangladesh is formed by Chattogram division, which is characterised by an intensifying poultry production in response to the increasing demand for animal-based protein, particularly in its urbanised centres. As our research facilities at Chattogram Veterinary and Animal Sciences University (CVASU) and the live bird markets (LBMs) used in this study were located in Chattogram city, the locations of farms and villages recruited were determined by the catchment area of the city's LBMs.[1] Considering also accessibility and practicability (reachable on a day trip), we included all sub-districts ('upazilas') in Chattogram district (apart from Sandwip, an island, and Karnaphuli): Anwara, Banshkhali, Boalkhali, Chandanaish, Fatikchhari, Hathazari, Lohagara, Mirsharai, Patiya, Rangunia, Raozan, Satkania, and Sitakunda. In addition to those, for backyard chickens, we selected two additional sub-districts – Chhagalnaiya in Feni district and Chakaria in Cox's Bazar district – bordering Chattogram district. Due to security concerns, we were not able to cover the districts in the east of Chattogram division (i.e., Rangamati, Khagrachhari, and Bandarban), along the Chattogram Hill Tracts, despite their substantial role in supplying LBMs with backyard poultry.[2]

An overview of the locations of farms and villages is presented in Appendix Figure 1.

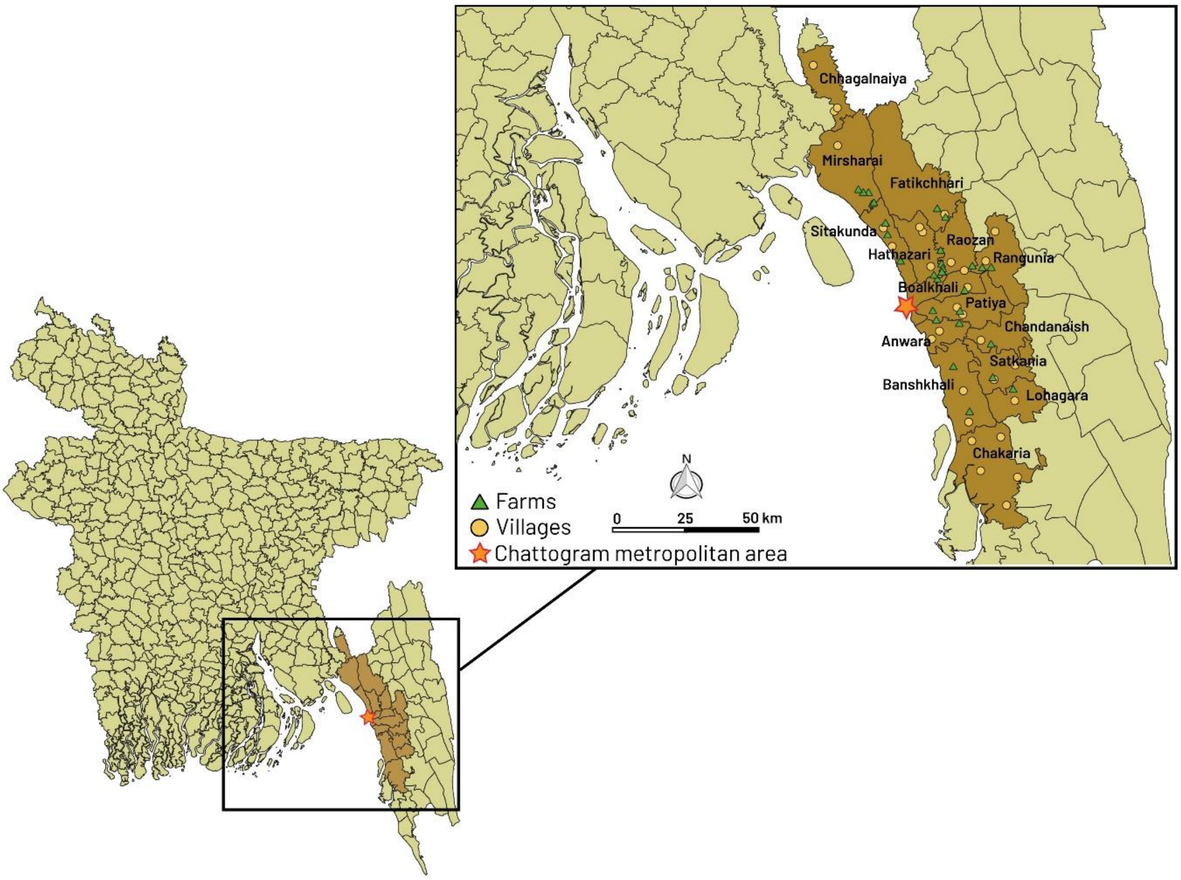

Locations of villages were not available for the first 4 pilot batches.

**Appendix Figure 1.** Map of recruited farms (n=30, green) and villages (n=30, yellow) in selected sub-districts (n=16, brown) in 3 districts of Chattogram division, Bangladesh.

We visited two LBMs in Chattogram city, Pahartali Kacha Bazar and Reazuddin Bazar for recruiting broiler and backyard chicken control groups, respectively. Pahartali Kacha Bazar was also location of the second part of the field experiment.

Backyard chickens were sampled between March–May 2019. After a short break during Ramadan in May, broiler sampling started in June 2019 and was completed before Eid ul-Adha in August 2019.

### Appendix B: Sample size estimation

We aimed to estimate a sample size yielding sufficient statistical power (1–β=0.80) to detect significant differences (α=0.05) in the frequencies of positive birds between intervention and control groups at T1. To do so, we simulated an experiment as follows. Let $c$ refer to the number of chickens in an intervention or control group, $p_g$ to the probability of a chicken population from which an intervention ($p_{g=int}$, a farm or a village) or a control group ($p_{g=cont}$, an LBM) was recruited being infected, and $v_g$ to the probability of a chicken being infected in an infected population. For each batch, the statuses of the two populations from which an intervention group and a control group were recruited were given by Bernoulli trials with probability of success $p_{g=int}$ and $p_{g=cont}$, respectively. For a group recruited from an infected population, the number of PCR-positive chickens was given by a Binomial trial with $c$ as the number of trials and $v_g$ as the probability of a success. $p_{g=int}$ was assumed at 0.1 (exceeding farm-level prevalence reported previously),[3] and $v_{g=int}$ at 0.3. $p_{g=cont}$ was assumed at 0.8 (market-level prevalence was reported to be nearly 100%),[4] and $v_{g=cont}$ at 0.2 (exceeding bird-level prevalence reported previously).[4] The overall numbers of positive and negative chickens for both intervention and control groups were computed and a Fisher's exact test performed. These steps were repeated 10,000 times, with statistical power being the proportion of simulations for which a significant difference was found. Note that $c$=5, as a larger number of birds would not have been manageable with respect to transport and storage. Therefore, to improve statistical power, we increased the number of groups instead of the number of birds within groups.[5]

### Appendix C: Sampling strategy

#### *Selection of sub-districts ('upazilas')*

The number of broiler farms and villages recruited in each sub-district ('upazila') was proportional to the estimated number of chickens they supplied to LBMs in Chattogram city.[6] For broilers, our study area covered 55.5% of the catchment area of LBMs in Chattogram city (Appendix Table 1), compared to only 17.1% for backyard chickens (Appendix Table 2).

**Appendix Table 1.** Estimation of the number of broiler farms recruited per sub-district

| District | Sub-district | Estimated proportion of broilers supplied to Chattogram LBMs | | Estimated number of farms to be recruited† | Rounded (actual) ‡ number of farms to be recruited |
|---|---|---|---|---|---|
| | | Raw* | Normalised | | |
| Chattogram | Anwara | 0.032 | 0.058 | 1.74 | 2 (0) |
| | Banshkhali | 0.030 | 0.054 | 1.62 | 2 (2) |
| | Boalkhali | 0.025 | 0.045 | 1.35 | 1 (1) |
| | Chandanaish | 0.020 | 0.036 | 1.08 | 1 (1) |
| | Fatikchhari | 0.032 | 0.058 | 1.74 | 2 (2) |
| | Hathazari | 0.101 | 0.182 | 5.46 | 6 (6) |
| | Lohagara | 0.023 | 0.041 | 1.23 | 1 (1) |
| | Mirsharai | 0.039 | 0.070 | 2.10 | 2 (3) |
| | Patiya | 0.053 | 0.095 | 2.85 | 3 (3) |
| | Rangunia | 0.025 | 0.045 | 1.35 | 1 (2) |
| | Raozan | 0.079 | 0.142 | 4.26 | 4 (3) |
| | Satkania | 0.019 | 0.034 | 1.02 | 1 (1) |
| | Sitakunda | 0.077 | 0.139 | 4.17 | 4 (5) |
| **Total** | | 0.555 | 1 | 30 | 30 (30) |

*As estimated in[6] †Calculated by multiplying the normalised proportions with the number of recruited farms (n=30) ‡Ad hoc changes were due to contacted feed dealers being unavailable or farms close to the end of the production cycle could not be located in the targeted sub-district.

| Appendix Table 2. Estimation of the number of villages recruited per sub-district | | | | | |
|---|---|---|---|---|---|
| District | Sub-district | Estimated proportion of backyard chickens supplied to Chattogram LBMs | | Estimated number of villages to be recruited† | Rounded number of villages to be recruited |
| | | Raw* | Normalised | | |
| Chattogram | Anwara | 0.009 | 0.053 | 1.59 | 2‡ |
| | Banshkhali | 0.009 | 0.053 | 1.59 | 2‡ |
| | Boalkhali | 0.009 | 0.053 | 1.59 | 1‡ |
| | Chandanaish | 0.009 | 0.053 | 1.59 | 1‡ |
| | Fatikchhari | 0.009 | 0.053 | 1.59 | 1‡ |
| | Hathazari | 0.015 | 0.088 | 2.64 | 3 |
| | Lohagara | 0.009 | 0.053 | 1.59 | 1‡ |
| | Mirsharai | 0.009 | 0.053 | 1.59 | 1‡ |
| | Patiya | 0.009 | 0.053 | 1.59 | 2‡ |
| | Rangunia | 0.012 | 0.070 | 2.10 | 2 |
| | Raozan | 0.011 | 0.064 | 1.92 | 2 |
| | Satkania | 0.009 | 0.053 | 1.59 | 2‡ |
| | Sitakunda | 0.010 | 0.058 | 1.74 | 2 |
| Cox's Bazar | Chakaria | 0.027 | 0.158 | 4.74 | 5 |
| Feni | Chhagalnaiya | 0.015 | 0.088 | 2.64 | 3 |
| **Total** | | 0.171 | 1 | 30 | 30§ |

*As estimated in[6] †Calculated by multiplying the normalised proportions with the number of villages to be recruited (n=30) ‡Randomly rounded up or down. §Four additional villages were recruited as pilot batches, they were located in Hathazari, Raozan and Sitakunda.

#### *Selection of chickens for intervention groups*

For broilers, within commercial farms, if more than one flock was present, the oldest flock was selected. Within flocks, 5 chickens fulfilling the following inclusion criteria were then randomly selected: healthy appearance, upright standing, and medium size. On the other hand, any of the following clinical signs of infectious diseases were exclusion criteria: anorexia, oedematous comb or wattles, sneezing, nasal discharge, swollen eyelids or sinuses, heavy breathing, diarrhoea, haemorrhages, or feather loss.

For backyard chickens, the 'Random points in layer bounds' function in QGIS 3.0.0–Girona (https://qgis.org/en/site/) was used to generate random coordinates within the selected sub-districts. Those coordinates were then entered in Google Maps (https://www.google.com/maps) to verify their location. If a point was adjacent to a neighbouring sub-district, within a national park, or too far from a major road, it was excluded from the study. The nearest village to each generated point, within a radius of 3 km, was selected. Within villages, 5 different rural households willing to sell backyard chickens were then recruited. Only one chicken was recruited at a time and neighbouring rural households were avoided. Within rural households, the same inclusion and exclusion criteria as for broilers applied. However, chickens were at least 3 months of age.

Chickens were always purchased at the current market price based on their weight.

Appendix D: Altered marketing chain (intervention)

We designed a marketing chain alteration (intervention) to reduce the risk of infection for intervention groups between poultry production sites and points of sale to consumers. Instead of mobilising the usual stakeholders involved in poultry transport and trade (e.g., mobile traders), our research team stepped into their shoes and delivered chickens to the LBM. Taking advantage of our own transport vehicle and research facilities, we standardised and implemented the intervention uniformly across all batches. While procedures were evaluated and optimised during the pilot phase of the field experiment, working with checklists and regular training of the research team ensured consistency and continuity.

For each intervention group, the first step was to purchase 5 chickens from either a commercial farm (in case of broilers) or 5 rural households (in case of backyard chickens) on the first day of the field experiment (Table 1). Each of those 5 chickens was uniquely identified with a leg band and sampled on site. A poultry cage was installed on the cargo bed of a pickup truck to facilitate transport of intervention groups, one at a time, to our research facilities at CVASU.

Intermediate stops were avoided to keep transport durations as short as possible. The pickup truck was never left unattended or parked in the vicinity of another farm or LBM. After each trip, we thoroughly cleaned (with water and detergent) and disinfected both pickup truck and accessories to reduce the risk of cross-contamination between successive intervention groups. Peroxygen-based Virkon$^{TM}$ S (LANXESS) was applied to all surfaces once dry.[7] Our research team disinfected their shoes after each trip and was required to both shower and change clothes between successive intervention groups. With the aim of reducing stress levels of chickens during transport, we provided feed and water ad libitum. We also placed a canvas cover on top of the poultry cage to protect intervention groups from adverse weather conditions (e.g., sun or rainfall).

The second step was to store intervention groups in our research facilities at CVASU for 2.5 d (second and third days of the field experiment, Table 1). For this purpose, we designed and commissioned the construction of a poultry shed on CVASU's main campus in Chattogram city.

The shed consisted of 4 isolated compartments with the capacity to host up to 4 intervention groups at the same time. We chose specific surface materials which are easy to clean and disinfect (cement for the floor and metal sheets for the walls), whereas bamboo was used to construct the frame. We ensured sufficient air circulation by adding wire mesh on the front doors and back, whereas fans were not provided. Drains were installed in the back of each compartment to facilitate drainage of cleaning water and covered by wire mesh to prevent rodents from entering the shed. Individual footbaths were constructed in front of each compartment to reduce pathogen introduction from outside. They were filled with a Virkon$^{TM}$ S (LANXESS) solution, which was replenished each morning and after rainfall. A canvas cover to protect chickens from adverse weather conditions and during the night completed the shed. Each compartment was further divided into a space for chickens and a space for storing an individual set of feeding (poultry feeder and drinker, feed and water containers) and cleaning tools (buckets and brushes). Those tools always remained inside their respective compartments and were never exchanged with each other. Bedding material was not provided. Chickens received water ad libitum and either industrial feed (in case of broilers) or raw rice (in case of backyard chickens). Each compartment was locked and assigned to an individual animal caretaker, who was the only one allowed to enter and interact with the respective intervention group for the following 2.5 d. Occupied compartments were cleaned with water and detergent once per day. More thorough cleaning and disinfection with Virkon™ S (LANXESS) (after surfaces had dried) was carried out each time an intervention group left the shed. A rotation system guaranteed that each compartment remained empty for 1–2 d between intervention groups. During those days, feeding and cleaning tools were soaked in disinfectant. Nevertheless, there were never more than 3 compartments occupied at the same time. In addition, lime (calcium oxide) was spread around the shed every 3–4 d. All those procedures aimed to reduce cross-contamination between intervention groups and pathogen introduction from the environment. Our animal caretakers closely monitored the chickens and reported any suspicious clinical signs to our research team. They visited the shed at least once per day and were required to comply with a rigorous set of biosecurity measures (e.g., washing hands frequently, showering and changing clothes between successive intervention groups, avoiding the LBM).

The last step was to deliver intervention groups to the LBM in the early morning (before 7 AM) of the fourth day of the field experiment (Table 1), using a public compressed natural gas (CNG) vehicle. As LBMs are highly contaminated with AIVs, we avoided using the pickup truck for the second part of the field experiment.

Appendix E: Sample collection

Each chicken recruited in this study was clearly identified with a leg band (intervention and control groups on the left and right leg, respectively) and always sampled individually. We collected oropharyngeal swab samples from those chickens at each time point (Table 1) to detect the presence of avian influenza viruses (AIVs) in the birds' respiratory tract. To obtain enough mucous, the trachea, choanal cleft, and upper palate were swabbed with a cotton swab. Our research team always wore sterile gloves and disinfected their hands between individual birds. We immediately placed swabs into individual sterile tubes with 3.5 mL viral transport medium (recipe below) and broke off the stick before screwing the cap. All tubes had been labelled with a unique identifier specifying the respective batch, stratum, chicken, and time point. After all swabs were collected, we immediately stored the tubes in a cool box to keep temperature at about 4°C during transport. At CVASU, we then extracted 0.7 mL from each tube and produced pools of 5 samples. However, all individual samples were kept and have since been stored at –80°C.

*Preparation of viral transport medium*

The viral transport medium preparation followed the Standard Operating Procedure of the UK Animal and Plant Health Agency (Ref. BPU 1551; Implementation date 27/11/2013), with penicillin G being replaced with benzathine penicillin.

Appendix F: Laboratory analysis

*Testing algorithm*

All samples collected during the first part of the field experiment (i.e., T0 and T1) were tested. As we aimed to detect the first time point at which a chicken became positive, samples collected during the second part (i.e., after T1) were tested against a given gene if samples collected at previous time points for the same chicken were negative for that gene. To minimise concerns about a positive sample having resulted from contamination of the bird's oropharynx,[8] we continued to test samples from chickens for which $C_t$ values were >33 at the preceding time point. The testing algorithm is presented in Appendix Figure 2. For the purpose of quality control, we retested >50 samples at random.

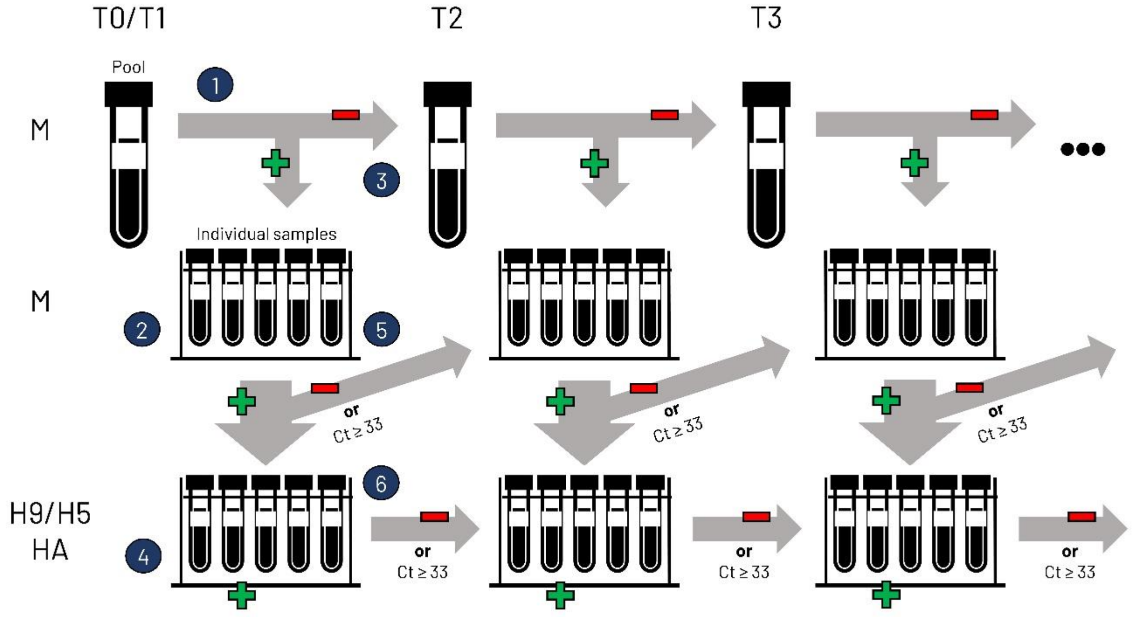

Ct, cycle threshold; HA, haemagglutinin; M, matrix; T, time point.

**Appendix Figure 2.** Testing algorithm for pools and individual samples. [1] test all pools of the first two time points (i.e., T0 and T1) for the M gene, [2] if a pool is positive, test all individual samples of the pool for the M gene, [3] if a pool is negative, test the pool of the following time point for the M gene and then continue with either [2] or [3], [4] test individual samples positive for the M gene for both H9 and H5 HA genes, [5] if individual samples are negative for the M gene or have a Ct⩾33, test all individual samples of the following time point for the M gene and then continue with [4] or [5],[6] if individual samples are negative for the H9/H5 HA gene or have a Ct⩾33, test individual samples of the following time point for the H9/H5 HA gene.

Seventy-five T0–T1 samples forming 13 control or intervention groups and belonging to 8 successive broiler batches were associated with similar $C_t$ values (ranging between 31 and 32). As the same $C_t$ values were obtained after retesting, including after re-extraction of viral RNA, contamination of the original samples during the first viral RNA extraction was considered the most plausible explanation. These samples were excluded, as well as all T2–T4 samples from the 5 batches (Batch no. 42–46) for which T1 samples were suspected to be contaminated.

Appendix G: Laboratory results

Appendix Figure 3 displays the detailed rRT-PCR test results (including $C_t$ values for the M, H9 HA, and H5 HA genes) for all time points (T0–T4).

Each row in those tables corresponds to an individual bird, while birds are arranged in groups (intervention or control) of 5 and batches of 10. Sample IDs comprise of all relevant information:

- Batch no. (1–65)
- Stratum (A=broilers, B=backyard chickens)
- Group (1=intervention, 0=control)
- Bird no. (1–650)
- Time point (0–4).

For example, the sample with sample ID '19B1–183–2' belongs to a backyard chicken (bird no. 183 in batch no. 19) in the intervention group and was collected at T2.

The first 4 batches (batch no. 1–4) were pilot batches and therefore not considered in the targeted sample size of 60 batches (see Appendix B). In addition, batch no. 55 was excluded after T0 and replaced with another batch, as all birds in the intervention group had suddenly died after experiencing thermal stress during transport. All birds in batch no. 7 dropped out of the field experiment after T3 when the stallholder accidentally sold them too soon. Contamination of samples during viral RNA extraction was suspected for a total of 75 samples (batch no. 45–49 at T0 and batch no. 42–46 at T1).

A total of 19 backyard chickens and 3 broilers were positive for AIVs other than the H9 or H5 subtypes.

| T0 | | | | | | T1 | | | | | | T2 | | | | | | T3 | | | | | | T4 | | | | | |
|---|---|---|---|---|---|---|---|---|---|---|---|---|---|---|---|---|---|---|---|---|---|---|---|---|---|---|---|---|---|
| Pool ID | Ct (M) | Sample ID | Ct (M) | Ct (H5 HA) | Ct (H9 HA) | Pool ID | Ct (M) | Sample ID | Ct (M) | Ct (H5 HA) | Ct (H9 HA) | Pool ID | Ct (M) | Sample ID | Ct (M) | Ct (H5 HA) | Ct (H9 HA) | Pool ID | Ct (M) | Sample ID | Ct (M) | Ct (H5 HA) | Ct (H9 HA) | Pool ID | Ct (M) | Sample ID | Ct (M) | Ct (H5 HA) | Ct (H9 HA) |
| | | 1B1-001-0 | | | | | | 1B1-001-1 | | | | | | 1B1-001-2 | | | | | | 1B1-001-3 | 32.81 | x | 34.52 | | | 1B1-001-4 | | x | 35.21 |
| | | 1B1-002-0 | | | | | | 1B1-002-1 | | | | | | 1B1-002-2 | | | | | | 1B1-002-3 | x | | | | | 1B1-002-4 | x | | |
| 1B1-0 | x | 1B1-003-0 | | | | 1B1-1 | x | 1B1-003-1 | | | | 1B1-2 | x | 1B1-003-2 | | | | 1B1-3 | 26.32 | 1B1-003-3 | x | | | 1B1-4 | | 1B1-003-4 | x | | |
| | | 1B1-004-0 | | | | | | 1B1-004-1 | | | | | | 1B1-004-2 | | | | | | 1B1-004-3 | 36.27 | x | 41.58 | | | 1B1-004-4 | 27.91 | x | 32.02 |
| | | 1B1-005-0 | | | | | | 1B1-005-1 | | | | | | 1B1-005-2 | | | | | | 1B1-005-3 | x | | | | | 1B1-005-4 | x | | |
| | | | | | | | | 1B0-006-1 | | | | | | 1B0-006-2 | | | | | | 1B0-006-3 | 21.86 | x | 23.96 | | | 1B0-006-4 | | x | |
| | | | | | | | | 1B0-007-1 | | | | | | 1B0-007-2 | | | | | | 1B0-007-3 | 33.31 | x | 35.13 | | | 1B0-007-4 | 24.84 | x | 27.16 |
| | | - | | | | 1B0-1 | x | 1B0-008-1 | | | | 1B0-2 | x | 1B0-008-2 | | | | 1B0-3 | 37.34 | 1B0-008-3 | 28.35 | x | 29.97 | 1B0-4 | | 1B0-008-4 | | x | |
| | | | | | | | | 1B0-009-1 | | | | | | 1B0-009-2 | | | | | | 1B0-009-3 | x | | | | | 1B0-009-4 | x | | |
| | | | | | | | | 1B0-010-1 | | | | | | 1B0-010-2 | | | | | | ~~1B0-010-3~~ | | | | | | ~~1B0-010-4~~ | | | |
| | | 2B1-011-0 | x | | | | | 2B1-011-1 | x | | | | | 2B1-011-2 | x | | | | | 2B1-011-3 | x | | | | | 2B1-011-4 | 38.12 | x | x |
| | | 2B1-012-0 | x | | | | | 2B1-012-1 | x | | | | | 2B1-012-2 | 32.66 | x | x | | | 2B1-012-3 | x | | | | | 2B1-012-4 | x | | |
| 2B1-0 | | 2B1-013-0 | x | | | 2B1-1 | | 2B1-013-1 | 29.49 | x | 39.47 | 2B1-2 | | 2B1-013-2 | x | | | 2B1-3 | | 2B1-013-3 | x | | | 2B1-4 | | 2B1-013-4 | x | | |
| | | 2B1-014-0 | x | | | | | 2B1-014-1 | x | | | | | 2B1-014-2 | x | | | | | 2B1-014-3 | 35.20 | x | x | | | 2B1-014-4 | 37.41 | x | x |
| | | 2B1-015-0 | 36.30 | x | x | | | 2B1-015-1 | 38.12 | x | x | | | 2B1-015-2 | x | | | | | 2B1-015-3 | x | | | | | 2B1-015-4 | x | | |
| | | | | | | | | 2B0-016-1 | x | | | | | 2B0-016-2 | 30.58 | x | x | | | 2B0-016-3 | x | | | | | 2B0-016-4 | 32.87 | 29.39 | x |
| | | | | | | | | 2B0-017-1 | x | | | | | 2B0-017-2 | x | | | | | 2B0-017-3 | 26.54 | 40.05 | 32.34 | | | 2B0-017-4 | 32.45 | x | 37.13 |
| | | - | | | | 2B0-1 | | 2B0-018-1 | x | | | 2B0-2 | | 2B0-018-2 | x | | | 2B0-3 | | 2B0-018-3 | x | | | 2B0-4 | | 2B0-018-4 | 37.64 | x | x |
| | | | | | | | | 2B0-019-1 | x | | | | | 2B0-019-2 | x | | | | | 2B0-019-3 | 32.68 | x | x | | | 2B0-019-4 | 28.72 | x | 32.26 |
| | | | | | | | | 2B0-020-1 | x | | | | | 2B0-020-2 | 36.65 | x | x | | | 2B0-020-3 | 32.02 | x | x | | | 2B0-020-4 | 29.85 | x | 39.78 |
| | | 3B1-021-0 | x | | | | | 3B1-021-1 | 30.11 | 32.99 | x | | | 3B1-021-2 | 32.47 | 35.17 | x | | | 3B1-021-3 | 22.59 | 24.98 | x | | | 3B1-021-4 | 23.51 | 26.30 | x |
| | | 3B1-022-0 | x | | | | | 3B1-022-1 | 32.39 | 34.74 | x | | | 3B1-022-2 | x | 42.84 | | | | 3B1-022-3 | 29.19 | 31.58 | x | | | 3B1-022-4 | 27.47 | 30.30 | x |
| 3B1-0 | | 3B1-023-0 | 33.38 | x | x | 3B1-1 | | ~~3B1-023-1~~ | | | | 3B1-2 | | ~~3B1-023-2~~ | | | | 3B1-3 | | ~~3B1-023-3~~ | | | | ~~3B1-4~~ | | ~~3B1-023-4~~ | | | |
| | | 3B1-024-0 | x | | | | | 3B1-024-1 | 25.09 | 27.59 | x | | | 3B1-024-2 | 24.90 | 28.72 | x | | | 3B1-024-3 | 21.12 | 24.40 | x | | | 3B1-024 | 22.18 | 25.54 | 34.18 |
| | | 3B1-025-0 | x | | | | | 3B1-025-1 | 22.45 | 24.30 | x | | | ~~3B1-025-2~~ | | | | | | ~~3B1-025-3~~ | | | | | | ~~3B1-025-4~~ | | | |
| | | | | | | | | 3B0-026-1 | x | | | | | 3B0-026-2 | 31.72 | 34.75 | x | | | 3B0-026-3 | 33.80 | 38.22 | x | | | 3B0-026-4 | 44.29 | x | x |
| | | | | | | | | 3B0-027-1 | x | | | | | 3B0-027-2 | 38.45 | x | x | | | 3B0-027-3 | 36.95 | x | x | | | 3B0-027-4 | 37.93 | x | x |
| | | - | | | | 3B0-1 | | 3B0-028-1 | x | | | 3B0-2 | | 3B0-028-2 | x | | | 3B0-3 | | 3B0-028-3 | 32.46 | 36.30 | x | ~~3B0-4~~ | | 3B0-028 | 18.11 | 21.95 | x |
| | | | | | | | | 3B0-029-1 | 22.02 | 24.47 | x | | | ~~3B0-029-2~~ | | | | | | ~~3B0-029-3~~ | | | | | | ~~3B0-029-4~~ | | | |
| | | | | | | | | 3B0-030-1 | 32.99 | 36.71 | x | | | 3B0-030-2 | 32.91 | 36.76 | x | | | 3B0-030-3 | 31.82 | 34.92 | 38.89 | | | 3B0-030 | 18.69 | 21.73 | 25.34 |
| | | 4B1-031-0 | | | | | | 4B1-031-1 | | | | | | 4B1-031-2 | 34.80 | 36.02 | x | | | 4B1-031-3 | x | x | x | | | 4B1-031-4 | x | x | x |
| | | 4B1-032-0 | | | | | | 4B1-032-1 | | | | | | 4B1-032-2 | 34.02 | 37.58 | x | | | 4B1-032-3 | x | x | x | | | 4B1-032-4 | x | x | x |
| 4B1-0 | x | 4B1-033-0 | | | | 4B1-1 | x | 4B1-033-1 | | | | 4B1-2 | 33.28 | 4B1-033-2 | 32.55 | 36.66 | x | 4B1-3 | | 4B1-033-3 | | 32.31 | x | 4B1-4 | | 4B1-033-4 | x | x | x |
| | | 4B1-034-0 | | | | | | 4B1-034-1 | | | | | | 4B1-034-2 | 30.09 | 33.91 | x | | | 4B1-034-3 | | 36.54 | x | | | 4B1-034-4 | | 37.25 | x |
| | | 4B1-035-0 | | | | | | 4B1-035-1 | | | | | | 4B1-035-2 | 35.57 | 37.10 | x | | | 4B1-035-3 | 40.30 | x | x | | | 4B1-035-4 | | 36.69 | x |
| | | | | | | | | 4B0-036-1 | | | | | | 4B0-036-2 | 34.50 | 37.19 | x | | | 4B0-036-3 | 37.69 | x | x | | | 4B0-036-4 | 23.21 | x | 26.07 |
| | | | | | | | | 4B0-037-1 | | | | | | 4B0-037-2 | x | | | | | 4B0-037-3 | x | | | | | 4B0-037 | 25.42 | x | 29.61 |
| | | - | | | | 4B0-1 | x | 4B0-038-1 | | | | 4B0-2 | 38.17 | 4B0-038-2 | 33.88 | 37.07 | x | 4B0-3 | | 4B0-038-3 | x | x | x | 4B0-4 | | 4B0-038-4 | x | x | 27.30 |
| | | | | | | | | 4B0-039-1 | | | | | | 4B0-039-2 | 32.70 | 35.47 | x | | | 4B0-039-3 | | x | 27.60 | | | 4B0-039-4 | 21.74 | x | |
| | | | | | | | | 4B0-040-1 | | | | | | 4B0-040-2 | x | | | | | 4B0-040-3 | x | | | | | 4B0-040-4 | x | | |

**T0**

| Pool ID | Ct (M) | Sample ID | Ct (M) | Ct (H5 HA) | Ct (H9 HA) |
|---|---|---|---|---|---|
| 5B1-0 | x | 5B1-041-0 | | | |
| | | 5B1-042-0 | | | |
| | | 5B1-043-0 | | | |
| | | 5B1-044-0 | | | |
| | | 5B1-045-0 | | | |
| - | | | | | |
| 6B1-0 | x | 6B1-051-0 | | | |
| | | 6B1-052-0 | | | |
| | | 6B1-053-0 | | | |
| | | 6B1-054-0 | | | |
| | | 6B1-055-0 | | | |
| - | | | | | |
| 7B1-0 | x | 7B1-061-0 | | | |
| | | 7B1-062-0 | | | |
| | | 7B1-063-0 | | | |
| | | 7B1-064-0 | | | |
| | | 7B1-065-0 | | | |
| - | | | | | |
| 8B1-0 | 32.80 | 8B1-071-0 | x | | |
| | | 8B1-072-0 | x | | |
| | | 8B1-073-0 | 39.97 | x | x |
| | | 8B1-074-0 | x | | |
| | | 8B1-075-0 | x | | |
| - | | | | | |
| 9B1-0 | x | 9B1-081-0 | | | |
| | | 9B1-082-0 | | | |
| | | 9B1-083-0 | | | |
| | | 9B1-084-0 | | | |
| | | 9B1-085-0 | | | |
| - | | | | | |

**T1**

| Pool ID | Ct (M) | Sample ID | Ct (M) | Ct (H5 HA) | Ct (H9 HA) |
|---|---|---|---|---|---|
| 5B1-1 | x | 5B1-041-1 | | | |
| | | 5B1-042-1 | | | |
| | | 5B1-043 | x | | |
| | | 5B1-044-1 | | | |
| | | 5B1-045-1 | | | |
| 5B0-1 | x | 5B0-046-1 | | | |
| | | 5B0-047-1 | | | |
| | | 5B0-048-1 | | | |
| | | 5B0-049-1 | | | |
| | | 5B0-050-1 | | | |
| 6B1-1 | x | 6B1-051-1 | | | |
| | | 6B1-052-1 | | | |
| | | 6B1-053-1 | | | |
| | | 6B1-054-1 | | | |
| | | 6B1-055-1 | | | |
| 6B0-1 | 27.45 | 6B0-056-1 | 24.58 | x | x |
| | | 6B0-057-1 | x | | |
| | | 6B0-058-1 | 43.49 | | |
| | | 6B0-059-1 | x | | |
| | | 6B0-060-1 | x | | |
| 7B1-1 | x | 7B1-061-1 | | | |
| | | 7B1-062-1 | | | |
| | | 7B1-063-1 | | | |
| | | 7B1-064-1 | | | |
| | | 7B1-065-1 | | | |
| 7B0-1 | x | 7B0-066-1 | | | |
| | | 7B0-067-1 | | | |
| | | 7B0-068-1 | | | |
| | | 7B0-069-1 | | | |
| | | 7B0-070-1 | | | |
| 8B1-1 | 33.47 | 8B1-071-1 | 38.81 | x | 40.57 |
| | | 8B1-072-1 | x | | |
| | | 8B1-073-1 | x | | |
| | | 8B1-074-1 | x | | |
| | | 8B1-075-1 | x | | |
| 8B0-1 | x | 8B0-076-1 | | | |
| | | 8B0-077-1 | | | |
| | | 8B0-078-1 | | | |
| | | 8B0-079-1 | | | |
| | | 8B0-080-1 | | | |
| 9B1-1 | x | 9B1-081-1 | | | |
| | | 9B1-082-1 | | | |
| | | 9B1-083-1 | | | |
| | | 9B1-084-1 | | | |
| | | 9B1-085-1 | | | |
| 9B0-1 | x | 9B0-086-1 | | | |
| | | 9B0-087-1 | | | |
| | | 9B0-088-1 | | | |
| | | 9B0-089-1 | | | |
| | | 9B0-090-1 | | | |

**T2**

| Pool ID | Ct (M) | Sample ID | Ct (M) | Ct (H5 HA) | Ct (H9 HA) |
|---|---|---|---|---|---|
| 5B1-2 | 42.11 | 5B1-041-2 | x | | |
| | | 5B1-042-2 | x | | |
| | | ~~5B1-043-2~~ | | | |
| | | 5B1-044-2 | x | | |
| | | 5B1-045-2 | x | | |
| 5B0-2 | 25.11 | 5B0-046-2 | 20.24 | x | 20.29 |
| | | 5B0-047-2 | x | | |
| | | 5B0-048-2 | 25.49 | x | 25.50 |
| | | 5B0-049-2 | x | | |
| | | 5B0-050-2 | x | | |
| 6B1-2 | 34.61 | 6B1-051-2 | 32.38 | x | 37.97 |
| | | 6B1-052-2 | 34.31 | x | x |
| | | 6B1-053-2 | 42.11 | | |
| | | 6B1-054-2 | 35.69 | x | x |
| | | 6B1-055-2 | x | | |
| 6B0-2 | | 6B0-056-2 | 20.21 | x | 22.76 |
| | | 6B0-057-2 | 36.45 | x | x |
| | | 6B0-058-2 | 36.62 | x | x |
| | | 6B0-059-2 | 32.13 | x | 36.81 |
| | | 6B0-060-2 | 25.52 | x | 27.80 |
| 7B1-2 | x | 7B1-061-2 | | | |
| | | 7B1-062-2 | | | |
| | | 7B1-063-2 | | | |
| | | 7B1-064-2 | | | |
| | | 7B1-065-2 | | | |
| 7B0-2 | 37.96 | 7B0-066-2 | x | | |
| | | 7B0-067-2 | 33.37 | 30.31 | x |
| | | 7B0-068-2 | 33.41 | x | x |
| | | 7B0-069-2 | x | | |
| | | 7B0-070-2 | 33.84 | x | x |
| 8B1-2 | | 8B1-071-2 | 36.51 | x | 36.88 |
| | | 8B1-072-2 | x | | |
| | | 8B1-073-2 | x | | |
| | | 8B1-074-2 | x | x | x |
| | | 8B1-075-2 | 33.37 | x | 36.12 |
| 8B0-2 | 26.72 | 8B0-076-2 | 33.41 | x | 38.53 |
| | | 8B0-077-2 | x | | |
| | | 8B0-078-2 | x | | |
| | | 8B0-079-2 | x | | |
| | | 8B0-080-2 | 37.96 | x | 22.36 |
| 9B1-2 | x | 9B1-081-2 | | | |
| | | 9B1-082-2 | | | |
| | | 9B1-083-2 | | | |
| | | 9B1-084-2 | | | |
| | | 9B1-085-2 | | | |
| 9B0-2 | x | 9B0-086-2 | | | |
| | | 9B0-087-2 | | | |
| | | 9B0-088-2 | | | |
| | | 9B0-089-2 | | | |
| | | 9B0-090-2 | | | |

**T3**

| Pool ID | Ct (M) | Sample ID | Ct (M) | Ct (H5 HA) | Ct (H9 HA) |
|---|---|---|---|---|---|
| 5B1-3 | | 5B1-041-3 | 33.52 | x | 32.12 |
| | | 5B1-042-3 | 27.93 | x | 29.98 |
| | | ~~5B1-043-3~~ | | | |
| | | 5B1-044-3 | 35.37 | x | 35.88 |
| | | 5B1-045-3 | 24.39 | x | 25.75 |
| 5B0-3 | | 5B0-046-3 | | x | |
| | | 5B0-047-3 | 21.59 | x | 23.57 |
| | | 5B0-048-3 | | x | |
| | | 5B0-049-3 | x | | |
| | | 5B0-050-3 | 36.98 | x | 36.84 |
| 6B1-3 | | 6B1-051-3 | | x | 28.32 |
| | | 6B1-052-3 | 36.66 | x | 33.94 |
| | | 6B1-053-3 | 34.87 | x | 35.43 |
| | | 6B1-054-3 | 24.66 | x | 26.18 |
| | | 6B1-055-3 | 35.65 | x | 37.28 |
| 6B0-3 | | 6B0-056-3 | | x | |
| | | 6B0-057-3 | 20.51 | x | 22.63 |
| | | 6B0-058-3 | 27.21 | x | 29.02 |
| | | 6B0-059-3 | | x | 24.84 |
| | | 6B0-060-3 | | x | |
| 7B1-3 | | 7B1-061-3 | 34.40 | x | 36.92 |
| | | 7B1-062-3 | 37.09 | x | 44.63 |
| | | 7B1-063-3 | x | | |
| | | 7B1-064-3 | 34.06 | x | 35.28 |
| | | 7B1-065-3 | 29.77 | x | 31.20 |
| 7B0-3 | | 7B0-066-3 | 38.29 | x | x |
| | | 7B0-067-3 | 30.67 | 32.93 | x |
| | | 7B0-068-3 | 28.98 | x | 30.87 |
| | | 7B0-069-3 | 38.47 | x | 41.40 |
| | | 7B0-070-3 | 31.84 | x | 33.83 |
| 8B1-3 | | 8B1-071-3 | 37.98 | x | 36.59 |
| | | 8B1-072-3 | 32.74 | x | 34.05 |
| | | 8B1-073-3 | 36.94 | x | 39.95 |
| | | 8B1-074-3 | x | x | x |
| | | 8B1-075-3 | 35.24 | x | 34.23 |
| 8B0-3 | | 8B0-076-3 | x | x | 37.72 |
| | | 8B0-077-3 | 30.66 | x | 33.30 |
| | | 8B0-078-3 | 29.63 | x | 32.00 |
| | | 8B0-079-3 | 23.54 | x | 25.23 |
| | | 8B0-080-3 | 23.48 | x | |
| 9B1-3 | 26.32 | 9B1-081-3 | x | | |
| | | 9B1-082-3 | 34.94 | x | 34.84 |
| | | 9B1-083-3 | x | | |
| | | 9B1-084-3 | 36.85 | x | x |
| | | 9B1-085-3 | x | | |
| 9B0-3 | x | 9B0-086-3 | | | |
| | | 9B0-087-3 | | | |
| | | 9B0-088-3 | | | |
| | | 9B0-089-3 | | | |
| | | 9B0-090-3 | | | |

**T4**

| Pool ID | Ct (M) | Sample ID | Ct (M) | Ct (H5 HA) | Ct (H9 HA) |
|---|---|---|---|---|---|
| 5B1-4 | | 5B1-041-4 | 31.61 | x | |
| | | 5B1-042-4 | | x | |
| | | ~~5B1-043-4~~ | | | |
| | | 5B1-044-4 | 27.73 | x | 33.25 |
| | | 5B1-045-4 | | x | |
| 5B0-4 | | 5B0-046-4 | | x | |
| | | 5B0-047-4 | | x | |
| | | 5B0-048-4 | | x | |
| | | 5B0-049 | 34.67 | x | 30.98 |
| | | 5B0-050-4 | 32.03 | x | 32.32 |
| 6B1-4 | | 6B1-051-4 | | x | |
| | | 6B1-052-4 | 34.49 | 39.93 | 40.84 |
| | | 6B1-053-4 | 31.08 | 27.96 | 33.27 |
| | | 6B1-054-4 | | x | |
| | | 6B1-055-4 | 35.19 | x | 38.59 |
| 6B0-4 | | 6B0-056-4 | | x | |
| | | 6B0-057-4 | | x | |
| | | 6B0-058-4 | | x | |
| | | 6B0-059-4 | | x | |
| | | 6B0-060-4 | | x | |
| ~~7B1-4~~ | | ~~7B1-061-4~~ | | | |
| | | ~~7B1-062-4~~ | | | |
| | | ~~7B1-063-4~~ | | | |
| | | ~~7B1-064-4~~ | | | |
| | | ~~7B1-065-4~~ | | | |
| ~~7B0-4~~ | | ~~7B0-066-4~~ | | | |
| | | ~~7B0-067-4~~ | | | |
| | | ~~7B0-068-4~~ | | | |
| | | ~~7B0-069-4~~ | | | |
| | | ~~7B0-070-4~~ | | | |
| 8B1-4 | | 8B1-071-4 | 31.70 | x | 31.07 |
| | | 8B1-072-4 | | x | 33.53 |
| | | 8B1-073-4 | 30.13 | 39.58 | 32.07 |
| | | 8B1-074-4 | x | | |
| | | 8B1-075-4 | 35.82 | x | 37.10 |
| 8B0-4 | | 8B0-076-4 | x | x | 37.46 |
| | | 8B0-077-4 | | x | 33.98 |
| | | 8B0-078 | | 21.88 | |
| | | 8B0-079-4 | | x | |
| | | 8B0-080-4 | | x | |
| 9B1-4 | | 9B1-081-4 | 26.38 | x | 32.09 |
| | | 9B1-082-4 | 24.73 | x | 26.63 |
| | | 9B1-083-4 | 35.24 | x | 41.73 |
| | | 9B1-084-4 | 26.73 | x | 28.71 |
| | | 9B1-085-4 | 21.26 | x | 44.80 |
| 9B0-4 | 25.38 | 9B0-086-4 | 21.36 | x | 23.98 |
| | | 9B0-087-4 | 22.69 | x | 23.34 |
| | | 9B0-088-4 | 25.24 | x | 26.31 |
| | | 9B0-089-4 | 27.48 | x | 28.39 |
| | | 9B0-090-4 | 25.03 | x | 25.78 |

| T0 | | | | | | T1 | | | | | | T2 | | | | | | T3 | | | | | | T4 | | | | | |
|---|---|---|---|---|---|---|---|---|---|---|---|---|---|---|---|---|---|---|---|---|---|---|---|---|---|---|---|---|---|
| Pool ID | Ct (M) | Sample ID | Ct (M) | Ct (H5 HA) | Ct (H9 HA) | Pool ID | Ct (M) | Sample ID | Ct (M) | Ct (H5 HA) | Ct (H9 HA) | Pool ID | Ct (M) | Sample ID | Ct (M) | Ct (H5 HA) | Ct (H9 HA) | Pool ID | Ct (M) | Sample ID | Ct (M) | Ct (H5 HA) | Ct (H9 HA) | Pool ID | Ct (M) | Sample ID | Ct (M) | Ct (H5 HA) | Ct (H9 HA) |
| | | 10B1-091-0 | | | | | | 10B1-091-1 | | | | | | 10B1-091-2 | 22.67 | x | x | | | 10B1-091-3 | | x | x | | | 10B1-091-4 | | 35.14 | x |
| | | 10B1-092-0 | | | | | | 10B1-092-1 | | | | | | 10B1-092-2 | x | | | | | 10B1-092-3 | 30.83 | 39.61 | 33.05 | | | 10B1-092-4 | | 28.42 | 28.48 |
| 10B1-0 | x | 10B1-093-0 | | | | 10B1-1 | x | 10B1-093-1 | | | | 10B1-2 | 39.27 | 10B1-093-2 | x | | | 10B1-3 | | 10B1-093-3 | 35.47 | x | 37.09 | 10B1-4 | | 10B1-093-4 | 26.76 | x | 31.11 |
| | | 10B1-094-0 | | | | | | 10B1-094-1 | | | | | | 10B1-094-2 | x | | | | | 10B1-094-3 | 40.98 | | | | | 10B1-094-4 | 36.69 | 38.42 | x |
| | | 10B1-095-0 | | | | | | 10B1-095-1 | | | | | | 10B1-095-2 | x | | | | | 10B1-095-3 | 27.94 | x | 30.23 | | | 10B1-095-4 | | x | |
| | | | | | | | | 10B0-096-1 | | | | | | 10B0-096-2 | | | | | | 10B0-096-3 | x | | | | | 10B0-096-4 | 23.97 | 38.37 | 27.12 |
| | | | | | | | | 10B0-097-1 | | | | | | 10B0-097-2 | | | | | | 10B0-097-3 | 37.77 | x | 38.96 | | | 10B0-097-4 | 21.01 | 29.14 | 32.01 |
| - | | | | | | 10B0-1 | x | 10B0-098-1 | | | | 10B0-2 | x | 10B0-098-2 | | | | 10B0-3 | | 10B0-098-3 | x | | | 10B0-4 | | 10B0-098 | 38.23 | x | x |
| | | | | | | | | 10B0-099-1 | | | | | | 10B0-099-2 | | | | | | 10B0-099-3 | 25.83 | 27.71 | 37.69 | | | 10B0-099 | | | x |
| | | | | | | | | 10B0-100-1 | | | | | | 10B0-100-2 | | | | | | 10B0-100-3 | x | | | | | 10B0-100-4 | 27.12 | x | 29.16 |
| | | 11B1-101-0 | | | | | | 11B1-101-1 | | | | | | 11B1-101-2 | 38.08 | 39.01 | x | | | 11B1-101-3 | x | 39.86 | x | | | 11B1-101-4 | 24.11 | 32.41 | x |
| | | 11B1-102-0 | | | | | | 11B1-102 | x | | | | | ~~11B1-102-2~~ | | | | | | ~~11B1-102-3~~ | | | | | | ~~11B1-102-4~~ | | | |
| 11B1-0 | x | 11B1-103-0 | | | | 11B1-1 | x | 11B1-103-1 | | | | 11B1-2 | | 11B1-103-2 | 39.09 | x | x | 11B1-3 | | 11B1-103-3 | 31.61 | 32.73 | x | 11B1-4 | | 11B1-103 | | | x |
| | | 11B1-104-0 | | | | | | 11B1-104-1 | | | | | | 11B1-104-2 | 35.55 | 37.01 | x | | | 11B1-104-3 | x | x | | | | 11B1-104-4 | 24.21 | 29.19 | x |
| | | 11B1-105-0 | | | | | | 11B1-105-1 | | | | | | 11B1-105-2 | 33.85 | x | 44.81 | | | 11B1-105-3 | 30.34 | 38.20 | 31.84 | | | 11B1-105-4 | | 30.34 | |
| | | | | | | | | 11B0-106-1 | 44.11 | | | | | 11B0-106-2 | 37.80 | 38.84 | x | | | 11B0-106-3 | x | x | | | | 11B0-106-4 | 28.17 | 29.46 | x |
| | | | | | | | | 11B0-107-1 | 33.06 | 34.70 | x | | | 11B0-107-2 | 25.58 | 28.34 | x | | | 11B0-107-3 | | | x | | | 11B0-107 | | | 34.39 |
| - | | | | | | 11B0-1 | 38.48 | 11B0-108-1 | x | | | 11B0-2 | | 11B0-108-2 | x | | | 11B0-3 | | 11B0-108-3 | 28.31 | x | 30.12 | 11B0-4 | | 11B0-108-4 | | x | |
| | | | | | | | | 11B0-109-1 | 38.29 | x | x | | | 11B0-109-2 | x | | | | | 11B0-109-3 | x | | | | | 11B0-109-4 | x | | |
| | | | | | | | | 11B0-110-1 | x | | | | | 11B0-110-2 | 32.58 | 34.82 | x | | | 11B0-110-3 | | 39.80 | x | | | 11B0-110-4 | | x | 31.16 |
| | | 12B1-111-0 | | | | | | 12B1-111-1 | | | | | | 12B1-111-2 | x | | | | | 12B1-111-3 | 36.30 | 39.19 | x | | | 12B1-111-4 | 35.13 | x | x |
| | | 12B1-112-0 | | | | | | 12B1-112-1 | | | | | | 12B1-112-2 | x | | | | | 12B1-112-3 | 25.94 | x | 27.54 | | | 12B1-112-4 | | 39.53 | |
| 12B1-0 | x | 12B1-113-0 | | | | 12B1-1 | x | 12B1-113-1 | | | | 12B1-2 | | 12B1-113-2 | x | | | 12B1-3 | | 12B1-113-3 | 39.36 | x | 39.29 | 12B1-4 | | 12B1-113-4 | 38.78 | x | 39.67 |
| | | 12B1-114-0 | | | | | | 12B1-114-1 | | | | | | 12B1-114-2 | x | | | | | 12B1-114-3 | 24.52 | x | 26.93 | | | 12B1-114-4 | | x | |
| | | 12B1-115-0 | | | | | | 12B1-115-1 | | | | | | 12B1-115-2 | 27.96 | x | 30.62 | | | 12B1-115-3 | | x | | | | 12B1-115-4 | | x | |
| | | | | | | | | 12B0-116-1 | x | | | | | 12B0-116-2 | x | | | | | 12B0-116-3 | 29.79 | x | 32.55 | | | 12B0-116-4 | | 40.73 | |
| | | | | | | | | 12B0-117-1 | 29.02 | 30.33 | x | | | 12B0-117-2 | 26.39 | | x | | | 12B0-117-3 | | | x | | | 12B0-117 | | | x |
| - | | | | | | 12B0-1 | 31.53 | 12B0-118-1 | x | | | 12B0-2 | | 12B0-118-2 | x | | | 12B0-3 | | 12B0-118-3 | x | | | 12B0-4 | | 12B0-118-4 | 23.08 | x | 26.17 |
| | | | | | | | | 12B0-119-1 | x | | | | | 12B0-119-2 | x | | | | | 12B0-119-3 | 38.80 | x | 40.40 | | | 12B0-119-4 | 21.27 | 35.28 | 24.20 |
| | | | | | | | | 12B0-120-1 | x | | | | | 12B0-120-2 | x | | | | | 12B0-120-3 | 33.80 | x | 36.94 | | | 12B0-120-4 | 22.60 | x | 25.36 |
| | | 13B1-121-0 | | | | | | 13B1-121-1 | | | | | | 13B1-121-2 | | | | | | 13B1-121-3 | | | | | | 13B1-121-4 | 28.91 | x | 30.92 |
| | | 13B1-122-0 | | | | | | 13B1-122-1 | | | | | | 13B1-122-2 | | | | | | 13B1-122-3 | | | | | | 13B1-122-4 | 23.43 | x | 24.83 |
| 13B1-0 | x | 13B1-123-0 | | | | 13B1-1 | x | 13B1-123-1 | | | | 13B1-2 | x | 13B1-123-2 | | | | 13B1-3 | x | 13B1-123-3 | | | | 13B1-4 | 26.22 | 13B1-123-4 | x | | |
| | | 13B1-124-0 | | | | | | 13B1-124-1 | | | | | | 13B1-124-2 | | | | | | 13B1-124-3 | | | | | | 13B1-124-4 | 26.77 | x | 27.24 |
| | | 13B1-125-0 | | | | | | 13B1-125-1 | | | | | | 13B1-125-2 | | | | | | 13B1-125-3 | | | | | | 13B1-125-4 | 24.60 | x | 25.81 |
| | | | | | | | | 13B0-126-1 | | | | | | 13B0-126-2 | | | | | | 13B0-126-3 | x | | | | | 13B0-126 | 31.61 | x | 33.60 |
| | | | | | | | | 13B0-127-1 | | | | | | 13B0-127-2 | | | | | | 13B0-127-3 | x | | | | | 13B0-127-4 | x | | |
| - | | | | | | 13B0-1 | x | 13B0-128-1 | | | | 13B0-2 | x | 13B0-128-2 | | | | 13B0-3 | 27.00 | 13B0-128-3 | 23.11 | x | 38.73 | 13B0-4 | | 13B0-128-4 | | x | 26.25 |
| | | | | | | | | 13B0-129-1 | | | | | | 13B0-129-2 | | | | | | 13B0-129-3 | 23.28 | x | 38.68 | | | 13B0-129-4 | | x | 29.46 |
| | | | | | | | | 13B0-130-1 | | | | | | 13B0-130-2 | | | | | | 13B0-130-3 | x | | | | | 13B0-130-4 | x | | |
| | | 14B1-131-0 | | | | | | 14B1-131-1 | | | | | | 14B1-131-2 | | | | | | 14B1-131-3 | | | | | | 14B1-131-4 | | | |
| | | 14B1-132-0 | | | | | | 14B1-132-1 | | | | | | 14B1-132-2 | | | | | | 14B1-132-3 | | | | | | 14B1-132-4 | | | |
| 14B1-0 | x | 14B1-133-0 | | | | 14B1-1 | x | 14B1-133-1 | | | | 14B1-2 | x | 14B1-133-2 | | | | 14B1-3 | x | 14B1-133-3 | | | | 14B1-4 | x | 14B1-133-4 | | | |
| | | 14B1-134-0 | | | | | | 14B1-134-1 | | | | | | 14B1-134-2 | | | | | | 14B1-134-3 | | | | | | 14B1-134-4 | | | |
| | | 14B1-135-0 | | | | | | 14B1-135-1 | | | | | | 14B1-135-2 | | | | | | 14B1-135-3 | | | | | | 14B1-135-4 | | | |
| | | | | | | | | 14B0-136-1 | | | | | | 14B0-136-2 | | | | | | 14B0-136-3 | | | | | | 14B0-136-4 | | | |
| | | | | | | | | 14B0-137-1 | | | | | | 14B0-137-2 | | | | | | 14B0-137-3 | | | | | | 14B0-137-4 | | | |
| - | | | | | | 14B0-1 | x | 14B0-138-1 | | | | 14B0-2 | x | 14B0-138-2 | | | | 14B0-3 | x | 14B0-138-3 | | | | 14B0-4 | x | 14B0-138-4 | | | |
| | | | | | | | | 14B0-139-1 | | | | | | 14B0-139-2 | | | | | | 14B0-139-3 | | | | | | 14B0-139-4 | | | |
| | | | | | | | | 14B0-140-1 | | | | | | 14B0-140-2 | | | | | | 14B0-140-3 | | | | | | 14B0-140-4 | | | |

**T0**

| Pool ID | Ct (M) | Sample ID | Ct (M) | Ct (H5 HA) | Ct (H9 HA) |
|---|---|---|---|---|---|
| | | 15B1-141-0 | | | |
| | | 15B1-142-0 | | | |
| 15B1-0 | x | 15B1-143-0 | | | |
| | | 15B1-144-0 | | | |
| | | 15B1-145-0 | | | |
| | | - | | | |
| | | 16B1-151-0 | | | |
| | | 16B1-152-0 | | | |
| 16B1-0 | x | 16B1-153-0 | | | |
| | | 16B1-154-0 | | | |
| | | 16B1-155-0 | | | |
| | | - | | | |
| | | 17B1-161-0 | | | |
| | | 17B1-162-0 | | | |
| 17B1-0 | x | 17B1-163-0 | | | |
| | | 17B1-164-0 | | | |
| | | 17B1-165-0 | | | |
| | | - | | | |
| | | 18B1-171-0 | x | | |
| | | 18B1-172-0 | 38.41 | x | x |
| 18B1-0 | 31.53 | 18B1-173-0 | x | | |
| | | 18B1-174-0 | x | | |
| | | 18B1-175-0 | 43.63 | | |
| | | - | | | |
| | | 19B1-181-0 | x | | |
| | | 19B1-182-0 | 37.23 | x | x |
| 19B1-0 | 25.76 | 19B1-183-0 | 39.45 | x | x |
| | | 19B1-184-0 | 39.45 | x | x |
| | | 19B1-185-0 | 22.88 | x | 21.89 |
| | | - | | | |

**T1**

| Pool ID | Ct (M) | Sample ID | Ct (M) | Ct (H5 HA) | Ct (H9 HA) |
|---|---|---|---|---|---|
| | | 15B1-141-1 | | | |
| | | 15B1-142-1 | | | |
| 15B1-1 | x | 15B1-143-1 | | | |
| | | 15B1-144-1 | | | |
| | | 15B1-145-1 | | | |
| | | 15B0-146-1 | | | |
| | | 15B0-147-1 | | | |
| 15B0-1 | x | 15B0-148-1 | | | |
| | | 15B0-149-1 | | | |
| | | 15B0-150-1 | | | |
| | | 16B1-151-1 | | | |
| | | 16B1-152-1 | | | |
| 16B1-1 | x | 16B1-153-1 | | | |
| | | 16B1-154-1 | | | |
| | | 16B1-155-1 | | | |
| | | 16B0-156-1 | | | |
| | | 16B0-157-1 | | | |
| 16B0-1 | x | 16B0-158-1 | | | |
| | | 16B0-159-1 | | | |
| | | 16B0-160-1 | | | |
| | | 17B1-161-1 | | | |
| | | 17B1-162-1 | | | |
| 17B1-1 | x | 17B1-163-1 | | | |
| | | 17B1-164-1 | | | |
| | | 17B1-165-1 | | | |
| | | 17B0-166-1 | | | |
| | | 17B0-167-1 | | | |
| 17B0-1 | x | 17B0-168-1 | | | |
| | | 17B0-169-1 | | | |
| | | 17B0-170-1 | | | |
| | | 18B1-171-1 | | | |
| | | 18B1-172-1 | | | |
| 18B1-1 | x | 18B1-173-1 | | | |
| | | 18B1-174-1 | | | |
| | | 18B1-175-1 | | | |
| | | 18B0-176-1 | | | |
| | | 18B0-177-1 | | | |
| 18B0-1 | x | 18B0-178-1 | | | |
| | | 18B0-179-1 | | | |
| | | 18B0-180-1 | | | |
| | | 19B1-181-1 | 34.80 | x | 33.97 |
| | | 19B1-182-1 | 36.31 | x | 36.92 |
| 19B1-1 | 24.85 | 19B1-183-1 | 39.28 | x | x |
| | | 19B1-184-1 | 21.73 | x | 21.31 |
| | | 19B1-185-1 | 25.84 | x | 25.13 |
| | | 19B0-186-1 | | | |
| | | 19B0-187-1 | | | |
| 19B0-1 | x | 19B0-188-1 | | | |
| | | 19B0-189-1 | | | |
| | | 19B0-190-1 | | | |

**T2**

| Pool ID | Ct (M) | Sample ID | Ct (M) | Ct (H5 HA) | Ct (H9 HA) |
|---|---|---|---|---|---|
| | | 15B1-141-2 | | | |
| | | 15B1-142-2 | | | |
| 15B1-2 | x | 15B1-143-2 | | | |
| | | 15B1-144-2 | | | |
| | | 15B1-145-2 | | | |
| | | 15B0-146-2 | 25.91 | x | 27.38 |
| | | 15B0-147-2 | x | | |
| 15B0-2 | 30.49 | 15B0-148-2 | x | | |
| | | 15B0-149-2 | x | | |
| | | 15B0-150-2 | x | | |
| | | 16B1-151-2 | | | |
| | | 16B1-152-2 | | | |
| 16B1-2 | x | 16B1-153-2 | | | |
| | | 16B1-154-2 | | | |
| | | 16B1-155-2 | | | |
| | | 16B0-156-2 | | | |
| | | 16B0-157-2 | | | |
| 16B0-2 | x | 16B0-158-2 | | | |
| | | 16B0-159-2 | | | |
| | | 16B0-160-2 | | | |
| | | 17B1-161-2 | | | |
| | | 17B1-162-2 | | | |
| 17B1-2 | x | 17B1-163-2 | | | |
| | | 17B1-164-2 | | | |
| | | 17B1-165-2 | | | |
| | | 17B0-166-2 | | | |
| | | 17B0-167-2 | | | |
| 17B0-2 | x | 17B0-168-2 | | | |
| | | 17B0-169-2 | | | |
| | | 17B0-170-2 | | | |
| | | 18B1-171-2 | | | |
| | | 18B1-172-2 | | | |
| 18B1-2 | x | 18B1-173-2 | | | |
| | | 18B1-174-2 | | | |
| | | 18B1-175-2 | | | |
| | | 18B0-176-2 | | | |
| | | 18B0-177-2 | | | |
| 18B0-2 | x | 18B0-178-2 | | | |
| | | 18B0-179-2 | | | |
| | | 18B0-180-2 | | | |
| | | 19B1-181-2 | 31.77 | x | 31.27 |
| | | 19B1-182-2 | 27.86 | x | 27.70 |
| 19B1-2 | | 19B1-183-2 | 38.39 | x | 35.84 |
| | | 19B1-184-2 | 14.81 | x | |
| | | 19B1-185-2 | 25.75 | x | |
| | | 19B0-186-2 | 27.45 | x | 30.37 |
| | | 19B0-187-2 | 32.97 | x | 36.16 |
| 19B0-2 | | 19B0-188-2 | x | | |
| | | 19B0-189-2 | x | | |
| | | 19B0-190-2 | x | | |

**T3**

| Pool ID | Ct (M) | Sample ID | Ct (M) | Ct (H5 HA) | Ct (H9 HA) |
|---|---|---|---|---|---|
| | | 15B1-141-3 | x | | |
| | | 15B1-142-3 | x | | |
| 15B1-3 | | 15B1-143-3 | 28.44 | x | 30.69 |
| | | 15B1-144-3 | x | | |
| | | 15B1-145-3 | x | | |
| | | 15B0-146-3 | | x | |
| | | 15B0-147-3 | x | | |
| 15B0-3 | | 15B0-148-3 | x | | |
| | | 15B0-149-3 | x | | |
| | | 15B0-150-3 | x | | |
| | | 16B1-151-3 | 33.52 | x | 36.99 |
| | | 16B1-152-3 | x | | |
| 16B1-3 | 33.99 | 16B1-153-3 | 28.40 | x | 31.69 |
| | | 16B1-154-3 | 35.37 | x | 33.72 |
| | | 16B1-155-3 | 39.25 | x | x |
| | | 16B0-156-3 | 32.73 | x | 36.44 |
| | | 16B0-157-3 | 24.97 | x | 28.61 |
| 16B0-3 | 28.69 | 16B0-158-3 | 29.78 | x | 32.57 |
| | | 16B0-159-3 | 25.61 | x | 38.85 |
| | | 16B0-160-3 | 26.54 | x | 29.34 |
| | | 17B1-161-3 | x | | |
| | | 17B1-162-3 | x | | |
| 17B1-3 | 36.36 | 17B1-163-3 | x | | |
| | | 17B1-164-3 | 30.06 | x | 31.69 |
| | | 17B1-165-3 | 31.48 | x | 33.72 |
| | | 17B0-166-3 | x | | |
| | | 17B0-167-3 | 34.41 | x | x |
| 17B0-3 | 38.89 | 17B0-168-3 | x | | |
| | | 17B0-169-3 | x | | |
| | | 17B0-170-3 | x | | |
| | | 18B1-171-3 | x | | |
| | | 18B1-172-3 | 28.08 | x | 31.11 |
| 18B1-3 | 37.69 | 18B1-173-3 | 31.42 | x | 36.33 |
| | | 18B1-174-3 | 38.15 | x | x |
| | | 18B1-175-3 | x | | |
| | | 18B0-176-3 | 32.36 | x | 35.12 |
| | | 18B0-177-3 | 23.81 | x | 27.00 |
| 18B0-3 | 30.89 | 18B0-178-3 | 22.19 | x | 24.14 |
| | | 18B0-179-3 | 39.62 | x | x |
| | | 18B0-180-3 | 38.76 | x | 39.38 |
| | | 19B1-181-3 | | x | |
| | | 19B1-182-3 | | x | |
| 19B1-3 | | 19B1-183-3 | 25.39 | x | 24.37 |
| | | 19B1-184-3 | | x | |
| | | 19B1-185-3 | | x | |
| | | 19B0-186-3 | | x | |
| | | 19B0-187-3 | | x | 34.71 |
| 19B0-3 | | 19B0-188-3 | x | | |
| | | 19B0-189-3 | x | | |
| | | 19B0-190-3 | x | | |

**T4**

| Pool ID | Ct (M) | Sample ID | Ct (M) | Ct (H5 HA) | Ct (H9 HA) |
|---|---|---|---|---|---|
| | | 15B1-141-4 | x | | |
| | | 15B1-142-4 | x | | |
| 15B1-4 | | 15B1-143-4 | | 40.15 | |
| | | 15B1-144-4 | 32.44 | 30.07 | x |
| | | 15B1-145-4 | 27.03 | x | 28.82 |
| | | 15B0-146-4 | | x | |
| | | 15B0-147 | 27.62 | x | 28.58 |
| 15B0-4 | | 15B0-148-4 | 24.77 | x | 24.65 |
| | | 15B0-149-4 | x | | |
| | | 15B0-150-4 | 25.19 | 34.98 | 25.37 |
| | | 16B1-151-4 | 35.02 | x | 42.05 |
| | | 16B1-152-4 | x | | |
| 16B1-4 | | 16B1-153-4 | | x | |
| | | 16B1-154-4 | x | | x |
| | | 16B1-155-4 | x | | |
| | | 16B0-156-4 | | x | 29.04 |
| | | 16B0-157-4 | | x | |
| 16B0-4 | | 16B0-158-4 | | x | |
| | | 16B0-159-4 | | x | 25.33 |
| | | 16B0-160-4 | | x | |
| | | 17B1-161-4 | 33.02 | x | 38.17 |
| | | 17B1-162-4 | 28.01 | x | 26.58 |
| 17B1-4 | | 17B1-163-4 | 44.96 | | |
| | | 17B1-164-4 | | x | |
| | | 17B1-165-4 | | x | 22.50 |
| | | 17B0-166-4 | x | | |
| | | 17B0-167-4 | 21.21 | x | 22.30 |
| 17B0-4 | | 17B0-168-4 | x | | |
| | | 17B0-169-4 | 28.82 | x | 29.36 |
| | | 17B0-170-4 | 30.94 | x | 29.57 |
| | | 18B1-171-4 | 36.89 | x | x |
| | | 18B1-172-4 | | x | |
| 18B1-4 | | 18B1-173-4 | | x | 35.47 |
| | | 18B1-174-4 | 26.44 | x | 27.28 |
| | | 18B1-175-4 | 42.61 | | |
| | | 18B0-176-4 | | x | 22.72 |
| | | 18B0-177-4 | | x | |
| 18B0-4 | | 18B0-178-4 | | x | |
| | | 18B0-179-4 | 26.34 | x | 26.66 |
| | | 18B0-180-4 | 24.49 | x | 25.50 |
| | | 19B1-181-4 | | x | |
| | | 19B1-182-4 | | x | |
| 19B1-4 | | 19B1-183-4 | | x | |
| | | 19B1-184-4 | | 37.65 | |
| | | 19B1-185-4 | | x | |
| | | 19B0-186-4 | | x | |
| | | 19B0-187-4 | | x | 35.18 |
| 19B0-4 | | 19B0-188-4 | x | | |
| | | 19B0-189-4 | x | | |
| | | 19B0-190-4 | x | | |

| T0 | | | | | | T1 | | | | | | T2 | | | | | | T3 | | | | | | T4 | | | | | |
|---|---|---|---|---|---|---|---|---|---|---|---|---|---|---|---|---|---|---|---|---|---|---|---|---|---|---|---|---|---|
| Pool ID | Ct (M) | Sample ID | Ct (M) | Ct (H5 HA) | Ct (H9 HA) | Pool ID | Ct (M) | Sample ID | Ct (M) | Ct (H5 HA) | Ct (H9 HA) | Pool ID | Ct (M) | Sample ID | Ct (M) | Ct (H5 HA) | Ct (H9 HA) | Pool ID | Ct (M) | Sample ID | Ct (M) | Ct (H5 HA) | Ct (H9 HA) | Pool ID | Ct (M) | Sample ID | Ct (M) | Ct (H5 HA) | Ct (H9 HA) |
| | | 20B1-191-0 | | | | | | 20B1-191-1 | | | | | | 20B1-191-2 | | | | | | 20B1-191-3 | x | | | | | 20B1-191-4 | 40.39 | | |
| | | 20B1-192-0 | | | | | | 20B1-192-1 | | | | | | 20B1-192-2 | | | | | | 20B1-192-3 | x | x | 41.37 | | | 20B1-192-4 | x | | |
| 20B1-0 | x | 20B1-193-0 | | | | 20B1-1 | x | 20B1-193-1 | | | | 20B1-2 | x | 20B1-193-2 | | | | 20B1-3 | | 20B1-193-3 | x | | | 20B1-4 | | 20B1-193-4 | x | | |
| | | 20B1-194-0 | | | | | | 20B1-194-1 | | | | | | 20B1-194-2 | | | | | | 20B1-194-3 | x | | | | | 20B1-194-4 | x | | |
| | | 20B1-195-0 | | | | | | 20B1-195-1 | | | | | | 20B1-195-2 | | | | | | 20B1-195-3 | 39.61 | x | 38.09 | | | 20B1-195-4 | x | | |
| | | | | | | | | 20B0-196-1 | | | | | | 20B0-196-2 | x | | | | | 20B0-196-3 | x | | | | | 20B0-196-4 | x | | |
| | | | | | | | | 20B0-197-1 | | | | | | 20B0-197-2 | 24.83 | x | 39.51 | | | 20B0-197-3 | | x | 33.54 | | | 20B0-197-4 | | x | x |
| | | - | | | | 20B0-1 | x | 20B0-198-1 | | | | 20B0-2 | 28.62 | 20B0-198-2 | x | | | 20B0-3 | | 20B0-198-3 | x | | | 20B0-4 | | 20B0-198-4 | x | | |
| | | | | | | | | 20B0-199-1 | | | | | | 20B0-199-2 | x | | | | | 20B0-199-3 | x | | | | | 20B0-199-4 | x | | |
| | | | | | | | | 20B0-200-1 | | | | | | 20B0-200-2 | 36.55 | x | 37.12 | | | 20B0-200-3 | 25.88 | x | 29.73 | | | 20B0-200-4 | | x | |
| | | 21B1-201-0 | | | | | | 21B1-201-1 | | | | | | 21B1-201-2 | x | | | | | 21B1-201-3 | 32.17 | x | 33.37 | | | 21B1-201-4 | | x | 35.77 |
| | | 21B1-202-0 | | | | | | 21B1-202-1 | | | | | | 21B1-202-2 | 36.91 | x | x | | | 21B1-202-3 | 42.83 | | | | | 21B1-202-4 | 37.28 | x | x |
| 21B1-0 | x | 21B1-203-0 | | | | 21B1-1 | x | 21B1-203-1 | | | | 21B1-2 | | 21B1-203-2 | x | | | 21B1-3 | | 21B1-203-3 | 24.25 | x | 24.57 | 21B1-4 | | 21B1-203-4 | | x | |
| | | 21B1-204-0 | | | | | | 21B1-204-1 | | | | | | 21B1-204-2 | 35.17 | x | 38.39 | | | 21B1-204-3 | 26.39 | x | 27.75 | | | 21B1-204-4 | | x | |
| | | 21B1-205-0 | | | | | | 21B1-205-1 | | | | | | 21B1-205-2 | 32.92 | x | 34.66 | | | 21B1-205-3 | | x | 28.60 | | | 21B1-205-4 | | x | |
| | | | | | | | | 21B0-206-1 | x | | | | | 21B0-206-2 | x | | | | | 21B0-206-3 | 20.42 | x | 21.47 | | | 21B0-206-4 | | x | |
| | | | | | | | | 21B0-207-1 | 38.28 | x | x | | | 21B0-207-2 | 34.27 | x | 34.89 | | | 21B0-207-3 | 27.41 | x | 27.02 | | | 21B0-207-4 | | x | |
| | | - | | | | 21B0-1 | 32.49 | 21B0-208-1 | 30.26 | x | 31.51 | 21B0-2 | | 21B0-208-2 | 19.16 | x | | 21B0-3 | | 21B0-208-3 | | x | | 21B0-4 | | 21B0-208-4 | | x | |
| | | | | | | | | 21B0-209-1 | 40.80 | | | | | 21B0-209-2 | 31.35 | x | 33.34 | | | 21B0-209-3 | | x | 22.81 | | | 21B0-209-4 | | x | |
| | | | | | | | | 21B0-210-1 | x | | | | | 21B0-210-2 | 42.82 | | | | | 21B0-210-3 | 23.57 | x | 24.08 | | | 21B0-210-4 | | x | |
| | | 22B1-211-0 | x | | | | | 22B1-211-1 | 24.85 | x | 25.49 | | | 22B1-211-2 | 25.95 | x | | | | 22B1-211-3 | | x | | | | 22B1-211-4 | | x | |
| | | 22B1-212-0 | x | | | | | 22B1-212-1 | 24.09 | x | 25.53 | | | 22B1-212-2 | 24.93 | x | | | | 22B1-212-3 | | x | | | | 22B1-212-4 | | x | |
| 22B1-0 | 27.83 | 22B1-213-0 | 41.27 | | | 22B1-1 | 25.30 | 22B1-213-1 | 25.51 | x | 27.07 | 22B1-2 | | 22B1-213-2 | 25.67 | x | | 22B1-3 | | 22B1-213-3 | | x | | 22B1-4 | | 22B1-213-4 | | x | |
| | | 22B1-214-0 | 24.68 | x | 26.49 | | | 22B1-214-1 | 32.32 | x | 30.10 | | | 22B1-214-2 | x | | x | | | 22B1-214-3 | x | | x | | | 22B1-214-4 | 24.90 | x | 27.19 |
| | | 22B1-215-0 | 35.49 | x | x | | | 22B1-215-1 | 41.90 | | | | | 22B1-215-2 | x | | | | | 22B1-215-3 | x | | | | | 22B1-215-4 | 36.00 | x | 39.78 |
| | | | | | | | | 22B0-216-1 | 43.76 | | | | | 22B0-216-2 | x | | | | | 22B0-216-3 | 23.80 | x | 25.23 | | | 22B0-216-4 | | x | |
| | | | | | | | | 22B0-217-1 | 37.90 | 41.11 | x | | | 22B0-217-2 | x | x | | | | 22B0-217-3 | 23.56 | x | 25.13 | | | 22B0-217-4 | | x | |
| | | - | | | | 22B0-1 | 36.95 | 22B0-218-1 | 44.51 | | | 22B0-2 | | 22B0-218-2 | x | | | 22B0-3 | | 22B0-218-3 | 23.19 | x | 24.24 | 22B0-4 | | 22B0-218-4 | | x | |
| | | | | | | | | 22B0-219-1 | x | | | | | 22B0-219-2 | x | | | | | 22B0-219-3 | x | | | | | 22B0-219-4 | 20.62 | x | 25.22 |
| | | | | | | | | 22B0-220-1 | x | | | | | 22B0-220-2 | x | | | | | 22B0-220-3 | 30.76 | x | 31.49 | | | 22B0-220-4 | | x | |
| | | 23B1-221-0 | 37.25 | x | x | | | 23B1-221-1 | x | | | | | 23B1-221-2 | 39.76 | x | x | | | 23B1-221-3 | 26.29 | x | 27.07 | | | 23B1-221-4 | | x | |
| | | 23B1-222-0 | 38.63 | x | 37.83 | | | 23B1-222-1 | 43.60 | | | | | 23B1-222-2 | x | | | | | 23B1-222-3 | 28.79 | 39.04 | 30.30 | | | 23B1-222-4 | | x | |
| 23B1-0 | 36.43 | 23B1-223-0 | x | | | 23B1-1 | 36.63 | 23B1-223-1 | 38.62 | x | x | 23B1-2 | | 23B1-223-2 | x | | | 23B1-3 | | 23B1-223-3 | 30.06 | x | 37.77 | 23B1-4 | | 23B1-223-4 | | x | 23.27 |
| | | 23B1-224-0 | 42.32 | | | | | 23B1-224-1 | x | | | | | 23B1-224-2 | x | | | | | 23B1-224-3 | 25.66 | x | 27.57 | | | 23B1-224-4 | | x | |
| | | 23B1-225-0 | 43.14 | | | | | 23B1-225-1 | x | | | | | 23B1-225-2 | x | | | | | 23B1-225-3 | x | | | | | 23B1-225-4 | 20.62 | x | 20.27 |
| | | | | | | | | 23B0-226-1 | | | | | | 23B0-226-2 | x | | | | | 23B0-226-3 | x | | | | | 23B0-226-4 | 20.91 | x | 28.26 |
| | | | | | | | | 23B0-227-1 | | | | | | 23B0-227-2 | x | | | | | 23B0-227-3 | x | | | | | 23B0-227-4 | 25.71 | x | 27.95 |
| | | - | | | | 23B0-1 | x | 23B0-228-1 | | | | 23B0-2 | | 23B0-228-2 | x | | | 23B0-3 | | 23B0-228-3 | 25.32 | x | 26.99 | 23B0-4 | | 23B0-228-4 | | x | |
| | | | | | | | | 23B0-229-1 | | | | | | 23B0-229-2 | 32.96 | x | 31.21 | | | 23B0-229-3 | x | | | | | 23B0-229-4 | 36.50 | x | 38.46 |
| | | | | | | | | 23B0-230-1 | | | | | | 23B0-230-2 | x | | | | | 23B0-230-3 | x | | | | | 23B0-230-4 | 21.30 | x | 22.26 |
| | | 24B1-231-0 | x | | | | | 24B1-231-1 | 38.23 | x | x | | | 24B1-231-2 | x | | | | | 24B1-231-3 | 23.92 | x | 22.39 | | | 24B1-231-4 | | x | |
| | | 24B1-232-0 | 39.00 | x | x | | | 24B1-232-1 | x | | | | | 24B1-232-2 | x | | | | | 24B1-232-3 | x | | | | | 24B1-232-4 | 26.18 | x | 27.00 |
| 24B1-0 | 31.02 | 24B1-233-0 | x | | | 24B1-1 | 38.34 | 24B1-233-1 | x | | | 24B1-2 | | 24B1-233-2 | x | | | 24B1-3 | 31.43 | 24B1-233-3 | 37.99 | x | x | 24B1-4 | | 24B1-233-4 | 37.28 | x | x |
| | | 24B1-234-0 | 29.24 | x | 31.23 | | | 24B1-234-1 | x | | x | | | 24B1-234-2 | x | | | | | 24B1-234-3 | 36.32 | x | x | | | 24B1-234-4 | x | | |
| | | 24B1-235-0 | x | | | | | 24B1-235-1 | x | | | | | 24B1-235-2 | x | | | | | 24B1-235-3 | 43.53 | | | | | 24B1-235-4 | 36.56 | x | 34.23 |
| | | | | | | | | 24B0-236-1 | | | | | | 24B0-236-2 | x | | | | | 24B0-236-3 | 35.06 | x | 44.45 | | | 24B0-236-4 | 24.08 | x | 25.13 |
| | | | | | | | | 24B0-237-1 | | | | | | 24B0-237-2 | x | | | | | 24B0-237-3 | 36.90 | x | 41.17 | | | 24B0-237-4 | 27.46 | x | 27.90 |
| | | - | | | | 24B0-1 | x | 24B0-238-1 | | | | 24B0-2 | | 24B0-238-2 | 44.1 | | | 24B0-3 | 33.47 | 24B0-238-3 | 23.22 | x | 30.20 | 24B0-4 | | 24B0-238-4 | | x | |
| | | | | | | | | 24B0-239-1 | | | | | | 24B0-239-2 | x | | | | | 24B0-239-3 | 38.69 | x | 33.45 | | | 24B0-239-4 | 36.12 | x | 37.65 |
| | | | | | | | | 24B0-240-1 | | | | | | 24B0-240-2 | x | | | | | 24B0-240-3 | 29.40 | x | 30.81 | | | 24B0-240-4 | | x | |

**T0**

| Pool ID | Ct (M) | Sample ID | Ct (M) | Ct (H5 HA) | Ct (H9 HA) |
|---|---|---|---|---|---|
| | | 25B1-241-0 | x | | |
| | | 25B1-242-0 | x | | |
| 25B1-0 | 34.30 | 25B1-243-0 | 43.97 | | |
| | | 25B1-244-0 | 38.40 | x | x |
| | | 25B1-245-0 | 43.57 | | |
| – | | | | | |
| | | 26B1-251-0 | | | |
| | | 26B1-252-0 | | | |
| 26B1-0 | x | 26B1-253-0 | | | |
| | | 26B1-254-0 | | | |
| | | 26B1-255-0 | | | |
| – | | | | | |
| | | 27B1-261-0 | x | | |
| | | 27B1-262-0 | 41.89 | | |
| 27B1-0 | 33.58 | 27B1-263-0 | x | | |
| | | 27B1-264-0 | x | | |
| | | 27B1-265-0 | 37.34 | x | x |
| – | | | | | |
| | | 28B1-271-0 | x | | |
| | | 28B1-272-0 | 37.62 | x | x |
| 28B1-0 | 32.8 | 28B1-273-0 | x | | |
| | | 28B1-274-0 | x | | |
| | | 28B1-275-0 | x | | |
| – | | | | | |
| | | 29B1-281-0 | 43.20 | | |
| | | 29B1-282-0 | 44.92 | | |
| 29B1-0 | 32.70 | 29B1-283-0 | 43.19 | | |
| | | 29B1-284-0 | x | | |
| | | 29B1-285-0 | 38.69 | x | 39.35 |
| – | | | | | |

**T1**

| Pool ID | Ct (M) | Sample ID | Ct (M) | Ct (H5 HA) | Ct (H9 HA) |
|---|---|---|---|---|---|
| | | 25B1-241-1 | x | | |
| | | 25B1-242-1 | x | | |
| 25B1-1 | 35.43 | 25B1-243-1 | x | | |
| | | 25B1-244-1 | x | | |
| | | 25B1-245-1 | x | | |
| | | 25B0-246-1 | x | | |
| | | 25B0-247-1 | x | | |
| 25B0-1 | 37.62 | 25B0-248-1 | 44.35 | | |
| | | 25B0-249-1 | x | | |
| | | 25B0-250-1 | 30.94 | x | x |
| | | 26B1-251-1 | x | | |
| | | 26B1-252-1 | 36.10 | x | 34.13 |
| 26B1-1 | 36.44 | 26B1-253-1 | x | | |
| | | 26B1-254-1 | x | | |
| | | 26B1-255-1 | x | | |
| | | 26B0-256-1 | 22.35 | x | 21.37 |
| | | 26B0-257-1 | x | | |
| 26B0-1 | 24.02 | 26B0-258-1 | 21.27 | x | 19.33 |
| | | 26B0-259-1 | 40.85 | | |
| | | 26B0-260-1 | 24.16 | x | 22.73 |
| | | 27B1-261-1 | x | | |
| | | 27B1-262-1 | x | | |
| 27B1-1 | 31.50 | 27B1-263-1 | x | | |
| | | 27B1-264-1 | x | | |
| | | 27B1-265-1 | 41.08 | | |
| | | 27B0-266-1 | 37.56 | x | x |
| | | 27B0-267-1 | x | | |
| 27B0-1 | 29.93 | 27B0-268-1 | 38.49 | x | x |
| | | 27B0-269-1 | x | | |
| | | 27B0-270-1 | x | | |
| | | 28B1-271-1 | 40.83 | | |
| | | 28B1-272-1 | x | | |
| 28B1-1 | 31.26 | 28B1-273-1 | x | | |
| | | 28B1-274-1 | 41.49 | | |
| | | 28B1-275-1 | 38.61 | 35.04 | x |
| | | 28B0-276-1 | 44.84 | | |
| | | 28B0-277-1 | 37.13 | x | x |
| 28B0-1 | 32.31 | 28B0-278-1 | 38.24 | x | x |
| | | 28B0-279-1 | 40.86 | | |
| | | 28B0-280-1 | x | | |
| | | 29B1-281-1 | 34.31 | x | 34.29 |
| | | 29B1-282-1 | 40.19 | | |
| 29B1-1 | 32.94 | 29B1-283-1 | 43.71 | | |
| | | 29B1-284-1 | 34.11 | x | 34.74 |
| | | 29B1-285-1 | 37.38 | x | 38.90 |
| | | 29B0-286-1 | 32.74 | x | 31.98 |
| | | 29B0-287-1 | 32.84 | x | 32.18 |
| 29B0-1 | 22.07 | 29B0-288-1 | 33.47 | x | 32.49 |
| | | 29B0-289-1 | 18.77 | x | 17.87 |
| | | 29B0-290-1 | x | | |

**T2**

| Pool ID | Ct (M) | Sample ID | Ct (M) | Ct (H5 HA) | Ct (H9 HA) |
|---|---|---|---|---|---|
| | | 25B1-241-2 | 41.81 | | |
| | | 25B1-242-2 | x | | |
| 25B1-2 | | 25B1-243-2 | x | | |
| | | 25B1-244-2 | x | | |
| | | 25B1-245-2 | x | | |
| | | 25B0-246-2 | 38.33 | x | 40.02 |
| | | 25B0-247-2 | 34.84 | x | 41.52 |
| 25B0-2 | | 25B0-248-2 | x | | |
| | | 25B0-249-2 | x | | |
| | | 25B0-250-2 | x | x | x |
| | | 26B1-251-2 | x | | |
| | | 26B1-252-2 | 32.9 | x | 33.77 |
| 26B1-2 | | 26B1-253-2 | x | | |
| | | 26B1-254-2 | x | | |
| | | 26B1-255-2 | x | | |
| | | 26B0-256-2 | 24.1 | x | |
| | | 26B0-257-2 | x | | |
| 26B0-2 | | 26B0-258-2 | x | | x |
| | | 26B0-259-2 | x | | |
| | | 26B0-260-2 | x | | x |
| | | 27B1-261-2 | x | | |
| | | 27B1-262-2 | 32.77 | x | 34.70 |
| 27B1-2 | | 27B1-263-2 | 34.03 | x | 35.96 |
| | | 27B1-264-2 | 33.45 | x | 37.60 |
| | | 27B1-265-2 | x | | |
| | | 27B0-266-2 | x | | |
| | | 27B0-267-2 | x | | |
| 27B0-2 | | 27B0-268-2 | x | | |
| | | 27B0-269-2 | x | | |
| | | 27B0-270-2 | x | | |
| | | 28B1-271-2 | 42.61 | | |
| | | 28B1-272-2 | x | | |
| 28B1-2 | | 28B1-273-2 | 32.01 | x | 33.71 |
| | | 28B1-274-2 | 37.60 | x | 40.40 |
| | | 28B1-275-2 | x | x | |
| | | 28B0-276-2 | x | | |
| | | 28B0-277 | x | | |
| 28B0-2 | | 28B0-278-2 | x | | |
| | | 28B0-279-2 | x | | |
| | | 28B0-280-2 | x | | |
| | | 29B1-281-2 | x | x | x |
| | | 29B1-282-2 | x | | |
| 29B1-2 | | 29B1-283-2 | 43.46 | | |
| | | 29B1-284-2 | x | | x |
| | | 29B1-285-2 | 31.96 | x | 30.24 |
| | | 29B0-286-2 | 29.26 | x | |
| | | 29B0-287-2 | 36.76 | | 35.22 |
| 29B0-2 | | 29B0-288-2 | 31.58 | x | |
| | | 29B0-289-2 | 21.72 | x | |
| | | 29B0-290-2 | x | | |

**T3**

| Pool ID | Ct (M) | Sample ID | Ct (M) | Ct (H5 HA) | Ct (H9 HA) |
|---|---|---|---|---|---|
| | | 25B1-241-3 | 21.82 | x | 23.27 |
| | | 25B1-242-3 | x | | |
| 25B1-3 | | 25B1-243-3 | x | | |
| | | 25B1-244-3 | x | | |
| | | 25B1-245-3 | x | | |
| | | 25B0-246-3 | 22.18 | x | 23.97 |
| | | 25B0-247-3 | 22.13 | x | 23.73 |
| 25B0-3 | | 25B0-248-3 | 23.74 | x | 25.79 |
| | | 25B0-249-3 | x | | |
| | | 25B0-250-3 | 25.69 | x | 27.67 |
| | | 26B1-251-3 | 39.11 | 34.13 | 33.87 |
| | | 26B1-252-3 | | 42.79 | 23.33 |
| 26B1-3 | | 26B1-253-3 | 24.80 | x | 23.21 |
| | | 26B1-254-3 | x | | |
| | | 26B1-255-3 | 30.60 | x | 29.31 |
| | | 26B0-256-3 | | x | |
| | | 26B0-257-3 | 22.06 | x | 20.87 |
| 26B0-3 | | 26B0-258-3 | 22.10 | x | 19.97 |
| | | 26B0-259-3 | 23.25 | x | 21.69 |
| | | 26B0-260-3 | 22.49 | x | 20.16 |
| | | 27B1-261-3 | x | | |
| | | 27B1-262-3 | | x | 34.78 |
| 27B1-3 | | 27B1-263 | 25.67 | x | 27.54 |
| | | 27B1-264-3 | 40.27 | | 38.47 |
| | | 27B1-265-3 | x | | |
| | | 27B0-266-3 | 32.63 | x | 35.85 |
| | | 27B0-267-3 | 34.02 | x | 36.01 |
| 27B0-3 | | 27B0-268-3 | 36.39 | x | 38.45 |
| | | 27B0-269-3 | 39.64 | 40.04 | 44.55 |
| | | 27B0-270 | x | | |
| | | 28B1-271-3 | 27.28 | x | 29.22 |
| | | 28B1-272-3 | 26.40 | x | 28.87 |
| 28B1-3 | | 28B1-273-3 | | 43.68 | 31.16 |
| | | 28B1-274-3 | 22.39 | 44.18 | 24.89 |
| | | 28B1-275-3 | 20.96 | x | 23.84 |
| | | 28B0-276-3 | x | | |
| | | ~~28B0-277-3~~ | | | |
| 28B0-3 | | 28B0-278-3 | 33.13 | 42.76 | 36.05 |
| | | 28B0-279-3 | 35.39 | x | 37.51 |
| | | 28B0-280-3 | 30.25 | x | 32.24 |
| | | 29B1-281-3 | 33.99 | x | 34.11 |
| | | 29B1-282-3 | x | | |
| 29B1-3 | | 29B1-283-3 | 30.69 | x | 30.37 |
| | | 29B1-284-3 | 36.92 | | 37.56 |
| | | 29B1-285-3 | | x | |
| | | 29B0-286-3 | | x | |
| | | 29B0-287-3 | 35.09 | | 35.22 |
| 29B0-3 | | 29B0-288-3 | | x | |
| | | 29B0-289-3 | | x | |
| | | 29B0-290-3 | 20.84 | x | 21.30 |

**T4**

| Pool ID | Ct (M) | Sample ID | Ct (M) | Ct (H5 HA) | Ct (H9 HA) |
|---|---|---|---|---|---|
| | | 25B1-241-4 | | x | |
| | | 25B1-242-4 | 31.89 | x | 34.47 |
| 25B1-4 | | 25B1-243-4 | x | | |
| | | 25B1-244-4 | 22.78 | x | 25.53 |
| | | 25B1-245-4 | 44.78 | | |
| | | 25B0-246-4 | | x | |
| | | 25B0-247-4 | | x | |
| 25B0-4 | | 25B0-248-4 | | x | |
| | | 25B0-249-4 | 19.32 | x | 27.57 |
| | | 25B0-250-4 | | x | |
| | | 26B1-251-4 | 23.78 | x | 22.02 |
| | | 26B1-252-4 | | x | 20.75 |
| 26B1-4 | | 26B1-253-4 | | x | |
| | | 26B1-254-4 | 37.34 | x | 35.08 |
| | | 26B1-255-4 | | x | |
| | | 26B0-256-4 | | x | |
| | | 26B0-257-4 | | 39.53 | |
| 26B0-4 | | 26B0-258-4 | | x | |
| | | 26B0-259-4 | | x | |
| | | 26B0-260-4 | | x | |
| | | 27B1-261-4 | 30.37 | x | 32.05 |
| | | 27B1-262-4 | | x | 27.99 |
| 27B1-4 | | ~~27B1-263-4~~ | | | |
| | | 27B1-264-4 | 37.19 | x | 42.58 |
| | | 27B1-265-4 | 41.24 | | |
| | | 27B0-266-4 | | x | 24.16 |
| | | 27B0-267-4 | 21.74 | x | 32.18 |
| 27B0-4 | | 27B0-268 | 35.34 | x | 34.45 |
| | | 27B0-269-4 | 25.02 | x | 29.97 |
| | | ~~27B0-270-4~~ | | | |
| | | 28B1-271-4 | | x | |
| | | 28B1-272-4 | | x | |
| 28B1-4 | | 28B1-273-4 | | x | |
| | | 28B1-274-4 | | x | |
| | | 28B1-275-4 | | x | |
| | | 28B0-276-4 | 26.64 | x | 30.31 |
| | | ~~28B0-277-4~~ | | | |
| 28B0-4 | | 28B0-278-4 | 30.43 | x | 35.75 |
| | | 28B0-279-4 | 22.64 | x | 26.45 |
| | | 28B0-280-4 | | x | |
| | | 29B1-281-4 | 35.51 | x | 34.01 |
| | | 29B1-282-4 | 34.14 | x | 32.56 |
| 29B1-4 | | 29B1-283-4 | | x | |
| | | 29B1-284-4 | 27.59 | x | 24.62 |
| | | 29B1-285-4 | | x | |
| | | 29B0-286-4 | | x | |
| | | 29B0-287-4 | 21.56 | x | 19.56 |
| 29B0-4 | | 29B0-288-4 | | x | |
| | | 29B0-289-4 | | x | |
| | | 29B0-290-4 | | x | |

| T0 | | | | | |
|---|---|---|---|---|---|
| Pool ID | Ct (M) | Sample ID | Ct (M) | Ct (H5 HA) | Ct (H9 HA) |
| | | 30B1-291-0 | x | | |
| | | 30B1-292-0 | 41.99 | | |
| 30B1-0 | 33.83 | 30B1-293-0 | x | | |
| | | 30B1-294-0 | 44.09 | | |
| | | 30B1-295-0 | 38.62 | x | x |
| | | - | | | |
| | | 31B1-301-0 | x | | |
| | | 31B1-302-0 | 44.52 | | |
| 31B1-0 | 33.53 | 31B1-303-0 | 37.16 | x | x |
| | | 31B1-304-0 | x | | |
| | | 31B1-305-0 | x | | |
| | | - | | | |
| | | 32B1-311-0 | x | | |
| | | 32B1-312-0 | x | | |
| 32B1-0 | 32.76 | 32B1-313-0 | x | | |
| | | 32B1-314-0 | x | | |
| | | 32B1-315-0 | 41.89 | | |
| | | - | | | |
| | | 33B1-321-0 | 38.15 | x | x |
| | | 33B1-322-0 | x | | |
| 33B1-0 | 32.36 | 33B1-323-0 | 37.87 | x | x |
| | | 33B1-324-0 | x | | |
| | | 33B1-325-0 | 36.21 | x | x |
| | | - | | | |
| | | 34B1-331-0 | x | | |
| | | 34B1-332-0 | 16.92 | x | x |
| 34B1-0 | 38.37 | 34B1-333-0 | x | | |
| | | 34B1-334-0 | x | | |
| | | 34B1-335-0 | x | | |
| | | - | | | |

| T1 | | | | | |
|---|---|---|---|---|---|
| Pool ID | Ct (M) | Sample ID | Ct (M) | Ct (H5 HA) | Ct (H9 HA) |
| | | 30B1-291-1 | x | | |
| | | 30B1-292-1 | x | | |
| 30B1-1 | 33.42 | 30B1-293-1 | 38.68 | x | x |
| | | 30B1-294-1 | x | | |
| | | 30B1-295-1 | x | | |
| | | 30B0-296-1 | x | | |
| | | 30B0-297-1 | x | | |
| 30B0-1 | 33.35 | 30B0-298-1 | 38.44 | x | x |
| | | 30B0-299-1 | 37.34 | x | x |
| | | 30B0-300-1 | x | | |
| | | 31B1-301-1 | x | | |
| | | 31B1-302-1 | x | | |
| 31B1-1 | 32.76 | 31B1-303-1 | 39.82 | x | x |
| | | 31B1-304-1 | x | | |
| | | 31B1-305-1 | x | | |
| | | 31B0-306-1 | x | | |
| | | 31B0-307-1 | 39.46 | x | x |
| 31B0-1 | 32.20 | 31B0-308-1 | x | | |
| | | 31B0-309-1 | x | | |
| | | 31B0-310-1 | x | | |
| | | 32B1-311-1 | x | | |
| | | 32B1-312-1 | x | | |
| 32B1-1 | 32.44 | 32B1-313-1 | x | | |
| | | 32B1-314-1 | x | | |
| | | 32B1-315-1 | x | | |
| | | 32B0-316-1 | x | | |
| | | 32B0-317-1 | x | | |
| 32B0-1 | 33.46 | 32B0-318-1 | x | | |
| | | 32B0-319-1 | x | | |
| | | 32B0-320-1 | x | | |
| | | 33B1-321-1 | x | | |
| | | 33B1-322 | 37.13 | x | x |
| 33B1-1 | 32.41 | 33B1-323-1 | x | | |
| | | 33B1-324-1 | 36.19 | x | x |
| | | 33B1-325-1 | x | | |
| | | 33B0-326-1 | x | | |
| | | 33B0-327-1 | x | | |
| 33B0-1 | 32.62 | 33B0-328-1 | 38.79 | x | x |
| | | 33B0-329-1 | x | | |
| | | 33B0-330-1 | x | | |
| | | 34B1-331-1 | x | | |
| | | 34B1-332-1 | x | | |
| 34B1-1 | 37.58 | 34B1-333-1 | 37.18 | x | x |
| | | 34B1-334-1 | x | | |
| | | 34B1-335-1 | x | | |
| | | 34B0-336-1 | | | |
| | | 34B0-337-1 | | | |
| 34B0-1 | x | 34B0-338-1 | | | |
| | | 34B0-339-1 | | | |
| | | 34B0-340-1 | | | |

| T2 | | | | | |
|---|---|---|---|---|---|
| Pool ID | Ct (M) | Sample ID | Ct (M) | Ct (H5 HA) | Ct (H9 HA) |
| | | 30B1-291-2 | x | | |
| | | 30B1-292-2 | 30.65 | x | 31.79 |
| 30B1-2 | | 30B1-293-2 | 36.93 | x | 38.73 |
| | | 30B1-294-2 | 39.80 | x | 40.62 |
| | | 30B1-295-2 | 31.96 | x | 32.80 |
| | | 30B0-296-2 | 31.62 | x | 32.57 |
| | | 30B0-297-2 | x | | |
| 30B0-2 | | 30B0-298-2 | 35.20 | x | 36.84 |
| | | 30B0-299-2 | 29.30 | x | x |
| | | 30B0-300-2 | 27.19 | x | 25.38 |
| | | 31B1-301-2 | x | | |
| | | 31B1-302-2 | x | | |
| 31B1-2 | | 31B1-303-2 | x | | |
| | | 31B1-304-2 | x | | |
| | | 31B1-305-2 | x | | |
| | | 31B0-306-2 | x | | |
| | | 31B0-307-2 | x | | |
| 31B0-2 | | 31B0-308-2 | x | | |
| | | 31B0-309-2 | x | | |
| | | 31B0-310-2 | 26.71 | x | 28.15 |
| | | 32B1-311-2 | x | | |
| | | 32B1-312-2 | 34.55 | x | 40.57 |
| 32B1-2 | 37.84 | 32B1-313-2 | 42.58 | | |
| | | 32B1-314-2 | 42.12 | | |
| | | 32B1-315-2 | x | | |
| | | 32B0-316-2 | | | |
| | | 32B0-317-2 | | | |
| 32B0-2 | x | 32B0-318-2 | | | |
| | | 32B0-319-2 | | | |
| | | 32B0-320-2 | | | |
| | | 33B1-321-2 | 38.07 | x | x |
| | | ~~33B1-322-2~~ | | | |
| 33B1-2 | | ~~33B1-323-2~~ | | | |
| | | 33B1-324-2 | 40.62 | | |
| | | 33B1-325-2 | 44.91 | | |
| | | 33B0-326-2 | 34.93 | x | 36.99 |
| | | 33B0-327-2 | 34.89 | x | 36.02 |
| 33B0-2 | | 33B0-328-2 | 37.25 | x | x |
| | | 33B0-329-2 | 35.97 | x | 38.10 |
| | | 33B0-330-2 | 36.57 | x | x |
| | | 34B1-331-2 | 33.26 | x | 34.39 |
| | | 34B1-332-2 | 34.61 | x | x |
| 34B1-2 | | 34B1-333-2 | 34.63 | x | 38.82 |
| | | 34B1-334-2 | 34.06 | x | 35.70 |
| | | 34B1-335-2 | 37.05 | x | 41.66 |
| | | 34B0-336-2 | 36.97 | x | 40.46 |
| | | 34B0-337-2 | 36.94 | x | x |
| 34B0-2 | | 34B0-338-2 | 39.21 | x | 40.35 |
| | | 34B0-339-2 | 40.69 | | |
| | | 34B0-340-2 | 36.07 | x | x |

| T3 | | | | | |
|---|---|---|---|---|---|
| Pool ID | Ct (M) | Sample ID | Ct (M) | Ct (H5 HA) | Ct (H9 HA) |
| | | 30B1-291-3 | 42.90 | | |
| | | 30B1-292-3 | | x | |
| 30B1-3 | | 30B1-293-3 | 30.67 | x | 30.21 |
| | | 30B1-294-3 | 26.55 | x | 28.43 |
| | | 30B1-295-3 | | x | |
| | | 30B0-296-3 | | x | |
| | | 30B0-297-3 | x | | |
| 30B0-3 | | 30B0-298-3 | 23.05 | x | 24.83 |
| | | 30B0-299-3 | | x | 29.02 |
| | | 30B0-300-3 | | x | |
| | | 31B1-301-3 | 39.49 | x | x |
| | | 31B1-302-3 | 36.18 | x | x |
| 31B1-3 | | 31B1-303-3 | 25.73 | x | 27.80 |
| | | 31B1-304-3 | 36.02 | x | x |
| | | 31B1-305-3 | 33.95 | x | 33.75 |
| | | 31B0-306-3 | 25.94 | x | 29.54 |
| | | 31B0-307-3 | 23.98 | x | 29.41 |
| 31B0-3 | | 31B0-308-3 | x | | |
| | | 31B0-309-3 | 26.22 | x | 29.00 |
| | | 31B0-310-3 | | x | |
| | | 32B1-311-3 | 35.76 | x | 39.40 |
| | | 32B1-312-3 | 21.28 | x | 23.02 |
| 32B1-3 | | 32B1-313-3 | x | | |
| | | 32B1-314-3 | 35.68 | x | 38.07 |
| | | 32B1-315-3 | 23.24 | x | 25.95 |
| | | 32B0-316-3 | 27.18 | x | 29.58 |
| | | 32B0-317-3 | 34.65 | x | 38.04 |
| 32B0-3 | | 32B0-318-3 | x | | |
| | | 32B0-319-3 | 34.96 | x | 36.89 |
| | | 32B0-320-3 | x | | |
| | | 33B1-321-3 | x | | |
| | | ~~33B1-322-3~~ | | | |
| 33B1-3 | | ~~33B1-323-3~~ | | | |
| | | 33B1-324-3 | x | | |
| | | 33B1-325-3 | 24.56 | x | 27.18 |
| | | 33B0-326-3 | x | | x |
| | | 33B0-327-3 | 25.90 | x | 28.57 |
| 33B0-3 | | 33B0-328-3 | 29.51 | x | 31.41 |
| | | 33B0-329-3 | 30.48 | x | 33.41 |
| | | 33B0-330-3 | x | x | x |
| | | 34B1-331-3 | 25.63 | x | 27.69 |
| | | 34B1-332-3 | 26.10 | x | 27.03 |
| 34B1-3 | | 34B1-333-3 | 22.96 | x | 24.23 |
| | | 34B1-334-3 | 23.73 | x | 24.43 |
| | | 34B1-335-3 | 25.63 | x | 26.47 |
| | | 34B0-336-3 | 25.33 | x | 26.48 |
| | | 34B0-337-3 | 26.03 | x | 26.99 |
| 34B0-3 | | 34B0-338-3 | 27.02 | x | 28.32 |
| | | 34B0-339-3 | 31.24 | x | 32.58 |
| | | 34B0-340-3 | 32.71 | 40.79 | 33.93 |

| T4 | | | | | |
|---|---|---|---|---|---|
| Pool ID | Ct (M) | Sample ID | Ct (M) | Ct (H5 HA) | Ct (H9 HA) |
| | | 30B1-291-4 | 34.92 | x | x |
| | | 30B1-292-4 | | x | |
| 30B1-4 | | 30B1-293-4 | | x | |
| | | 30B1-294-4 | | x | |
| | | 30B1-295-4 | | x | |
| | | 30B0-296-4 | | x | |
| | | 30B0-297-4 | 31.80 | x | 33.85 |
| 30B0-4 | | 30B0-298-4 | | x | |
| | | 30B0-299-4 | | x | |
| | | 30B0-300-4 | | x | |
| | | 31B1-301-4 | 22.42 | x | 23.06 |
| | | 31B1-302-4 | 33.14 | x | 33.70 |
| 31B1-4 | | 31B1-303-4 | | x | |
| | | 31B1-304-4 | 24.79 | x | 25.83 |
| | | 31B1-305-4 | 39.64 | x | x |
| | | 31B0-306-4 | | x | |
| | | 31B0-307-4 | | x | |
| 31B0-4 | | 31B0-308-4 | x | | |
| | | 31B0-309-4 | | x | |
| | | 31B0-310-4 | | x | |
| | | 32B1-311-4 | 34.29 | x | 36.03 |
| | | 32B1-312-4 | | x | |
| 32B1-4 | | 32B1-313-4 | 40.25 | | |
| | | 32B1-314-4 | 35.16 | x | 35.89 |
| | | 32B1-315-4 | | x | |
| | | 32B0-316-4 | | x | |
| | | 32B0-317-4 | 25.73 | x | 26.58 |
| 32B0-4 | | 32B0-318-4 | 18.52 | x | 17.02 |
| | | 32B0-319-4 | 22.64 | x | 28.75 |
| | | 32B0-320-4 | 25.64 | x | 37.14 |
| | | 33B1-321-4 | 34.89 | x | 43.04 |
| | | ~~33B1-322-4~~ | | | |
| 33B1-4 | | ~~33B1-323-4~~ | | | |
| | | 33B1-324-4 | 19.32 | x | 27.66 |
| | | 33B1-325-4 | | x | |
| | | 33B0-326-4 | 39.87 | x | 43.94 |
| | | 33B0-327-4 | | x | |
| 33B0-4 | | 33B0-328-4 | | x | |
| | | 33B0-329-4 | | x | 26.42 |
| | | 33B0-330-4 | 29.26 | x | 31.75 |
| | | 34B1-331-4 | | x | |
| | | 34B1-332-4 | | x | |
| 34B1-4 | | 34B1-333-4 | | x | |
| | | 34B1-334-4 | | x | |
| | | 34B1-335-4 | | x | |
| | | 34B0-336-4 | | x | |
| | | 34B0-337-4 | | x | |
| 34B0-4 | | 34B0-338-4 | | x | |
| | | 34B0-339-4 | | x | |
| | | 34B0-340-4 | | x | 29.01 |

**T0**

| Pool ID | Ct (M) | Sample ID | Ct (M) | Ct (H5 HA) | Ct (H9 HA) |
|---|---|---|---|---|---|
| | | 35A1-341-0 | 21.31 | | 20.34 |
| | | 35A1-342-0 | 22.78 | | 23.71 |
| 35A1-0 | 22.27 | 35A1-343-0 | 20.33 | x | 22.25 |
| | | 35A1-344-0 | 21.71 | | 22.29 |
| | | 35A1-345-0 | 20.21 | | 21.22 |
| | | - | | | |
| | | 36A1-351-0 | | | |
| | | 36A1-352-0 | | | |
| 36A1-0 | x | 36A1-353-0 | | | |
| | | 36A1-354-0 | | | |
| | | 36A1-355-0 | | | |
| | | - | | | |
| | | 37A1-361-0 | | | |
| | | 37A1-362-0 | | | |
| 37A1-0 | x | 37A1-363-0 | | | |
| | | 37A1-364-0 | | | |
| | | 37A1-365-0 | | | |
| | | - | | | |
| | | 38A1-371-0 | | | |
| | | 38A1-372-0 | | | |
| 38A1-0 | x | 38A1-373-0 | | | |
| | | 38A1-374-0 | | | |
| | | 38A1-375-0 | | | |
| | | - | | | |
| | | 39A1-381-0 | | | |
| | | 39A1-382-0 | | | |
| 39A1-0 | x | 39A1-383-0 | | | |
| | | 39A1-384-0 | | | |
| | | 39A1-385-0 | | | |
| | | - | | | |

**T1**

| Pool ID | Ct (M) | Sample ID | Ct (M) | Ct (H5 HA) | Ct (H9 HA) |
|---|---|---|---|---|---|
| | | 35A1-341-1 | 23.57 | | 24.79 |
| | | 35A1-342-1 | 23.57 | | 26.26 |
| 35A1-1 | 26.17 | 35A1-343-1 | 23.69 | x | 24.34 |
| | | 35A1-344-1 | 25.19 | | 25.82 |
| | | 35A1-345-1 | 22.44 | | 23.94 |
| | | 35A0-346-1 | x | | |
| | | 35A0-347-1 | x | | |
| 35A0-1 | 22.77 | 35A0-348-1 | 20.40 | x | 21.67 |
| | | 35A0-349-1 | x | | |
| | | 35A0-350-1 | x | | |
| | | 36A1-351-1 | | | |
| | | 36A1-352-1 | | | |
| 36A1-1 | x | 36A1-353-1 | | | |
| | | 36A1-354-1 | | | |
| | | 36A1-355-1 | | | |
| | | 36A0-356-1 | | | |
| | | 36A0-357-1 | | | |
| 36A0-1 | x | 36A0-358-1 | | | |
| | | 36A0-359-1 | | | |
| | | 36A0-360-1 | | | |
| | | 37A1-361-1 | | | |
| | | 37A1-362-1 | | | |
| 37A1-1 | x | 37A1-363-1 | | | |
| | | 37A1-364-1 | | | |
| | | 37A1-365-1 | | | |
| | | 37A0-366-1 | | | |
| | | 37A0-367-1 | | | |
| 37A0-1 | x | 37A0-368-1 | | | |
| | | 37A0-369-1 | | | |
| | | 37A0-370-1 | | | |
| | | 38A1-371-1 | | | |
| | | 38A1-372-1 | | | |
| 38A1-1 | x | 38A1-373-1 | | | |
| | | 38A1-374-1 | | | |
| | | 38A1-375-1 | | | |
| | | 38A0-376-1 | | | |
| | | 38A0-377-1 | | | |
| 38A0-1 | x | 38A0-378-1 | | | |
| | | 38A0-379-1 | | | |
| | | 38A0-380-1 | | | |
| | | 39A1-381-1 | | | |
| | | 39A1-382-1 | | | |
| 39A1-1 | x | 39A1-383-1 | | | |
| | | 39A1-384-1 | | | |
| | | 39A1-385-1 | | | |
| | | 39A0-386-1 | | | |
| | | 39A0-387-1 | | | |
| 39A0-1 | x | 39A0-388-1 | | | |
| | | 39A0-389-1 | | | |
| | | 39A0-390-1 | | | |

**T2**

| Pool ID | Ct (M) | Sample ID | Ct (M) | Ct (H5 HA) | Ct (H9 HA) |
|---|---|---|---|---|---|
| | | 35A1-341-2 | | | |
| | | 35A1-342-2 | | | |
| 35A1-2 | | 35A1-343-2 | | x | |
| | | 35A1-344-2 | | | |
| | | 35A1-345-2 | | | |
| | | 35A0-346-2 | x | | |
| | | 35A0-347-2 | 30.02 | x | 32.27 |
| 35A0-2 | | 35A0-348-2 | | x | |
| | | 35A0-349-2 | x | | |
| | | 35A0-350-2 | 32.70 | x | 32.64 |
| | | 36A1-351-2 | | | |
| | | 36A1-352-2 | | | |
| 36A1-2 | 32.87 | 36A1-353-2 | 28.41 | x | 30.24 |
| | | 36A1-354-2 | x | | |
| | | 36A1-355-2 | x | | |
| | | 36A0-356-2 | x | | |
| | | 36A0-357-2 | x | | |
| 36A0-2 | 31.63 | 36A0-358-2 | x | | |
| | | 36A0-359-2 | x | | |
| | | 36A0-360-2 | 25.71 | x | 28.30 |
| | | 37A1-361-2 | 28.06 | | 30.27 |
| | | 37A1-362-2 | 21.47 | | 23.90 |
| 37A1-2 | 25.97 | 37A1-363-2 | 24.35 | x | 26.81 |
| | | 37A1-364-2 | x | | |
| | | 37A1-365-2 | 27.90 | | 29.64 |
| | | 37A0-366-2 | 30.64 | | 32.75 |
| | | 37A0-367-2 | 23.82 | | 25.84 |
| 37A0-2 | 27.43 | 37A0-368-2 | 27.53 | x | 29.80 |
| | | 37A0-369-2 | 23.83 | | 26.36 |
| | | 37A0-370-2 | 26.88 | | 29.64 |
| | | 38A1-371-2 | | | |
| | | 38A1-372-2 | | | |
| 38A1-2 | x | 38A1-373-2 | | | |
| | | 38A1-374-2 | | | |
| | | 38A1-375-2 | | | |
| | | 38A0-376-2 | | | |
| | | 38A0-377-2 | | | |
| 38A0-2 | x | 38A0-378-2 | | | |
| | | 38A0-379-2 | | | |
| | | 38A0-380-2 | | | |
| | | 39A1-381-2 | x | | |
| | | 39A1-382-2 | 28.76 | x | 28.78 |
| 39A1-2 | 33.51 | 39A1-383-2 | x | | |
| | | 39A1-384-2 | 34.97 | x | 38.30 |
| | | 39A1-385-2 | x | | |
| | | 39A0-386-2 | 24.20 | x | 26.39 |
| | | 39A0-387-2 | x | | |
| 39A0-2 | 29.79 | 39A0-388-2 | x | | |
| | | 39A0-389-2 | x | | |
| | | 39A0-390-2 | x | | |

**T3**

| Pool ID | Ct (M) | Sample ID | Ct (M) | Ct (H5 HA) | Ct (H9 HA) |
|---|---|---|---|---|---|
| | | 35A1-341-3 | | | |
| | | 35A1-342-3 | 33.48 | | |
| 35A1-3 | | 35A1-343-3 | 32.96 | x | |
| | | 35A1-344-3 | | | |
| | | 35A1-345-3 | | | |
| | | 35A0-346-3 | 21.48 | | 22.40 |
| | | 35A0-347-3 | | | |
| 35A0-3 | | 35A0-348-3 | | x | |
| | | 35A0-349-3 | 19.99 | | 18.71 |
| | | 35A0-350-3 | | | |
| | | 36A1-351-3 | | | |
| | | 36A1-352-3 | | | |
| 36A1-3 | | 36A1-353-3 | | x | |
| | | 36A1-354-3 | 24.82 | x | 27.69 |
| | | 36A1-355-3 | 21.01 | x | 24.35 |
| | | 36A0-356-3 | 31.92 | | 35.39 |
| | | 36A0-357-3 | 21.54 | | 24.63 |
| 36A0-3 | | 36A0-358-3 | 23.61 | x | 26.59 |
| | | 36A0-359-3 | 22.98 | | 25.73 |
| | | 36A0-360-3 | | | |
| | | 37A1-361-3 | | | |
| | | 37A1-362-3 | 20.94 | | |
| 37A1-3 | | 37A1-363-3 | | x | |
| | | 37A1-364-3 | | | |
| | | 37A1-365-3 | | | |
| | | 37A0-366-3 | | | |
| | | 37A0-367-3 | | | |
| 37A0-3 | | 37A0-368-3 | | x | |
| | | 37A0-369-3 | | | |
| | | 37A0-370-3 | | | |
| | | 38A1-371-3 | 21.93 | | 25.13 |
| | | 38A1-372-3 | 20.86 | | 20.77 |
| 38A1-3 | 22.05 | 38A1-373-3 | 21.42 | x | 21.34 |
| | | 38A1-374-3 | 20.71 | | 20.50 |
| | | 38A1-375-3 | 23.67 | | 22.87 |
| | | 38A0-376-3 | 21.74 | | 22.47 |
| | | 38A0-377-3 | 18.60 | | 18.21 |
| 38A0-3 | 21.60 | 38A0-378-3 | 24.48 | x | 24.25 |
| | | 38A0-379-3 | 21.21 | | 29.88 |
| | | 38A0-380-3 | 22.47 | | 22.75 |
| | | 39A1-381-3 | 26.17 | | 28.77 |
| | | 39A1-382-3 | | | |
| 39A1-3 | | 39A1-383-3 | 21.70 | x | 21.84 |
| | | 39A1-384-3 | 23.56 | | 26.40 |
| | | 39A1-385-3 | 22.88 | | 22.94 |
| | | 39A0-386-3 | | | |
| | | 39A0-387-3 | 23.87 | | 23.42 |
| 39A0-3 | | 39A0-388-3 | 24.58 | x | 24.03 |
| | | 39A0-389-3 | 20.46 | | 21.51 |
| | | 39A0-390-3 | 31.75 | | 31.55 |

**T4**

| Pool ID | Ct (M) | Sample ID | Ct (M) | Ct (H5 HA) | Ct (H9 HA) |
|---|---|---|---|---|---|
| | | 35A1-341-4 | | | |
| | | 35A1-342-4 | 30.94 | | |
| 35A1-4 | | 35A1-343-4 | | x | |
| | | 35A1-344-4 | | | |
| | | 35A1-345-4 | | | |
| | | 35A0-346-4 | | | |
| | | 35A0-347-4 | | | |
| 35A0-4 | | 35A0-348-4 | | x | |
| | | 35A0-349-4 | | | |
| | | 35A0-350-4 | | | |
| | | 36A1-351-4 | | | |
| | | 36A1-352-4 | | | |
| 36A1-4 | | 36A1-353-4 | | x | |
| | | 36A1-354-4 | | x | |
| | | 36A1-355-4 | | x | |
| | | 36A0-356-4 | | | 21.38 |
| | | 36A0-357-4 | | | |
| 36A0-4 | | 36A0-358-4 | | x | |
| | | 36A0-359-4 | | | |
| | | 36A0-360-4 | | | |
| | | 37A1-361-4 | | x | |
| | | 37A1-362-4 | | | |
| 37A1-4 | | 37A1-363-4 | | x | |
| | | 37A1-364-4 | | | |
| | | 37A1-365-4 | | x | |
| | | 37A0-366-4 | | | |
| | | 37A0-367-4 | | | |
| 37A0-4 | | 37A0-368-4 | | x | |
| | | 37A0-369-4 | | | |
| | | 37A0-370-4 | | | |
| | | 38A1-371-4 | 21.05 | | |
| | | 38A1-372-4 | | | |
| 38A1-4 | | 38A1-373-4 | | x | |
| | | 38A1-374-4 | | | |
| | | 38A1-375-4 | | | |
| | | 38A0-376-4 | | | |
| | | 38A0-377-4 | | | |
| 38A0-4 | | 38A0-378-4 | | x | |
| | | 38A0-379-4 | | | |
| | | 38A0-380-4 | | | |
| | | 39A1-381-4 | | | |
| | | 39A1-382-4 | | | |
| 39A1-4 | | 39A1-383-4 | | x | |
| | | 39A1-384-4 | | | |
| | | 39A1-385-4 | | | |
| | | 39A0-386-4 | | | |
| | | 39A0-387-4 | | | |
| 39A0-4 | | 39A0-388-4 | | x | |
| | | 39A0-389-4 | 21.08 | | |
| | | 39A0-390-4 | | | |

| T0 | | | | | | T1 | | | | | | T2 | | | | | | T3 | | | | | | T4 | | | | | |
|---|---|---|---|---|---|---|---|---|---|---|---|---|---|---|---|---|---|---|---|---|---|---|---|---|---|---|---|---|---|
| Pool ID | Ct (M) | Sample ID | Ct (M) | Ct (H5 HA) | Ct (H9 HA) | Pool ID | Ct (M) | Sample ID | Ct (M) | Ct (H5 HA) | Ct (H9 HA) | Pool ID | Ct (M) | Sample ID | Ct (M) | Ct (H5 HA) | Ct (H9 HA) | Pool ID | Ct (M) | Sample ID | Ct (M) | Ct (H5 HA) | Ct (H9 HA) | Pool ID | Ct (M) | Sample ID | Ct (M) | Ct (H5 HA) | Ct (H9 HA) |
| | | 40A1-391-0 | | | | | | 40A1-391-1 | | | | | | 40A1-391-2 | 25.74 | | 28.34 | | | 40A1-391-3 | | | | | | 40A1-391-4 | | | |
| | | 40A1-392-0 | | | | | | 40A1-392-1 | | | | | | 40A1-392-2 | 31.22 | | 34.36 | | | 40A1-392-3 | | | 25.32 | | | 40A1-392-4 | | | |
| 40A1-0 | x | 40A1-393-0 | | | | 40A1-1 | x | 40A1-393-1 | | | | 40A1-2 | 28.46 | 40A1-393-2 | 26.87 | x | 28.72 | 40A1-3 | | 40A1-393-3 | | x | | 40A1-4 | | 40A1-393-4 | | x | |
| | | 40A1-394-0 | | | | | | 40A1-394-1 | | | | | | 40A1-394-2 | 27.24 | | 30.57 | | | 40A1-394-3 | | | | | | 40A1-394-4 | | | |
| | | 40A1-395-0 | | | | | | 40A1-395-1 | | | | | | 40A1-395-2 | 25.16 | | 27.32 | | | 40A1-395-3 | | | | | | 40A1-395-4 | | | |
| | | | | | | | | 40A0-396-1 | | | | | | 40A0-396-2 | 30.70 | | 33.80 | | | 40A0-396-3 | | | 22.19 | | | 40A0-396-4 | | | |
| | | | | | | | | 40A0-397-1 | | | | | | 40A0-397-2 | x | | | | | 40A0-397-3 | 20.23 | | 23.51 | | | 40A0-397-4 | | | |
| | | - | | | | 40A0-1 | x | 40A0-398-1 | | | | 40A0-2 | 34.52 | 40A0-398-2 | 35.95 | x | x | 40A0-3 | | 40A0-398-3 | 24.29 | x | 27.32 | 40A0-4 | | 40A0-398-4 | | x | |
| | | | | | | | | 40A0-399-1 | | | | | | 40A0-399-2 | 36.11 | | x | | | 40A0-399-3 | 25.76 | | 28.64 | | | 40A0-399-4 | | | |
| | | | | | | | | 40A0-400-1 | | | | | | 40A0-400-2 | 34.37 | | 38.77 | | | 40A0-400-3 | 23.29 | | 25.58 | | | 40A0-400-4 | | | |
| | | 41A1-401-0 | | | | | | 41A1-401-1 | | | | | | 41A1-401-2 | 22.91 | | 25.97 | | | 41A1-401-3 | | | | | | 41A1-401-4 | | | |
| | | 41A1-402-0 | | | | | | 41A1-402-1 | | | | | | 41A1-402-2 | 23.94 | | 26.95 | | | 41A1-402-3 | | | | | | 41A1-402-4 | | | |
| 41A1-0 | x | 41A1-403-0 | | | | 41A1-1 | x | 41A1-403-1 | | | | 41A1-2 | 23.88 | 41A1-403-2 | 21.25 | x | 23.49 | 41A1-3 | | 41A1-403-3 | | x | | 41A1-4 | | 41A1-403-4 | | x | |
| | | 41A1-404-0 | | | | | | 41A1-404-1 | | | | | | 41A1-404-2 | 25.70 | | 27.20 | | | 41A1-404-3 | | | | | | 41A1-404-4 | | | |
| | | 41A1-405-0 | | | | | | 41A1-405-1 | | | | | | 41A1-405-2 | 27.09 | | 29.31 | | | 41A1-405-3 | | | | | | 41A1-405-4 | | | |
| | | | | | | | | 41A0-406-1 | | | | | | 41A0-406-2 | 23.79 | | 26.61 | | | 41A0-406-3 | | | | | | 41A0-406-4 | | | |
| | | | | | | | | 41A0-407-1 | | | | | | 41A0-407-2 | 22.73 | | 25.50 | | | 41A0-407-3 | | | | | | 41A0-407-4 | | | |
| | | - | | | | 41A0-1 | x | 41A0-408-1 | | | | 41A0-2 | 26.07 | 41A0-408-2 | x | x | | 41A0-3 | | 41A0-408-3 | 22.36 | x | 25.58 | 41A0-4 | | 41A0-408-4 | | x | |
| | | | | | | | | 41A0-409-1 | | | | | | 41A0-409-2 | 29.54 | | 31.66 | | | 41A0-409-3 | | | | | | 41A0-409-4 | | | |
| | | | | | | | | 41A0-410-1 | | | | | | 41A0-410-2 | 25.48 | | 27.35 | | | 41A0-410-3 | | | | | | 41A0-410-4 | 21.87 | | |
| | | 42A1-411-0 | | | | | | 42A1-411-1 | 31.83 | | 30.45 | | | 42A1-411-2 | | | | | | 42A1-411-3 | | | | | | 42A1-411-4 | | | |
| | | 42A1-412-0 | | | | | | 42A1-412-1 | 31.57 | | 32.13 | | | 42A1-412-2 | 33.01 | | 24.11 | | | 42A1-412-3 | 22.40 | | | | | 42A1-412-4 | | | |
| 42A1-0 | x | 42A1-413-0 | | | | 42A1-1 | 32.09 | 42A1-413-1 | 31.52 | x | 32.93 | 42A1-2 | | ~~42A1-413-2~~ | | x | | 42A1-3 | | ~~42A1-413-3~~ | | x | | 42A1-4 | | ~~42A1-413-4~~ | | x | |
| | | 42A1-414-0 | | | | | | 42A1-414-1 | 31.84 | | 30.36 | | | 42A1-414-2 | | | | | | 42A1-414-3 | | | | | | 42A1-414-4 | | | |
| | | 42A1-415-0 | | | | | | 42A1-415-1 | 31.77 | | 30.73 | | | 42A1-415-2 | 32.44 | | 39.55 | | | 42A1-415-3 | | | 20.49 | | | 42A1-415-4 | | | |
| | | | | | | | | 42A0-416-1 | 31.68 | | 30.39 | | | 42A0-416-2 | | | | | | 42A0-416-3 | | | | | | 42A0-416-4 | | | |
| | | | | | | | | 42A0-417-1 | 31.28 | | 30.43 | | | 42A0-417-2 | | | | | | 42A0-417-3 | | | | | | 42A0-417-4 | | | |
| | | - | | | | 42A0-1 | 31.72 | 42A0-418-1 | 31.75 | x | 31.00 | 42A0-2 | | 42A0-418-2 | | x | | 42A0-3 | | 42A0-418-3 | | x | | 42A0-4 | | 42A0-418-4 | | x | |
| | | | | | | | | 42A0-419-1 | 31.45 | | 30.68 | | | 42A0-419-2 | 32.85 | | 35.36 | | | 42A0-419-3 | | | 20.04 | | | 42A0-419-4 | | | |
| | | | | | | | | 42A0-420-1 | 31.35 | | 30.13 | | | 42A0-420-2 | 31.36 | | 32.16 | | | 42A0-420-3 | | | | | | 42A0-420-4 | | | |
| | | 43A1-421-0 | | | | | | 43A1-421-1 | 31.56 | | 30.25 | | | 43A1-421-2 | | | | | | 43A1-421-3 | | | | | | 43A1-421-4 | | | |
| | | 43A1-422-0 | | | | | | 43A1-422-1 | 31.68 | | 30.67 | | | 43A1-422-2 | 29.86 | | 32.24 | | | 43A1-422-3 | | | | | | 43A1-422-4 | | | |
| 43A1-0 | x | 43A1-423-0 | | | | 43A1-1 | 32.11 | 43A1-423-1 | 31.54 | x | 30.23 | 43A1-2 | | 43A1-423-2 | | x | | 43A1-3 | | 43A1-423-3 | | x | | 43A1-4 | | 43A1-423-4 | | x | |
| | | 43A1-424-0 | | | | | | 43A1-424-1 | 31.08 | | 29.79 | | | 43A1-424-2 | | | | | | 43A1-424-3 | | | | | | 43A1-424-4 | | | |
| | | 43A1-425-0 | | | | | | 43A1-425-1 | 31.03 | | 30.74 | | | 43A1-425-2 | 31.07 | | 34.20 | | | 43A1-425-3 | | | 19.44 | | | 43A1-425-4 | | | |
| | | | | | | | | 43A0-426-1 | 31.18 | | 30.34 | | | 43A0-426-2 | | | | | | 43A0-426-3 | | | | | | 43A0-426-4 | | | |
| | | | | | | | | 43A0-427-1 | 31.17 | | 31.74 | | | 43A0-427-2 | 33.94 | | 36.14 | | | 43A0-427-3 | 20.58 | | 20.03 | | | 43A0-427-4 | | | |
| | | - | | | | 43A0-1 | 32.21 | 43A0-428-1 | 31.20 | x | 30.32 | 43A0-2 | | 43A0-428-2 | | x | | 43A0-3 | | 43A0-428-3 | | x | | 43A0-4 | | 43A0-428-4 | | x | |
| | | | | | | | | 43A0-429-1 | 31.43 | | 30.10 | | | 43A0-429-2 | | | | | | 43A0-429-3 | | | | | | 43A0-429-4 | | | |
| | | | | | | | | 43A0-430-1 | 31.27 | | 29.47 | | | 43A0-430-2 | 23.37 | | 21.85 | | | 43A0-430-3 | | | | | | 43A0-430-4 | | | |
| | | 44A1-431-0 | | | | | | 44A1-431-1 | 31.28 | | 29.81 | | | 44A1-431-2 | | | | | | 44A1-431-3 | | | | | | 44A1-431-4 | | | |
| | | 44A1-432-0 | | | | | | 44A1-432-1 | 31.54 | | 29.51 | | | 44A1-432-2 | 31.45 | | 33.58 | | | 44A1-432-3 | | | 26.60 | | | 44A1-432-4 | | | |
| 44A1-0 | x | 44A1-433-0 | | | | 44A1-1 | 31.59 | 44A1-433-1 | 31.75 | x | 31.47 | 44A1-2 | | 44A1-433-2 | 32.64 | x | 34.59 | 44A1-3 | | 44A1-433-3 | | x | 23.30 | 44A1-4 | | 44A1-433-4 | | x | |
| | | 44A1-434-0 | | | | | | 44A1-434-1 | 31.24 | | 30.31 | | | 44A1-434-2 | | | | | | 44A1-434-3 | | | | | | 44A1-434-4 | | | |
| | | 44A1-435-0 | | | | | | 44A1-435-1 | 31.24 | | 30.66 | | | 44A1-435-2 | | | | | | 44A1-435-3 | | | | | | 44A1-435-4 | | | |
| | | | | | | | | 44A0-436-1 | 31.47 | | 30.85 | | | 44A0-436-2 | | | | | | 44A0-436-3 | | | | | | 44A0-436-4 | | | |
| | | | | | | | | 44A0-437-1 | 31.25 | | 30.04 | | | 44A0-437-2 | 31.87 | | 33.98 | | | 44A0-437-3 | | | 20.06 | | | 44A0-437-4 | | | |
| | | - | | | | 44A0-1 | 31.81 | 44A0-438-1 | 31.65 | x | 29.77 | 44A0-2 | | 44A0-438-2 | 32.57 | x | 34.32 | 44A0-3 | | 44A0-438-3 | | x | 24.36 | 44A0-4 | | 44A0-438-4 | | x | |
| | | | | | | | | 44A0-439-1 | 31.72 | | 30.30 | | | 44A0-439-2 | | | | | | 44A0-439-3 | | | | | | 44A0-439-4 | | | |
| | | | | | | | | 44A0-440-1 | 31.38 | | 30.03 | | | 44A0-440-2 | | | | | | 44A0-440-3 | | | | | | 44A0-440-4 | | | |

| T0 | | | | | | T1 | | | | | | T2 | | | | | | T3 | | | | | | T4 | | | | | |
|---|---|---|---|---|---|---|---|---|---|---|---|---|---|---|---|---|---|---|---|---|---|---|---|---|---|---|---|---|---|
| Pool ID | Ct (M) | Sample ID | Ct (M) | Ct (H5 HA) | Ct (H9 HA) | Pool ID | Ct (M) | Sample ID | Ct (M) | Ct (H5 HA) | Ct (H9 HA) | Pool ID | Ct (M) | Sample ID | Ct (M) | Ct (H5 HA) | Ct (H9 HA) | Pool ID | Ct (M) | Sample ID | Ct (M) | Ct (H5 HA) | Ct (H9 HA) | Pool ID | Ct (M) | Sample ID | Ct (M) | Ct (H5 HA) | Ct (H9 HA) |
| | | 45A1-441-0 | 31.66 | | 31.50 | | | 45A1-441-1 | 31.50 | | | | | 45A1-441-2 | | | | | | 45A1-441-3 | | | | | | 45A1-441-4 | | | |
| | | 45A1-442-0 | 31.88 | | 30.68 | | | 45A1-442-1 | 30.12 | | | | | 45A1-442-2 | 31.94 | | 34.06 | | | 45A1-442-3 | | | 22.03 | | | 45A1-442-4 | | | |
| 45A1-0 | 31.97 | 45A1-443-0 | 31.88 | x | 31.09 | 45A1-1 | 31.28 | 45A1-443-1 | 31.75 | x | | 45A1-2 | | 45A1-443-2 | | x | | 45A1-3 | | 45A1-443-3 | | x | | 45A1-4 | | 45A1-443-4 | | x | |
| | | 45A1-444-0 | 31.77 | | 30.65 | | | 45A1-444-1 | 31.12 | | | | | 45A1-444-2 | | | | | | 45A1-444-3 | | | | | | 45A1-444-4 | | | |
| | | 45A1-445-0 | 31.66 | | 35.45 | | | 45A1-445-1 | 31.17 | | 33.51 | | | 45A1-445-2 | 30.44 | | 32.29 | | | 45A1-445-3 | | | | | | 45A1-445-4 | | | |
| | | | | | | | | 45A0-446-1 | 31.99 | | 30.83 | | | 45A0-446-2 | | | | | | 45A0-446-3 | | | | | | 45A0-446-4 | | | |
| | | | | | | | | 45A0-447-1 | 31.70 | | 30.36 | | | 45A0-447-2 | | | | | | 45A0-447-3 | | | | | | 45A0-447-4 | | | |
| | | - | | | | 45A0-1 | 31.94 | 45A0-448-1 | 31.80 | x | 30.15 | 45A0-2 | | 45A0-448-2 | 27.40 | x | 29.24 | 45A0-3 | | 45A0-448-3 | | x | | 45A0-4 | | 45A0-448-4 | | x | |
| | | | | | | | | 45A0-449-1 | 31.65 | | 30.11 | | | 45A0-449-2 | | | | | | 45A0-449-3 | | | | | | 45A0-449-4 | | | |
| | | | | | | | | 45A0-450-1 | 31.13 | | 30.48 | | | 45A0-450-2 | 32.40 | | 36.24 | | | 45A0-450-3 | | | 24.20 | | | 45A0-450-4 | | | |
| | | 46A1-451-0 | 31.25 | | 30.00 | | | 46A1-451-1 | 31.73 | | | | | 46A1-451-2 | | | | | | 46A1-451-3 | | | | | | 46A1-451-4 | | | |
| | | 46A1-452-0 | 31.56 | | 30.97 | | | 46A1-452-1 | 31.18 | | | | | 46A1-452-2 | | | | | | 46A1-452-3 | | | | | | 46A1-452-4 | | | |
| 46A1-0 | 31.36 | 46A1-453-0 | 30.97 | x | 30.71 | 46A1-1 | 31.47 | 46A1-453-1 | 31.40 | x | | 46A1-2 | | 46A1-453-2 | x | x | | 46A1-3 | | 46A1-453-3 | 25.90 | x | | 46A1-4 | | 46A1-453-4 | | x | |
| | | 46A1-454-0 | 31.41 | | 30.12 | | | 46A1-454-1 | 31.46 | | | | | 46A1-454-2 | 37.64 | | x | | | 46A1-454-3 | 23.77 | | 23.30 | | | 46A1-454-4 | | | |
| | | 46A1-455-0 | 31.39 | | 30.10 | | | 46A1-455-1 | 30.93 | | | | | 46A1-455-2 | | | | | | 46A1-455-3 | | | | | | 46A1-455-4 | | | |
| | | | | | | | | 46A0-456-1 | 31.55 | | 30.14 | | | 46A0-456-2 | 25.91 | | 29.91 | | | 46A0-456-3 | | | | | | 46A0-456-4 | | | |
| | | | | | | | | 46A0-457-1 | 31.70 | | 30.06 | | | 46A0-457-2 | | | | | | 46A0-457-3 | | | | | | 46A0-457-4 | | | |
| | | - | | | | 46A0-1 | 31.72 | 46A0-458-1 | 31.70 | x | 30.44 | 46A0-2 | | 46A0-458-2 | | x | | 46A0-3 | | 46A0-458-3 | | x | | 46A0-4 | | 46A0-458-4 | | x | |
| | | | | | | | | 46A0-459-1 | 31.54 | | 30.42 | | | 46A0-459-2 | | | | | | 46A0-459-3 | | | | | | 46A0-459-4 | 24.09 | | 26.30 |
| | | | | | | | | 46A0-460-1 | 27.45 | | 25.91 | | | 46A0-460-2 | 22.79 | | 23.62 | | | 46A0-460-3 | | | | | | 46A0-460-4 | | | |
| | | 47A1-461-0 | 31.16 | | 32.68 | | | 47A1-461-1 | x | | x | | | 47A1-461-2 | 29.29 | | 29.60 | | | 47A1-461-3 | | | | | | 47A1-461-4 | | | |
| | | 47A1-462-0 | 31.28 | | 31.94 | | | 47A1-462-1 | x | | x | | | 47A1-462-2 | 32.40 | | 33.95 | | | 47A1-462-3 | | | 21.94 | | | 47A1-462-4 | | | |
| 47A1-0 | 31.77 | 47A1-463-0 | 31.28 | x | 32.98 | 47A1-1 | x | 47A1-463-1 | x | | x | 47A1-2 | 33.55 | 47A1-463-2 | 30.10 | x | 31.87 | 47A1-3 | | 47A1-463-3 | | x | | 47A1-4 | | 47A1-463-4 | | x | |
| | | 47A1-464-0 | 31.35 | | 33.37 | | | 47A1-464-1 | x | | x | | | 47A1-464-2 | 30.25 | | 30.37 | | | 47A1-464-3 | | | | | | 47A1-464-4 | | | |
| | | 47A1-465-0 | 31.56 | | 32.48 | | | 47A1-465-1 | x | | x | | | 47A1-465-2 | 32.36 | | 33.78 | | | 47A1-465-3 | | | 22.64 | | | 47A1-465-4 | | | |
| | | | | | | | | 47A0-466-1 | x | | x | | | 47A0-466-2 | 32.24 | | 33.82 | | | 47A0-466-3 | | | 22.09 | | | 47A0-466-4 | | | |
| | | | | | | | | 47A0-467-1 | x | | x | | | 47A0-467-2 | 23.43 | | 26.40 | | | 47A0-467-3 | | | | | | 47A0-467-4 | | | |
| | | - | | | | 47A0-1 | x | 47A0-468-1 | x | | x | 47A0-2 | 28.84 | 47A0-468-2 | 31.98 | x | 33.77 | 47A0-3 | | 47A0-468-3 | | x | x | 47A0-4 | | 47A0-468-4 | | x | 21.07 |
| | | | | | | | | 47A0-469-1 | x | | x | | | 47A0-469-2 | 31.92 | | 33.64 | | | 47A0-469-3 | | | 21.37 | | | 47A0-469-4 | | | |
| | | | | | | | | 47A0-470-1 | x | | x | | | 47A0-470-2 | 36.09 | | x | | | 47A0-470-3 | 24.59 | | 27.54 | | | 47A0-470-4 | | | |
| | | 48A1-471-0 | 31.57 | | 32.24 | | | 48A1-471-1 | 36.90 | | 34.25 | | | 48A1-471-2 | 33.46 | | 36.50 | | | 48A1-471-3 | 20.71 | | 24.32 | | | 48A1-471-4 | | | |
| | | 48A1-472-0 | 31.49 | | 33.13 | | | 48A1-472-1 | x | | x | | | 48A1-472-2 | 27.92 | | 30.34 | | | 48A1-472-3 | | | | | | 48A1-472-4 | | | |
| 48A1-0 | 32.39 | 48A1-473-0 | 31.61 | x | 32.72 | 48A1-1 | x | 48A1-473-1 | x | | x | 48A1-2 | 31.45 | 48A1-473-2 | 32.90 | x | 34.44 | 48A1-3 | | 48A1-473-3 | | x | 21.86 | 48A1-4 | | 48A1-473-4 | | x | |
| | | 48A1-474-0 | 31.83 | | 33.04 | | | 48A1-474-1 | x | | x | | | 48A1-474-2 | 33.41 | | 35.41 | | | 48A1-474-3 | 23.87 | | 25.56 | | | 48A1-474-4 | | | |
| | | 48A1-475-0 | 31.87 | | 33.10 | | | 48A1-475-1 | x | | x | | | 48A1-475-2 | 33.00 | | 33.17 | | | 48A1-475-3 | 23.77 | | 26.40 | | | 48A1-475-4 | | | |
| | | | | | | | | 48A0-476-1 | x | | x | | | 48A0-476-2 | 29.98 | | 32.18 | | | 48A0-476-3 | | | | | | 48A0-476-4 | | | |
| | | | | | | | | 48A0-477-1 | x | | x | | | 48A0-477-2 | 31.98 | | 35.56 | | | 48A0-477-3 | | | 24.96 | | | 48A0-477-4 | | | |
| | | - | | | | 48A0-1 | x | 48A0-478-1 | 37.84 | | 37.98 | 48A0-2 | 33.11 | 48A0-478-2 | 35.48 | x | 38.62 | 48A0-3 | | 48A0-478-3 | 23.17 | x | 26.44 | 48A0-4 | | 48A0-478-4 | | x | |
| | | | | | | | | 48A0-479-1 | x | | x | | | 48A0-479-2 | 32.53 | | 34.96 | | | 48A0-479-3 | | | 22.15 | | | 48A0-479-4 | | | |
| | | | | | | | | 48A0-480-1 | x | | x | | | 48A0-480-2 | 30.18 | | 32.00 | | | 48A0-480-3 | | | | | | 48A0-480-4 | | | |
| | | 49A1-481-0 | 31.60 | | 32.91 | | | 49A1-481-1 | 37.80 | | 42.28 | | | 49A1-481-2 | | | | | | 49A1-481-3 | 30.64 | x | 33.64 | | | 49A1-481-4 | | | 23.20 |
| | | 49A1-482-0 | 31.48 | | 32.70 | | | 49A1-482-1 | x | | x | | | 49A1-482-2 | | | | | | 49A1-482-3 | 31.18 | x | 34.31 | | | 49A1-482-4 | | | 22.02 |
| 49A1-0 | 32.55 | 49A1-483-0 | 31.67 | x | 32.86 | 49A1-1 | x | 49A1-483-1 | x | | | 49A1-2 | x | ~~49A1-483-2~~ | | | | 49A1-3 | 25.69 | ~~49A1-483-3~~ | | | | 49A1-4 | | ~~49A1-483-4~~ | | x | |
| | | 49A1-484-0 | 31.10 | | 32.45 | | | 49A1-484-1 | x | | x | | | 49A1-484-2 | | | | | | 49A1-484-3 | 25.56 | x | 28.86 | | | 49A1-484-4 | | | |
| | | 49A1-485-0 | 32.17 | | 33.33 | | | 49A1-485-1 | x | | x | | | 49A1-485-2 | x | | | | | ~~49A1-485-3~~ | | | | | | ~~49A1-485-4~~ | | | |
| | | | | | | | | 49A0-486-1 | x | | x | | | 49A0-486-2 | x | | | | | 49A0-486-3 | x | | | | | 49A0-486-4 | 21.64 | | 24.89 |
| | | | | | | | | 49A0-487-1 | x | | x | | | 49A0-487-2 | 19.68 | x | 21.92 | | | 49A0-487-3 | | x | | | | 49A0-487-4 | | | |
| | | - | | | | 49A0-1 | x | 49A0-488-1 | x | | x | 49A0-2 | 23.99 | 49A0-488-2 | x | | | 49A0-3 | | 49A0-488-3 | 29.96 | x | 32.78 | 49A0-4 | | 49A0-488-4 | | x | |
| | | | | | | | | 49A0-489-1 | 36.98 | | 41.85 | | | 49A0-489-2 | 36.80 | x | x | | | 49A0-489-3 | 32.82 | x | 36.41 | | | 49A0-489-4 | | | 24.02 |
| | | | | | | | | 49A0-490-1 | x | | x | | | 49A0-490-2 | x | | | | | 49A0-490-3 | x | | | | | 49A0-490-4 | 20.00 | | 23.46 |

**T0**

| Pool ID | Ct (M) | Sample ID | Ct (M) | Ct (H5 HA) | Ct (H9 HA) |
|---|---|---|---|---|---|
| | | 50A1-491-0 | | | |
| | | 50A1-492-0 | | | |
| 50A1-0 | x | 50A1-493-0 | | | |
| | | 50A1-494-0 | | | |
| | | 50A1-495-0 | | | |
| | | - | | | |
| | | 51A1-501-0 | | | |
| | | 51A1-502-0 | | | |
| 51A1-0 | x | 51A1-503-0 | | | |
| | | 51A1-504-0 | | | |
| | | 51A1-505-0 | | | |
| | | - | | | |
| | | 52A1-511-0 | | | |
| | | 52A1-512-0 | | | |
| 52A1-0 | x | 52A1-513-0 | | | |
| | | 52A1-514-0 | | | |
| | | 52A1-515-0 | | | |
| | | - | | | |
| | | 53A1-521-0 | | | |
| | | 53A1-522-0 | | | |
| 53A1-0 | x | 53A1-523-0 | | | |
| | | 53A1-524-0 | | | |
| | | 53A1-525-0 | | | |
| | | - | | | |
| | | 54A1-531-0 | | | |
| | | 54A1-532-0 | | | |
| 54A1-0 | x | 54A1-533-0 | | | |
| | | 54A1-534-0 | | | |
| | | 54A1-535-0 | | | |
| | | - | | | |

**T1**

| Pool ID | Ct (M) | Sample ID | Ct (M) | Ct (H5 HA) | Ct (H9 HA) |
|---|---|---|---|---|---|
| | | 50A1-491-1 | | | |
| | | 50A1-492-1 | | | |
| 50A1-1 | x | 50A1-493-1 | | | |
| | | 50A1-494-1 | | | |
| | | 50A1-495-1 | | | |
| | | 50A0-496-1 | 23.48 | x | 21.38 |
| | | 50A0-497-1 | 42.68 | | |
| 50A0-1 | 24.92 | 50A0-498-1 | x | | |
| | | 50A0-499-1 | 37.83 | x | 39.70 |
| | | 50A0-500-1 | 36.36 | x | 42.32 |
| | | 51A1-501-1 | | | |
| | | 51A1-502-1 | | | |
| 51A1-1 | x | 51A1-503-1 | | | |
| | | 51A1-504-1 | | | |
| | | 51A1-505-1 | | | |
| | | 51A0-506-1 | x | | |
| | | 51A0-507-1 | 25.25 | x | 29.91 |
| 51A0-1 | 27.88 | 51A0-508-1 | 32.91 | x | 33.12 |
| | | 51A0-509-1 | 24.29 | x | 27.57 |
| | | 51A0-510-1 | x | | |
| | | 52A1-511-1 | | | |
| | | 52A1-512-1 | | | |
| 52A1-1 | x | 52A1-513-1 | | | |
| | | 52A1-514-1 | | | |
| | | 52A1-515-1 | | | |
| | | 52A0-516-1 | 21.60 | | 23.18 |
| | | 52A0-517-1 | 25.44 | | 24.00 |
| 52A0-1 | 23.61 | 52A0-518-1 | 21.50 | x | 20.20 |
| | | 52A0-519-1 | 24.28 | | 28.96 |
| | | 52A0-520-1 | 22.58 | | 24.54 |
| | | 53A1-521-1 | | | |
| | | 53A1-522-1 | | | |
| 53A1-1 | x | 53A1-523-1 | | | |
| | | 53A1-524-1 | | | |
| | | 53A1-525-1 | | | |
| | | 53A0-526-1 | | | |
| | | 53A0-527-1 | | | |
| 53A0-1 | x | 53A0-528-1 | | | |
| | | 53A0-529-1 | | | |
| | | 53A0-530-1 | | | |
| | | 54A1-531-1 | x | | |
| | | 54A1-532-1 | x | | |
| 54A1-1 | 36.45 | 54A1-533-1 | x | | |
| | | 54A1-534-1 | x | | |
| | | 54A1-535-1 | 41.79 | | |
| | | 54A0-536-1 | | | |
| | | 54A0-537-1 | | | |
| 54A0-1 | x | 54A0-538-1 | | | |
| | | 54A0-539-1 | | | |
| | | 54A0-540-1 | | | |

**T2**

| Pool ID | Ct (M) | Sample ID | Ct (M) | Ct (H5 HA) | Ct (H9 HA) |
|---|---|---|---|---|---|
| | | 50A1-491-2 | 27.99 | | 29.89 |
| | | 50A1-492-2 | 29.18 | | 31.49 |
| 50A1-2 | 29.09 | 50A1-493-2 | 28.93 | x | 31.14 |
| | | 50A1-494-2 | 30.59 | | 33.43 |
| | | 50A1-495-2 | 30.49 | | 31.91 |
| | | 50A0-496-2 | | | |
| | | 50A0-497-2 | 28.63 | | 31.30 |
| 50A0-2 | | 50A0-498-2 | 31.76 | x | 33.42 |
| | | 50A0-499-2 | 21.58 | | 23.96 |
| | | 50A0-500-2 | x | | x |
| | | 51A1-501-2 | 34.09 | | 33.30 |
| | | 51A1-502-2 | 34.64 | | 37.78 |
| 51A1-2 | 31.96 | 51A1-503-2 | 37.79 | x | x |
| | | 51A1-504-2 | 31.09 | | 32.34 |
| | | 51A1-505-2 | 28.62 | | 30.49 |
| | | 51A0-506-2 | 26.75 | | 26.49 |
| | | 51A0-507-2 | | | |
| 51A0-2 | | 51A0-508-2 | | x | 23.32 |
| | | 51A0-509-2 | | | |
| | | 51A0-510-2 | 31.18 | | 33.51 |
| | | 52A1-511-2 | 32.94 | | 36.41 |
| | | 52A1-512-2 | 30.95 | | 33.25 |
| 52A1-2 | 32.31 | 52A1-513-2 | 31.61 | x | 33.46 |
| | | 52A1-514-2 | 34.04 | | 35.39 |
| | | 52A1-515-2 | 33.13 | | 34.64 |
| | | 52A0-516-2 | | | |
| | | 52A0-517-2 | | | |
| 52A0-2 | | 52A0-518-2 | | x | |
| | | 52A0-519-2 | | | |
| | | 52A0-520-2 | | | |
| | | 53A1-521-2 | | | |
| | | 53A1-522-2 | | | |
| 53A1-2 | x | 53A1-523-2 | | | |
| | | 53A1-524-2 | | | |
| | | 53A1-525-2 | | | |
| | | 53A0-526-2 | | | |
| | | 53A0-527-2 | | | |
| 53A0-2 | x | 53A0-528-2 | | | |
| | | 53A0-529-2 | | | |
| | | 53A0-530-2 | | | |
| | | 54A1-531-2 | x | | |
| | | 54A1-532-2 | 27.15 | | 28.52 |
| 54A1-2 | 31.13 | 54A1-533-2 | 30.56 | x | 32.93 |
| | | 54A1-534-2 | 33.90 | | 37.92 |
| | | 54A1-535-2 | 35.20 | | 44.58 |
| | | 54A0-536-2 | x | | |
| | | 54A0-537-2 | 31.49 | | 33.84 |
| 54A0-2 | 36.97 | 54A0-538-2 | x | x | |
| | | 54A0-539-2 | 24.24 | | 26.35 |
| | | 54A0-540-2 | 35.47 | | 39.54 |

**T3**

| Pool ID | Ct (M) | Sample ID | Ct (M) | Ct (H5 HA) | Ct (H9 HA) |
|---|---|---|---|---|---|
| | | 50A1-491-3 | | | |
| | | 50A1-492-3 | | | |
| 50A1-3 | | 50A1-493-3 | | x | |
| | | 50A1-494-3 | | | 22.64 |
| | | 50A1-495-3 | | | |
| | | 50A0-496-3 | | | |
| | | 50A0-497-3 | | | |
| 50A0-3 | | 50A0-498-3 | | x | 23.70 |
| | | 50A0-499-3 | | | |
| | | 50A0-500-3 | 20.91 | | 23.57 |
| | | 51A1-501-3 | 22.17 | | 23.96 |
| | | 51A1-502-3 | 20.29 | | 24.33 |
| 51A1-3 | | 51A1-503-3 | 24.75 | x | 27.91 |
| | | 51A1-504-3 | | | |
| | | 51A1-505-3 | | | |
| | | 51A0-506-3 | | | |
| | | 51A0-507-3 | | | |
| 51A0-3 | | 51A0-508-3 | | x | |
| | | 51A0-509-3 | | | |
| | | 51A0-510-3 | | | 24.35 |
| | | 52A1-511-3 | | | 23.06 |
| | | 52A1-512-3 | | | 22.68 |
| 52A1-3 | | 52A1-513-3 | | x | 25.06 |
| | | 52A1-514-3 | 21.64 | | 24.92 |
| | | 52A1-515-3 | 20.93 | | 24.59 |
| | | 52A0-516-3 | | | |
| | | 52A0-517-3 | | | |
| 52A0-3 | | 52A0-518-3 | | x | |
| | | 52A0-519-3 | | | |
| | | 52A0-520-3 | | | |
| | | 53A1-521-3 | 20.70 | | 24.65 |
| | | 53A1-522-3 | 19.00 | | 21.39 |
| 53A1-3 | 20.73 | 53A1-523-3 | 20.93 | x | 24.39 |
| | | 53A1-524-3 | 20.80 | | 23.98 |
| | | 53A1-525-3 | 22.98 | | 26.05 |
| | | 53A0-526-3 | 22.36 | | 25.45 |
| | | 53A0-527-3 | 22.52 | | 25.40 |
| 53A0-3 | 22.35 | 53A0-528-3 | 22.80 | x | 25.98 |
| | | 53A0-529-3 | 22.34 | | 25.44 |
| | | 53A0-530-3 | 20.41 | | 23.56 |
| | | 54A1-531-3 | 20.18 | | 23.47 |
| | | 54A1-532-3 | | | |
| 54A1-3 | | 54A1-533-3 | | x | |
| | | 54A1-534-3 | 20.48 | | 23.68 |
| | | 54A1-535-3 | 19.98 | | 23.04 |
| | | 54A0-536-3 | 23.24 | | 26.62 |
| | | 54A0-537-3 | | | 24.50 |
| 54A0-3 | | 54A0-538-3 | 27.92 | x | 30.79 |
| | | 54A0-539-3 | | | |
| | | 54A0-540-3 | 21.00 | | 24.50 |

**T4**

| Pool ID | Ct (M) | Sample ID | Ct (M) | Ct (H5 HA) | Ct (H9 HA) |
|---|---|---|---|---|---|
| | | 50A1-491-4 | | | |
| | | 50A1-492-4 | | | |
| 50A1-4 | | 50A1-493-4 | | x | |
| | | 50A1-494-4 | | | |
| | | 50A1-495-4 | | | |
| | | 50A0-496-4 | 22.61 | | |
| | | 50A0-497-4 | | | |
| 50A0-4 | | 50A0-498-4 | | x | |
| | | 50A0-499-4 | | | |
| | | 50A0-500-4 | | | |
| | | 51A1-501-4 | | | |
| | | 51A1-502-4 | | | |
| 51A1-4 | | 51A1-503-4 | | x | |
| | | 51A1-504-4 | | | |
| | | 51A1-505-4 | | | |
| | | 51A0-506-4 | | | |
| | | 51A0-507-4 | | | |
| 51A0-4 | | 51A0-508-4 | | x | |
| | | 51A0-509-4 | | | |
| | | 51A0-510-4 | | | |
| | | 52A1-511-4 | | | |
| | | 52A1-512-4 | | | |
| 52A1-4 | | 52A1-513-4 | | x | |
| | | 52A1-514-4 | | | |
| | | 52A1-515-4 | | | |
| | | 52A0-516-4 | | | |
| | | 52A0-517-4 | | | |
| 52A0-4 | | 52A0-518-4 | | x | |
| | | 52A0-519-4 | | | |
| | | 52A0-520-4 | | | |
| | | 53A1-521-4 | | | |
| | | 53A1-522-4 | | | |
| 53A1-4 | | 53A1-523-4 | | x | |
| | | 53A1-524-4 | | | |
| | | 53A1-525-4 | 23.35 | | |
| | | 53A0-526-4 | | | |
| | | 53A0-527-4 | | | |
| 53A0-4 | | 53A0-528-4 | | x | |
| | | 53A0-529-4 | | | |
| | | 53A0-530-4 | | | |
| | | 54A1-531-4 | | | |
| | | 54A1-532-4 | | | |
| 54A1-4 | | 54A1-533-4 | | x | |
| | | 54A1-534-4 | | | |
| | | 54A1-535-4 | | | |
| | | 54A0-536-4 | | | |
| | | 54A0-537-4 | | | |
| 54A0-4 | | 54A0-538-4 | | x | |
| | | 54A0-539-4 | | | |
| | | 54A0-540-4 | 19.80 | | |

**T0**

| Pool ID | Ct (M) | Sample ID | Ct (M) | Ct (H5 HA) | Ct (H9 HA) |
|---|---|---|---|---|---|
| | | 55A1-541-0 | | | |
| | | 55A1-542-0 | | | |
| 55A1-0 | x | 55A1-543-0 | | | |
| | | 55A1-544-0 | | | |
| | | 55A1-545-0 | | | |
| | | - | | | |
| | | 56A1-551-0 | | | |
| | | 56A1-552-0 | | | |
| 56A1-0 | x | 56A1-553-0 | | | |
| | | 56A1-554-0 | | | |
| | | 56A1-555-0 | | | |
| | | - | | | |
| | | 57A1-561-0 | | | |
| | | 57A1-562-0 | | | |
| 57A1-0 | x | 57A1-563-0 | | | |
| | | 57A1-564-0 | | | |
| | | 57A1-565-0 | | | |
| | | - | | | |
| | | 58A1-571-0 | | | |
| | | 58A1-572-0 | | | |
| 58A1-0 | x | 58A1-573-0 | | | |
| | | 58A1-574-0 | | | |
| | | 58A1-575-0 | | | |
| | | - | | | |
| | | 59A1-581-0 | | | |
| | | 59A1-582-0 | | | |
| 59A1-0 | x | 59A1-583-0 | | | |
| | | 59A1-584-0 | | | |
| | | 59A1-585-0 | | | |
| | | - | | | |

**T1**

| Pool ID | Ct (M) | Sample ID | Ct (M) | Ct (H5 HA) | Ct (H9 HA) |
|---|---|---|---|---|---|
| | | 55A1-541-1 | | | |
| | | 55A1-542-1 | | | |
| 55A1-1 | | 55A1-543-1 | | | |
| | | 55A1-544-1 | | | |
| | | 55A1-545-1 | | | |
| | | 55A0-546-1 | | | |
| | | 55A0-547-1 | | | |
| 55A0-1 | | 55A0-548-1 | | | |
| | | 55A0-549-1 | | | |
| | | 55A0-550-1 | | | |
| | | 56A1-551-1 | | | |
| | | 56A1-552-1 | | | |
| 56A1-1 | x | 56A1-553-1 | | | |
| | | 56A1-554-1 | | | |
| | | 56A1-555-1 | | | |
| | | 56A0-556-1 | | | |
| | | 56A0-557-1 | | | |
| 56A0-1 | x | 56A0-558-1 | | | |
| | | 56A0-559-1 | | | |
| | | 56A0-560-1 | | | |
| | | 57A1-561-1 | | | |
| | | 57A1-562-1 | | | |
| 57A1-1 | x | 57A1-563-1 | | | |
| | | 57A1-564-1 | | | |
| | | 57A1-565-1 | | | |
| | | 57A0-566-1 | | | |
| | | 57A0-567-1 | | | |
| 57A0-1 | x | 57A0-568-1 | | | |
| | | 57A0-569-1 | | | |
| | | 57A0-570-1 | | | |
| | | 58A1-571-1 | 43.24 | | |
| | | 58A1-572-1 | 24.88 | x | 24.02 |
| 58A1-1 | 28.59 | 58A1-573-1 | 35.24 | x | x |
| | | 58A1-574-1 | 39.44 | x | x |
| | | 58A1-575-1 | x | | |
| | | 58A0-576-1 | | | |
| | | 58A0-577-1 | | | |
| 58A0-1 | x | 58A0-578-1 | | | |
| | | 58A0-579-1 | | | |
| | | 58A0-580-1 | | | |
| | | 59A1-581-1 | | | |
| | | 59A1-582-1 | | | |
| 59A1-1 | x | 59A1-583-1 | | | |
| | | 59A1-584-1 | | | |
| | | 59A1-585-1 | | | |
| | | 59A0-586-1 | | | |
| | | 59A0-587-1 | | | |
| 59A0-1 | x | 59A0-588-1 | | | |
| | | 59A0-589-1 | | | |
| | | 59A0-590-1 | | | |

**T2**

| Pool ID | Ct (M) | Sample ID | Ct (M) | Ct (H5 HA) | Ct (H9 HA) |
|---|---|---|---|---|---|
| | | 55A1-541-2 | | | |
| | | 55A1-542-2 | | | |
| 55A1-2 | | 55A1-543-2 | | | |
| | | 55A1-544-2 | | | |
| | | 55A1-545-2 | | | |
| | | 55A0-546-2 | | | |
| | | 55A0-547-2 | | | |
| 55A0-2 | | 55A0-548-2 | | | |
| | | 55A0-549-2 | | | |
| | | 55A0-550-2 | | | |
| | | 56A1-551-2 | | | |
| | | 56A1-552-2 | | | |
| 56A1-2 | x | 56A1-553-2 | | | |
| | | 56A1-554-2 | | | |
| | | 56A1-555-2 | | | |
| | | 56A0-556-2 | 26.97 | | 29.10 |
| | | 56A0-557-2 | 30.68 | | 32.66 |
| 56A0-2 | 24.65 | 56A0-558-2 | 21.21 | x | 23.20 |
| | | 56A0-559-2 | 23.71 | | 25.90 |
| | | 56A0-560-2 | 24.80 | | 26.15 |
| | | 57A1-561-2 | x | | |
| | | 57A1-562-2 | 33.77 | | 35.99 |
| 57A1-2 | 30.26 | 57A1-563-2 | 31.94 | x | 34.09 |
| | | 57A1-564-2 | 28.70 | | 31.14 |
| | | 57A1-565-2 | 31.13 | | 31.00 |
| | | 57A0-566-2 | 26.42 | | 27.95 |
| | | 57A0-567-2 | 28.54 | | 30.05 |
| 57A0-2 | 33.24 | 57A0-568-2 | 32.16 | x | x |
| | | 57A0-569-2 | 29.83 | | 32.53 |
| | | 57A0-570-2 | 30.81 | | 31.13 |
| | | 58A1-571-2 | x | | |
| | | 58A1-572-2 | | | |
| 58A1-2 | | 58A1-573-2 | 32.29 | x | 33.73 |
| | | 58A1-574-2 | 28.47 | | x |
| | | 58A1-575-2 | 34.22 | | 38.72 |
| | | 58A0-576-2 | 28.87 | | 29.62 |
| | | 58A0-577-2 | x | | |
| 58A0-2 | 32.60 | 58A0-578-2 | 33.44 | x | 34.38 |
| | | 58A0-579-2 | 32.35 | | 35.13 |
| | | 58A0-580-2 | 35.44 | | 40.10 |
| | | 59A1-581-2 | 23.66 | | 24.91 |
| | | 59A1-582-2 | 36.03 | | 35.21 |
| 59A1-2 | 26.34 | 59A1-583-2 | x | x | |
| | | 59A1-584-2 | 33.88 | | 37.95 |
| | | 59A1-585-2 | 25.69 | | 26.67 |
| | | 59A0-586-2 | 23.93 | | 25.26 |
| | | 59A0-587-2 | 29.91 | | 30.85 |
| 59A0-2 | 24.86 | 59A0-588-2 | 25.06 | x | 26.35 |
| | | 59A0-589-2 | 24.32 | | 25.87 |
| | | 59A0-590-2 | 21.06 | | 22.74 |

**T3**

| Pool ID | Ct (M) | Sample ID | Ct (M) | Ct (H5 HA) | Ct (H9 HA) |
|---|---|---|---|---|---|
| | | 55A1-541-3 | | | |
| | | 55A1-542-3 | | | |
| 55A1-3 | | 55A1-543-3 | | | |
| | | 55A1-544-3 | | | |
| | | 55A1-545-3 | | | |
| | | 55A0-546-3 | | | |
| | | 55A0-547-3 | | | |
| 55A0-3 | | 55A0-548-3 | | | |
| | | 55A0-549-3 | | | |
| | | 55A0-550-3 | | | |
| | | 56A1-551-3 | 22.24 | | 25.80 |
| | | 56A1-552-3 | 20.63 | | 23.61 |
| 56A1-3 | 20.86 | 56A1-553-3 | 18.54 | x | 21.45 |
| | | 56A1-554-3 | 21.12 | | 23.80 |
| | | 56A1-555-3 | 24.34 | | 26.62 |
| | | 56A0-556-3 | | | |
| | | 56A0-557-3 | | | |
| 56A0-3 | | 56A0-558-3 | | x | |
| | | 56A0-559-3 | | | |
| | | 56A0-560-3 | | | |
| | | 57A1-561-3 | 24.25 | | 26.58 |
| | | 57A1-562-3 | 24.82 | | 27.90 |
| 57A1-3 | | 57A1-563-3 | | x | 26.67 |
| | | 57A1-564-3 | | | |
| | | 57A1-565-3 | | | |
| | | 57A0-566-3 | | | |
| | | 57A0-567-3 | | | |
| 57A0-3 | | 57A0-568-3 | | x | 22.76 |
| | | 57A0-569-3 | | | |
| | | 57A0-570-3 | | | |
| | | 58A1-571-3 | 33.09 | | 36.54 |
| | | 58A1-572-3 | | | |
| 58A1-3 | | 58A1-573-3 | | x | 21.98 |
| | | 58A1-574-3 | | | 22.34 |
| | | 58A1-575-3 | 22.16 | | 23.35 |
| | | 58A0-576-3 | | | |
| | | 58A0-577-3 | 22.19 | | 25.21 |
| 58A0-3 | | 58A0-578-3 | 20.03 | x | 22.73 |
| | | 58A0-579-3 | | | 23.00 |
| | | 58A0-580-3 | 22.94 | | 25.79 |
| | | 59A1-581-3 | | | |
| | | 59A1-582-3 | 23.65 | | 24.22 |
| 59A1-3 | | 59A1-583-3 | 31.14 | x | 33.52 |
| | | 59A1-584-3 | 23.00 | | 25.10 |
| | | 59A1-585-3 | | | |
| | | 59A0-586-3 | | | |
| | | 59A0-587-3 | | | |
| 59A0-3 | | 59A0-588-3 | | x | |
| | | 59A0-589-3 | | | |
| | | 59A0-590-3 | | | |

**T4**

| Pool ID | Ct (M) | Sample ID | Ct (M) | Ct (H5 HA) | Ct (H9 HA) |
|---|---|---|---|---|---|
| | | 55A1-541-4 | | | |
| | | 55A1-542-4 | | | |
| 55A1-4 | | 55A1-543-4 | | | |
| | | 55A1-544-4 | | | |
| | | 55A1-545-4 | | | |
| | | 55A0-546-4 | | | |
| | | 55A0-547-4 | | | |
| 55A0-4 | | 55A0-548-4 | | | |
| | | 55A0-549-4 | | | |
| | | 55A0-550-4 | | | |
| | | 56A1-551-4 | | | |
| | | 56A1-552-4 | | | |
| 56A1-4 | | 56A1-553-4 | | x | |
| | | 56A1-554-4 | | | |
| | | 56A1-555-4 | | | |
| | | 56A0-556-4 | | | |
| | | 56A0-557-4 | | | |
| 56A0-4 | | 56A0-558-4 | | x | |
| | | 56A0-559-4 | | | |
| | | 56A0-560-4 | | | |
| | | 57A1-561-4 | | | |
| | | 57A1-562-4 | | | |
| 57A1-4 | | 57A1-563-4 | | x | |
| | | 57A1-564-4 | | | |
| | | 57A1-565-4 | | | |
| | | 57A0-566-4 | | | |
| | | 57A0-567-4 | | | |
| 57A0-4 | | 57A0-568-4 | | x | |
| | | 57A0-569-4 | | | |
| | | 57A0-570-4 | | | |
| | | 58A1-571-4 | 30.84 | | 33.61 |
| | | 58A1-572-4 | | | |
| 58A1-4 | | 58A1-573-4 | | x | |
| | | 58A1-574-4 | | | |
| | | 58A1-575-4 | | | |
| | | 58A0-576-4 | | | |
| | | 58A0-577-4 | | | |
| 58A0-4 | | 58A0-578-4 | | x | |
| | | 58A0-579-4 | | | |
| | | 58A0-580-4 | | | |
| | | 59A1-581-4 | | | |
| | | 59A1-582-4 | | | |
| 59A1-4 | | 59A1-583-4 | | x | 23.00 |
| | | 59A1-584-4 | 19.95 | | |
| | | 59A1-585-4 | 22.32 | | |
| | | 59A0-586-4 | | | |
| | | 59A0-587-4 | | | |
| 59A0-4 | | 59A0-588-4 | | x | |
| | | 59A0-589-4 | | | |
| | | 59A0-590-4 | | | |

| T0 | | | | | | T1 | | | | | | T2 | | | | | | T3 | | | | | | T4 | | | | | |
|---|---|---|---|---|---|---|---|---|---|---|---|---|---|---|---|---|---|---|---|---|---|---|---|---|---|---|---|---|---|
| Pool ID | Ct (M) | Sample ID | Ct (M) | Ct (H5 HA) | Ct (H9 HA) | Pool ID | Ct (M) | Sample ID | Ct (M) | Ct (H5 HA) | Ct (H9 HA) | Pool ID | Ct (M) | Sample ID | Ct (M) | Ct (H5 HA) | Ct (H9 HA) | Pool ID | Ct (M) | Sample ID | Ct (M) | Ct (H5 HA) | Ct (H9 HA) | Pool ID | Ct (M) | Sample ID | Ct (M) | Ct (H5 HA) | Ct (H9 HA) |
| | | 60A1-591-0 | | | | | | 60A1-591-1 | | | | | | 60A1-591-2 | 37.59 | | x | | | 60A1-591-3 | 20.34 | | 23.90 | | | 60A1-591-4 | | | |
| | | 60A1-592-0 | | | | | | 60A1-592-1 | | | | | | 60A1-592-2 | 30.93 | | x | | | 60A1-592-3 | | | 24.10 | | | 60A1-592-4 | | | |
| 60A1-0 | x | 60A1-593-0 | | | | 60A1-1 | x | 60A1-593-1 | | | | 60A1-2 | 36.20 | 60A1-593-2 | 30.96 | x | 32.41 | 60A1-3 | | 60A1-593-3 | | x | | 60A1-4 | | 60A1-593-4 | | x | |
| | | 60A1-594-0 | | | | | | 60A1-594-1 | | | | | | 60A1-594-2 | 30.71 | | 32.24 | | | 60A1-594-3 | | | | | | 60A1-594-4 | | | |
| | | 60A1-595-0 | | | | | | 60A1-595-1 | | | | | | 60A1-595-2 | 29.12 | | 30.12 | | | 60A1-595-3 | | | | | | 60A1-595-4 | | | |
| | | | | | | | | 60A0-596-1 | | | | | | 60A0-596-2 | 24.70 | | 26.19 | | | 60A0-596-3 | | | | | | 60A0-596-4 | | | |
| | | | | | | | | 60A0-597-1 | | | | | | 60A0-597-2 | 21.12 | | 21.80 | | | 60A0-597-3 | | | | | | 60A0-597-4 | | | |
| | | - | | | | 60A0-1 | x | 60A0-598-1 | | | | 60A0-2 | 19.99 | 60A0-598-2 | 19.81 | x | 21.81 | 60A0-3 | | 60A0-598-3 | | x | | 60A0-4 | | 60A0-598-4 | | x | |
| | | | | | | | | 60A0-599-1 | | | | | | 60A0-599-2 | 21.39 | | 23.62 | | | 60A0-599-3 | | | | | | 60A0-599-4 | | | |
| | | | | | | | | 60A0-600-1 | | | | | | 60A0-600-2 | 23.08 | | 24.59 | | | 60A0-600-3 | | | | | | 60A0-600-4 | | | |
| | | 61A1-601-0 | | | | | | 61A1-601-1 | | | | | | 61A1-601-2 | x | | | | | 61A1-601-3 | 22.16 | | 24.57 | | | 61A1-601-4 | | | |
| | | 61A1-602-0 | | | | | | 61A1-602-1 | | | | | | 61A1-602-2 | 30.44 | | 32.60 | | | 61A1-602-3 | | | | | | 61A1-602-4 | | | |
| 61A1-0 | x | 61A1-603-0 | | | | 61A1-1 | x | 61A1-603-1 | | | | 61A1-2 | 32.39 | 61A1-603-2 | 30.22 | x | 30.62 | 61A1-3 | | 61A1-603-3 | | x | | 61A1-4 | | 61A1-603-4 | | x | |
| | | 61A1-604-0 | | | | | | 61A1-604-1 | | | | | | 61A1-604-2 | 30.95 | | 32.48 | | | 61A1-604-3 | | | | | | 61A1-604-4 | | | |
| | | 61A1-605-0 | | | | | | 61A1-605-1 | | | | | | 61A1-605-2 | x | | | | | 61A1-605-3 | 21.85 | | 25.22 | | | 61A1-605-4 | | | |
| | | | | | | | | 61A0-606-1 | | | | | | 61A0-606-2 | 20.82 | | 22.96 | | | 61A0-606-3 | | | | | | 61A0-606-4 | | | |
| | | | | | | | | 61A0-607-1 | | | | | | 61A0-607-2 | 23.36 | | 25.33 | | | 61A0-607-3 | | | | | | 61A0-607-4 | 21.68 | | |
| | | - | | | | 61A0-1 | x | 61A0-608-1 | | | | 61A0-2 | 22.80 | 61A0-608-2 | 31.98 | x | 34.49 | 61A0-3 | | 61A0-608-3 | | x | 24.15 | 61A0-4 | | 61A0-608-4 | | x | |
| | | | | | | | | 61A0-609-1 | | | | | | 61A0-609-2 | 24.25 | | 26.53 | | | 61A0-609-3 | | | | | | 61A0-609-4 | | | |
| | | | | | | | | 61A0-610-1 | | | | | | 61A0-610-2 | 26.76 | | 28.56 | | | 61A0-610-3 | | | | | | 61A0-610-4 | | | |
| | | 62A1-611-0 | | | | | | 62A1-611-1 | | | | | | 62A1-611-2 | 20.99 | | 22.67 | | | 62A1-611-3 | | | | | | 62A1-611-4 | | | |
| | | 62A1-612-0 | | | | | | 62A1-612-1 | | | | | | 62A1-612-2 | 21.56 | | 23.26 | | | 62A1-612-3 | | | | | | 62A1-612-4 | | | |
| 62A1-0 | x | 62A1-613-0 | | | | 62A1-1 | x | 62A1-613-1 | | | | 62A1-2 | 23.33 | 62A1-613-2 | 21.93 | x | 24.10 | 62A1-3 | | 62A1-613-3 | | x | | 62A1-4 | | 62A1-613-4 | | x | |
| | | 62A1-614-0 | | | | | | 62A1-614-1 | | | | | | 62A1-614-2 | 20.62 | | 22.68 | | | 62A1-614-3 | | | | | | 62A1-614-4 | | | |
| | | 62A1-615-0 | | | | | | 62A1-615-1 | | | | | | 62A1-615-2 | 22.41 | | 24.28 | | | 62A1-615-3 | | | | | | 62A1-615-4 | | | |
| | | | | | | | | 62A0-616-1 | | | | | | 62A0-616-2 | 28.63 | | 29.97 | | | 62A0-616-3 | | | | | | 62A0-616-4 | | | |
| | | | | | | | | 62A0-617-1 | | | | | | 62A0-617-2 | 19.52 | | 20.68 | | | 62A0-617-3 | | | | | | 62A0-617-4 | | | |
| | | - | | | | 62A0-1 | x | 62A0-618-1 | | | | 62A0-2 | 24.43 | 62A0-618-2 | 25.81 | x | 27.72 | 62A0-3 | | 62A0-618-3 | | x | | 62A0-4 | | 62A0-618-4 | | x | |
| | | | | | | | | 62A0-619-1 | | | | | | 62A0-619-2 | 24.23 | | 23.77 | | | 62A0-619-3 | | | | | | 62A0-619-4 | | | |
| | | | | | | | | 62A0-620-1 | | | | | | 62A0-620-2 | 29.81 | | 31.03 | | | 62A0-620-3 | | | | | | 62A0-620-4 | | | |
| | | 63A1-621-0 | | | | | | 63A1-621-1 | | | | | | 63A1-621-2 | 31.28 | | 32.71 | | | 63A1-621-3 | | | | | | 63A1-621-4 | | | |
| | | 63A1-622-0 | | | | | | 63A1-622-1 | | | | | | 63A1-622-2 | 30.85 | | 31.62 | | | 63A1-622-3 | | | | | | 63A1-622-4 | | | |
| 63A1-0 | x | 63A1-623-0 | | | | 63A1-1 | x | 63A1-623-1 | | | | 63A1-2 | 31.93 | 63A1-623-2 | 33.19 | x | 36.01 | 63A1-3 | | 63A1-623-3 | 25.67 | x | 28.12 | 63A1-4 | | 63A1-623-4 | | x | |
| | | 63A1-624-0 | | | | | | 63A1-624-1 | | | | | | 63A1-624-2 | 28.27 | | 28.19 | | | 63A1-624-3 | | | | | | 63A1-624-4 | | | |
| | | 63A1-625-0 | | | | | | 63A1-625-1 | | | | | | 63A1-625-2 | 36.99 | | x | | | 63A1-625-3 | 25.74 | | 29.02 | | | 63A1-625-4 | | | |
| | | | | | | | | 63A0-626-1 | | | | | | 63A0-626-2 | 35.54 | | 37.39 | | | 63A0-626-3 | 24.47 | | 26.55 | | | 63A0-626-4 | | | |
| | | | | | | | | 63A0-627-1 | | | | | | 63A0-627-2 | x | | | | | 63A0-627-3 | 23.62 | | 26.10 | | | 63A0-627-4 | | | |
| | | - | | | | 63A0-1 | x | 63A0-628-1 | | | | 63A0-2 | 30.05 | 63A0-628-2 | 35.88 | x | 42.40 | 63A0-3 | | 63A0-628-3 | 25.76 | x | 27.53 | 63A0-4 | | 63A0-628-4 | | x | |
| | | | | | | | | 63A0-629-1 | | | | | | 63A0-629-2 | 34.02 | | 34.13 | | | 63A0-629-3 | 23.33 | | 23.38 | | | 63A0-629-4 | | | |
| | | | | | | | | 63A0-630-1 | | | | | | 63A0-630-2 | 28.04 | | 28.12 | | | 63A0-630-3 | | | | | | 63A0-630-4 | | | |
| | | 64A1-631-0 | | | | | | 64A1-631-1 | | | | | | 64A1-631-2 | 37.17 | | 37.67 | | | 64A1-631-3 | 34.42 | | 34.88 | | | 64A1-631-4 | 25.45 | | 28.50 |
| | | 64A1-632-0 | | | | | | 64A1-632-1 | | | | | | 64A1-632-2 | 34.30 | | 33.41 | | | 64A1-632-3 | 24.37 | | 27.03 | | | 64A1-632-4 | | | |
| 64A1-0 | x | 64A1-633-0 | | | | 64A1-1 | x | 64A1-633-1 | | | | 64A1-2 | 36.04 | 64A1-633-2 | 33.21 | x | 33.91 | 64A1-3 | | 64A1-633-3 | 27.91 | x | 30.85 | 64A1-4 | | 64A1-633-4 | | x | |
| | | 64A1-634-0 | | | | | | 64A1-634-1 | | | | | | 64A1-634-2 | 38.52 | | x | | | 64A1-634-3 | 26.23 | | 38.40 | | | 64A1-634-4 | | | 21.86 |
| | | 64A1-635-0 | | | | | | 64A1-635-1 | | | | | | 64A1-635-2 | 36.20 | | 42.50 | | | 64A1-635-3 | 27.80 | | 29.96 | | | 64A1-635-4 | | | |
| | | | | | | | | 64A0-636-1 | | | | | | 64A0-636-2 | 33.83 | | 44.07 | | | 64A0-636-3 | 26.57 | | 28.93 | | | 64A0-636-4 | | | |
| | | | | | | | | 64A0-637-1 | | | | | | 64A0-637-2 | 30.76 | | 31.90 | | | 64A0-637-3 | | | | | | 64A0-637-4 | | | |
| | | - | | | | 64A0-1 | x | 64A0-638-1 | | | | 64A0-2 | 32.76 | 64A0-638-2 | 32.72 | x | 33.18 | 64A0-3 | | 64A0-638-3 | | x | 29.04 | 64A0-4 | | 64A0-638-4 | | x | |
| | | | | | | | | 64A0-639-1 | | | | | | 64A0-639-2 | x | | | | | 64A0-639-3 | 24.88 | | 25.79 | | | 64A0-639-4 | 19.74 | | |
| | | | | | | | | 64A0-640-1 | | | | | | 64A0-640-2 | 31.77 | | 31.69 | | | 64A0-640-3 | | | | | | 64A0-640-4 | | | |

| T0 | | | | | |
|---|---|---|---|---|---|
| Pool ID | Ct (M) | Sample ID | Ct (M) | Ct (H5 HA) | Ct (H9 HA) |
| | | 65A1-641-0 | | | |
| | | 65A1-642-0 | | | |
| 65A1-0 | x | 65A1-643-0 | | | |
| | | 65A1-644-0 | | | |
| | | 65A1-645-0 | | | |
| | | - | | | |

| T1 | | | | | |
|---|---|---|---|---|---|
| Pool ID | Ct (M) | Sample ID | Ct (M) | Ct (H5 HA) | Ct (H9 HA) |
| | | 65A1-641-1 | | | |
| | | 65A1-642-1 | | | |
| 65A1-1 | x | 65A1-643-1 | | | |
| | | 65A1-644-1 | | | |
| | | 65A1-645-1 | | | |
| | | 65A0-646-1 | | | |
| | | 65A0-647-1 | | | |
| 65A0-1 | x | 65A0-648-1 | | | |
| | | 65A0-649-1 | | | |
| | | 65A0-650-1 | | | |

| T2 | | | | | |
|---|---|---|---|---|---|
| Pool ID | Ct (M) | Sample ID | Ct (M) | Ct (H5 HA) | Ct (H9 HA) |
| | | 65A1-641-2 | | | |
| | | 65A1-642-2 | | | |
| 65A1-2 | x | 65A1-643-2 | | | |
| | | 65A1-644-2 | | | |
| | | 65A1-645-2 | | | |
| | | 65A0-646-2 | x | | |
| | | 65A0-647-2 | 36.70 | | x |
| 65A0-2 | 33.93 | 65A0-648-2 | 31.14 | x | 28.83 |
| | | 65A0-649-2 | 29.54 | | 28.87 |
| | | 65A0-650-2 | 34.01 | | 42.43 |

| T3 | | | | | |
|---|---|---|---|---|---|
| Pool ID | Ct (M) | Sample ID | Ct (M) | Ct (H5 HA) | Ct (H9 HA) |
| | | 65A1-641-3 | 23.34 | | 24.74 |
| | | 65A1-642-3 | 24.41 | | 24.05 |
| 65A1-3 | 24.46 | 65A1-643-3 | 24.36 | x | 27.90 |
| | | 65A1-644-3 | 26.26 | | 28.84 |
| | | 65A1-645-3 | 24.55 | | 27.33 |
| | | 65A0-646-3 | 26.64 | | 26.74 |
| | | 65A0-647-3 | 21.46 | | 24.21 |
| 65A0-3 | | 65A0-648-3 | | x | |
| | | 65A0-649-3 | | | |
| | | 65A0-650-3 | 25.18 | | 27.44 |

| T4 | | | | | |
|---|---|---|---|---|---|
| Pool ID | Ct (M) | Sample ID | Ct (M) | Ct (H5 HA) | Ct (H9 HA) |
| | | 65A1-641-4 | | | |
| | | 65A1-642-4 | | | |
| 65A1-4 | | 65A1-643-4 | | x | |
| | | 65A1-644-4 | | | |
| | | 65A1-645-4 | | | |
| | | 65A0-646-4 | | | |
| | | 65A0-647-4 | | | |
| 65A0-4 | | 65A0-648-4 | | x | |
| | | 65A0-649-4 | | | |
| | | 65A0-650-4 | | | 22.14 |

Pool/Sample tested

Sample assumed positive

Pool/Sample re-extracted and re-tested

Sample re-tested

Sample taken from dead bird

Bird dropped out (e.g. death)

A, broiler; B, backyard chicken; Ct, cycle threshold; M, matrix; HA, haemagglutinin; T, time point; x, negative.

**Appendix Figure 3.** Detailed diagnostic test results for broilers and backyard chickens for all time points throughout the field experiment (T0–T4).

### Appendix H: Other figures

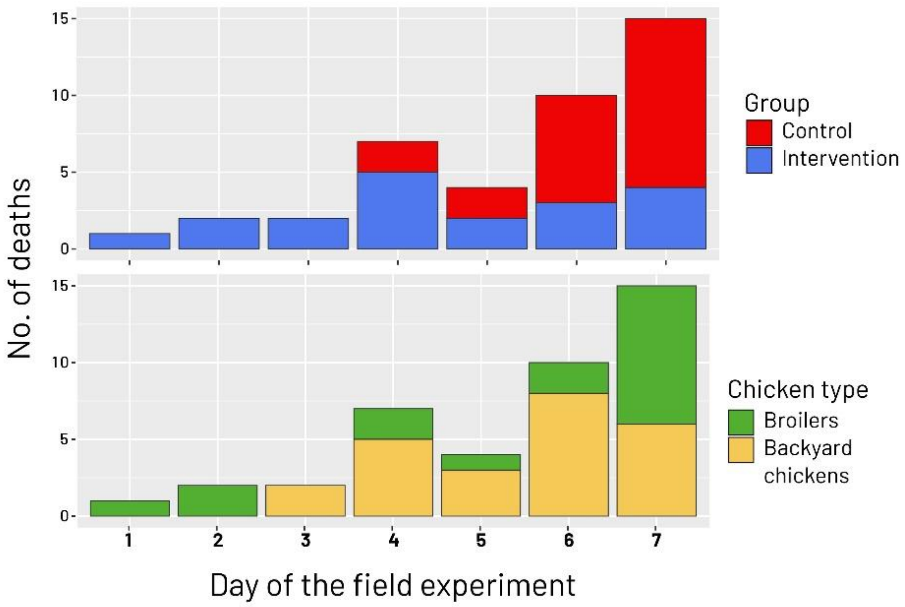

**Appendix Figure 4.** Number of chickens dropping out of the study due to sudden death.
